# A Community-Driven Single-Cell PBMC Reference Integrating Landmark Datasets Spanning Health and Disease

**DOI:** 10.64898/2026.09.21.749940

**Authors:** Ana-Maria Cujba, Sergio Aguilar-Fernandez, Adrien Antoinette, Wamia Said, Michaela F. Mueller, Kian Hong Kock, Jacquelyn Nestor, Radhika Sonthalia, Eliora V. Buyamin, Pragya Rawat, Alexander Predeus, Lorenz Kretschmer, Lijiang Fei, Kamil Slowikowski, Pritha Sen, Christopher V. Cosgriff, MGH COVID-19 Collection & Processing Team, Olli Dufva, Mohamad Askari, Kyle Kimler, Mary Futey, Ida Zucchi, Arsenios Chatzigeorgiou, Parisa Nejad, Liying Jin, Blake Bowen, Andrian Yang, Rik G. H. Lindeboom, Rachelly Normand, Stathis Megas, Yoshinari Ando, Ankita Chatterjee, Jong-Eun Park, Partha P. Majumder, Ponpan Matangkasombut, Varodom Charoensawan, Jay W. Shin, Woong-Yang Park, Asian Immune Diversity Atlas (AIDA) Network, Stephen Sansom, Berthold Göttgens, Joseph E. Powell, Lloyd Bod, Holger Heyn, Fabian J. Theis, Sarah A. Teichmann, Malte D. Luecken, Shyam Prabhakar, Alexandra-Chloé Villani, Gary Reynolds

**Affiliations:** Cambridge Stem Cell Institute, University of Cambridge, Cambridge, UK; Center for Immunology and Inflammatory Diseases, Department of Medicine, Massachusetts General Hospital, Boston, MA, USA; Krantz Family Center for Cancer Research, Department of Medicine, Massachusetts General Hospital, Boston, MA, USA; Broad Institute of MIT and Harvard, Cambridge, MA, USA; Harvard Medical School, Boston, MA, USA; Institute of Computational Biology, Helmholtz Munich, Neuherberg/Munich, Germany; TUM School of Life Sciences Weihenstephan, Technical University of Munich, Munich, Germany; School of Computation, Information and Technology, Technical University of Munich, Munich, Germany; Genome Institute of Singapore (GIS), Agency for Science, Technology and Research (A*STAR), Republic of Singapore; Wellcome Sanger Institute, Cambridge, UK; Sapienza University of Rome, Rome, Italy; Transplant, Oncology, and Immunocompromised Host Group, Division of Infectious Diseases, Brigham and Women’s Hospital, Boston, MA, USA; Cell Discovery Network, Boston Children’s Hospital, Boston, MA, USA; Department of Biomedical Informatics, Harvard Medical School, Boston, MA, USA; European Bioinformatics Institute (EMBL-EBI), Cambridge, UK; UC Santa Cruz Genomics Institute, Santa Cruz, CA, USA; Kennedy Institute of Rheumatology, University of Oxford, Oxford, UK; Garvan Institute for Medical Research, Darlinghurst, Australia; Cellular Genomics Futures Institute, University of New South Wales, Sydney, Australia; The Netherlands Cancer Institute, Amsterdam, The Netherlands; Medical University of Vienna, Vienna, Austria; RIKEN Center for Integrative Medical Sciences, Yokohama, Kanagawa, Japan; John C. Martin Centre for Liver Research and Innovations, Sonarpur, Kolkata, India; Graduate School of Medical Science and Engineering, Korea Advanced Institute of Science and Technology (KAIST), Republic of Korea; Indian Statistical Institute, Kolkata, India; Single-cell Omics and Systems Biology of Diseases Research Unit, Faculty of Science, Mahidol University, Bangkok, Thailand; Department of Microbiology, Faculty of Science, Mahidol University, Bangkok, Thailand; Siriraj Genomics, Faculty of Medicine Siriraj Hospital, Mahidol University, Bangkok, Thailand; Department of Biochemistry, Faculty of Medicine Siriraj Hospital, Mahidol University, Bangkok, Thailand; Integrative Computational BioScience (ICBS) center, Mahidol University, Nakhon Pathom, Thailand; Department of Biochemistry, Faculty of Science, Mahidol University, Bangkok, Thailand; Laboratory for Advanced Genomics Circuit, RIKEN Center for Integrative Medical Sciences, Yokohama, Kanagawa, Japan; Department of Biochemistry, Yong Loo Lin School of Medicine, National University of Singapore, Republic of Singapore; Department of Molecular Cell Biology, Sungkyunkwan University School of Medicine, Suwon, Republic of Korea; Department of Haematology, University of Cambridge, Cambridge, UK; Centro Nacional de Análisis Genómico (CNAG), Barcelona, Spain; Department of Medicine, University of Cambridge, Cambridge, UK; CIFAR Macmillan Multi-scale Human Programme, CIFAR, Toronto, Canada; Institute of Lung Health and Immunity, Helmholtz Munich, Member of the German Center for Lung Research (DZL), Munich, Germany; Division of Medical Oncology-Research, National Cancer Centre Singapore, Republic of Singapore

## Abstract

Blood provides an accessible window into human health, yet the absence of a peripheral blood mononuclear cells (PBMCs) reference with standardized immune cell annotations has constrained comparisons between single-cell RNA sequencing (scRNAseq) studies. Here, we present HARP (**H**uman Cell **A**tlas **R**eference for **P**BMCs), an integrated atlas of ∼9 million PBMCs from >2,600 donors across 15 studies spanning four continents, encompassing diverse populations, neonates to 97 years, across health and diverse immune-related diseases. We developed optimized integration workflows, including novel label-free metrics to assess integration quality, and generated community-driven consensus annotations for 192 immune cell subsets, identifying rare populations representing as few as 0.004% of PBMCs. HARP reveals coordinated cellular modules associated with age, sex, and disease, identifies female-biased interferon and inflammatory gene programs and their perturbation by COVID-19, and uncovers a novel sexual dimorphism in prostaglandin signaling. Finally, we introduce *scTiger,* a hierarchical label-transfer framework that accurately projects these 192 cell annotations onto ∼18 million additional PBMCs, providing a robust and transferable reference for harmonizing future PBMC studies. HARP provides a high-resolution community-driven reference atlas for PBMC annotation and analysis, as a basis for emerging clinical applications of single-cell genomics.

## Introduction

The complexity of the human immune system is reflected in the remarkable cellular diversity of blood, which comprises numerous transcriptional states shaped by aging, sex, disease, genetics, and environmental exposures. Blood is uniquely suited to high-resolution immune profiling because it is easily accessible, yields large numbers of cells from a single draw, and requires minimal processing. Quantification of major cell lineages already forms the basis of routine clinical testing^1^ and, even at this coarse resolution, supports diagnosis, disease monitoring, and risk stratification.

Single-cell RNA sequencing (scRNAseq) has transformed blood profiling by enabling the discovery of novel and rare immune cell types, including *AXL SIGLEC6*^Hi^ dendritic cells (DCs)^2^, early tissue-resident memory T cells^3^, and *CLU*^Hi^ monocytes that are associated with sepsis and adverse clinical outcomes^4^. Large-scale cohort studies comprising hundreds of donors have further demonstrated how transcriptionally defined cell states are shaped by factors such as age, sex, ethnicity^5–8^, and disease^4,9–12^. Together, these studies illustrate how increasing both donor numbers and cell counts enables discovery of rare cell populations while revealing significant demographic influences on cellular composition and function. Although several PBMC single-cell RNAseq atlases have recently been developed, they have generally been generated using a limited number of studies and vary substantially in cohort size and composition, experimental protocols, data processing pipelines, integration strategies, and cell annotation frameworks. As a result, existing references capture complementary aspects of human peripheral immunity but do not provide a harmonized community reference with standardized immune cell annotation, limiting reproducibility, interoperability, and direct comparison across studies.

The Human Cell Atlas (HCA) seeks to generate a comprehensive reference map of all human tissues that capture the breadth of human biological diversity^13^. Achieving this vision for the blood compartment requires coordination across the scientific community to harmonize and integrate landmark datasets generated across studies, assays, and diverse populations while preserving genuine biological variation. Beyond overcoming data integration challenges, a community reference atlas requires consensus cellular annotation grounded in established immunological knowledge that has broad community agreement, together with robust methods for transferring those annotations to newly generated datasets to facilitate meaningful comparisons. Several approaches have been taken to address this last goal, such as mapping datasets onto a shared low-dimensional space^3,14–16^ or utilizing machine learning–based classifiers^17^, with each offering distinct trade-offs in accuracy, scalability, computational efficiency, and interpretability.

Here, we present HARP (**H**uman Cell **A**tlas **R**eference for **P**BMCs), an HCA reference atlas comprising ∼9M cells from more than 2,600 donors across 15 landmark peripheral blood mononuclear cells (PBMCs) studies spanning four continents. By integrating diverse healthy and disease cohorts, including SARS-CoV-2 infection, we distinguish reproducible biological variation from study-specific technical effects. We develop quantitative and qualitative benchmarking strategies that evaluate integration quality while ensuring preservation of cell-specific transcriptional signatures. HARP provides consensus annotations for 192 immune cell subsets spanning abundant and very rare populations. Leveraging these harmonized annotations, we identify previously undescribed immune cell states shared across populations, uncover coordinated cellular and transcriptional programs associated with age, sex, and disease, and introduce *scTiger*, a hierarchical framework for accurate annotation transfer to independent datasets. Altogether, HARP establishes a harmonized community reference atlas for human PBMCs that enables reproducible biological discovery across diverse populations while establishing a robust foundation for high-resolution annotation, interpretation and harmonization of future PBMC single-cell studies.

## Results

### Building a multi-cohort PBMC reference atlas

We constructed the **H**uman Cell **A**tlas **R**eference for **P**BMCs (*HARP*) by integrating 15 large scRNAseq PBMC datasets generated across Asia, Australia, Europe and North America (Fig. 1A)^3,7–12,18–23^. These studies were selected through engagement with the

**Figure 1:**
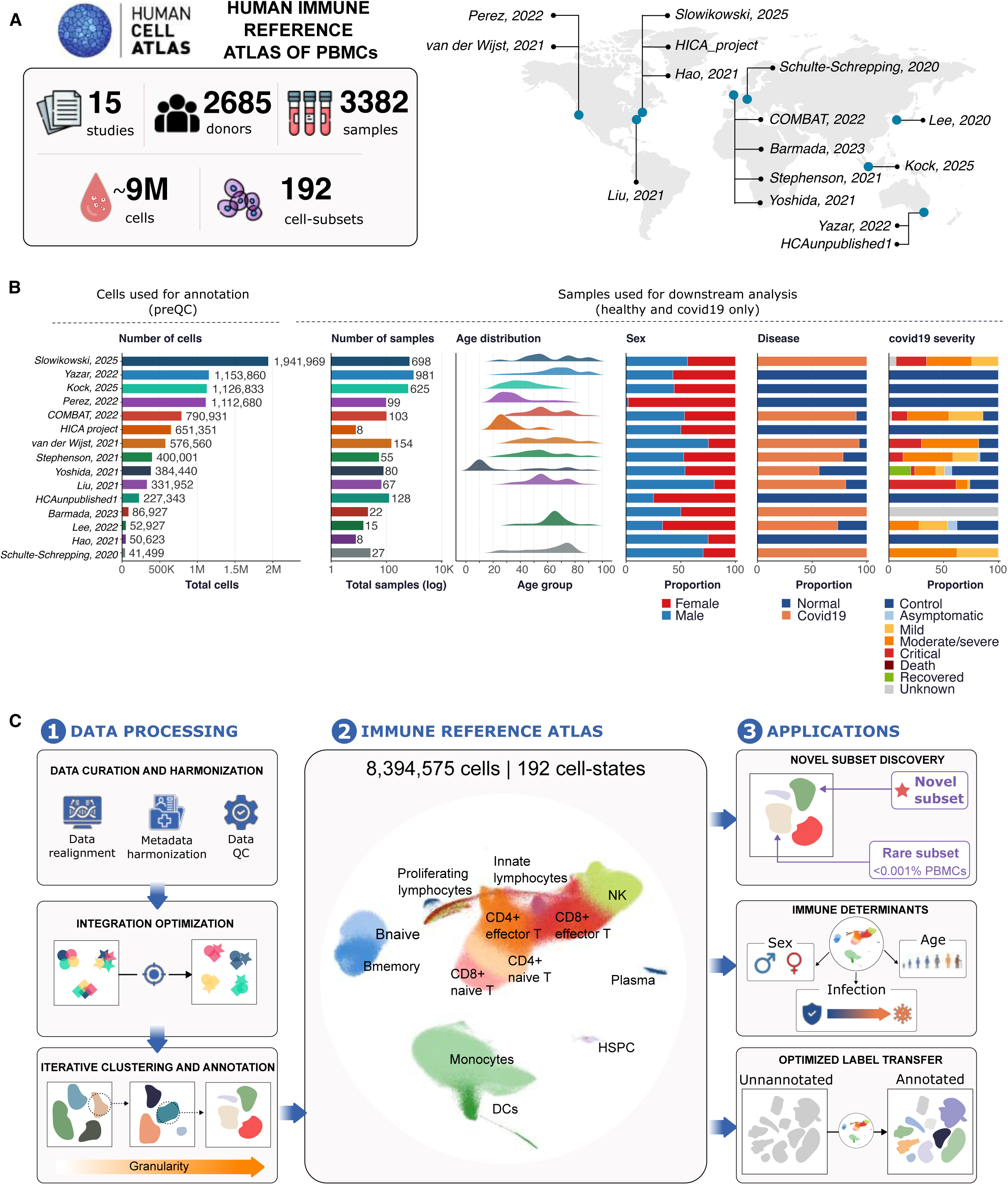
A human immune reference map. (A) Overview of the studies, donors, samples, and cells in HARP, together with the geographic distribution (one representative site indicated per study) of the landmark studies integrated. **(B)** Charts of key cohort demographics and sample metadata. Left to right: (1) Cell number before exclusion of low quality cells per study. (2) Number of samples per study. (3) Age distribution of samples within each study where data has been made available. (4) Proportion of female/male samples per study. (5) Proportion of healthy/SARS-CoV-2 samples per study. (6) Distribution of SARS-CoV-2 severity categories across samples per study. **(C)** Schematic of the computational workflow used for atlas construction and applications of HARP.

HCA Immune Bionetwork based on their size, population diversity, and use of a common droplet-based 10X Genomics platform. After removing low-quality cells and doublets, HARP comprised 8,394,584 cells from 2,685 donors, spanning newborns to 97 years of age and balanced for sex (49.8% female). Alongside healthy donors (46.6%), the atlas incorporated data from SARS-CoV-2 infected donors (40.7%) across the spectrum of disease severity (21.9% mild, 43.4% moderate/severe, 27.9% critical), together with additional disease samples from the contributing studies (Fig. 1B). Clinical and demographic metadata were systematically compiled and harmonized via a manual process involving study authors and clinician input (Methods) across all donors and studies, before downstream analysis (Fig. 1C). In sum, these datasets capture broad geographic, demographic, and clinical diversity, providing the foundation for a robust reference atlas of human PBMCs at high resolution (Fig. 1C).

### Biologically informed integration optimization enables robust integration across PBMC studies

Variation in sample preparation, sequencing, and bioinformatic preprocessing introduces technical differences that obscure biological signals and complicate integration across studies^24,25^. To establish a foundation for high-quality, biologically faithful consensus annotations, we systematically benchmarked each stage of atlas construction, including quality control, feature selection, and batch integration.

Initial integration of publicly deposited count matrices revealed a major limitation: differences in reference genomes used for alignment, gene annotations, and gene filtering reduced the shared feature space to only 16,004 genes (Fig. 2A). Critically, these missing genes included canonical lineage-defining genes, such as *CLEC9A* (cDC1), *TRAV1-2* (mucosal-associated invariant T [MAIT] cells), *TRDV2* and *TRGV9* (gamma-delta T [γδT] cells), cytokines (*IL5, IL21, IFNA1, IFNB1*), and CCL-family chemokine genes relevant to SARS-CoV-2 biology^9,26^ (Fig. 2A), illustrating how differences in upstream processing can compromise biological interpretation. Realignment of raw FASTQ files using a unified reference genome and gene annotation (GRCh38-GENCODE v32) and a standardized preprocessing pipeline (Methods) substantially reduced variance explained by technical covariates (Supplementary Fig. 1A), indicating reduced batch effects from bioinformatic processing alone.

**Figure 2:**
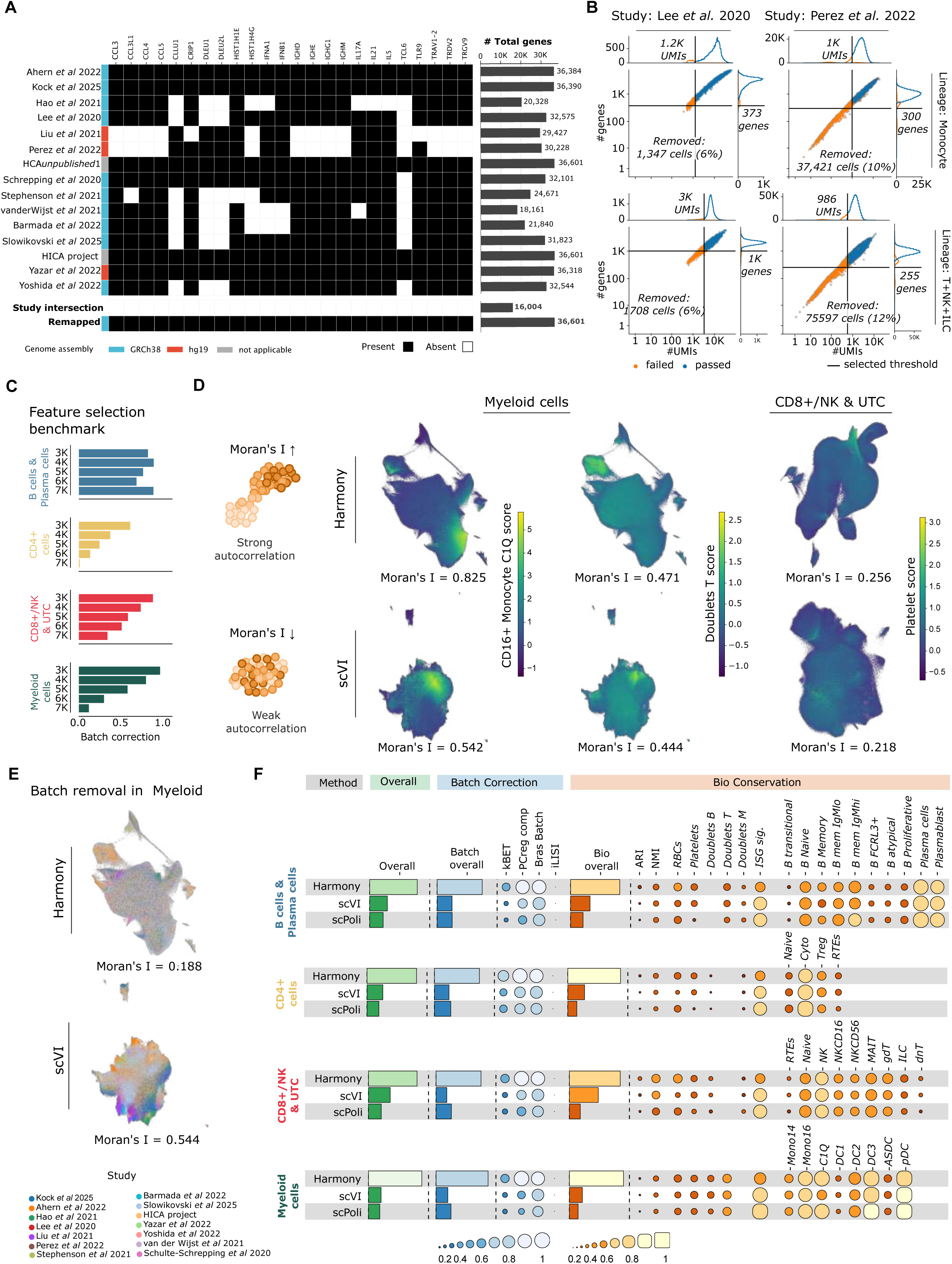
Benchmarking and optimization of data integration reveals biologically informed strategies for robust PBMC atlas construction. **(A)** Overview of biologically important genes missing from publicly available versions of the datasets in this study. **(B)** Examples of fitted Gaussian distributions for QC metrics (x axis: number of UMIs, y-axis: number of detected genes) by lineage (top: monocytes, bottom: T, NK, and innate lymphoid cells [ILC]) and study (columns), including automated cut-offs for low-quality cells. **(C)** Bar graphs of scaled, average batch correction scores computed as a composite of established integration metrics, split by lineage, for different numbers of highly variable genes. **(D)** Left: Schematic of Moran’s I metric. Right: Moran’s I scores projected onto UMAP embeddings for the indicated lineages. Embeddings were derived using Harmony (top) and scVI (bottom) during hyperparameter tuning. Cells were scored for their expression of a CD16+ monocyte (left), T cell doublet (middle) and platelet (right) Moran’s I score. Higher Moran’s I scores indicate stronger autocorrelation of the indicated gene modules. **(E)** Application of Moran’s I score^87^ to assess degree of batch mixing across studies, with lower scores indicating better mixing for example UMAP embeddings generated with Harmony (top) and scVI (bottom). The Moran’s I was calculated directly on study covariate and the kNN graph. **(F)** Benchmark of integration methods, with overall scores, batch correction metric scores, and biological conservation metric scores (including scIB^32^ and Moran’s I scores) depicted per lineage.

Differences in quality control metric distributions across datasets, caused by different assay sensitivities or study-specific filtering decisions, introduced compositional differences that generated clusters enriched for low-quality cells, impairing downstream integration (Supplementary Fig. 1B). To address this, we applied dataset- and lineage-specific filtering (Fig. 2B) by fitting Gaussian curves to total counts, numbers of detected genes, fraction of mitochondrial reads, and removing outliers as described previously^7,27^. Using a composite batch correction score (Methods), we found that increasing the numbers of features from 3000 to 7000 highly variable genes progressively increased study-specific clustering and reduced overall integration quality (Fig. 2C), consistent with previous observations^28^. We therefore considered smaller feature sets for subsequent integration benchmarking to ensure better batch mixing required for later cell type annotation.

We next compared global and lineage-specific integration strategies (Supplementary Fig. 1C, D) across three scalable batch correction methods (Harmony^29^, scVI^30^ and scPoli^31^), together with global- and lineage-specific feature selection (CD4 T cells, CD8 T cells, B cells, myeloid, other), as the performance of different integration methods is known to vary according to underlying dataset characteristics^32^. Because immune cell lineages are defined by distinct transcriptional programs, we hypothesized that lineage-specific feature selection would better preserve fine-grained biological variation than a single global model (Supplementary Fig. 1C). Each integration method was systematically optimized across multiple hyperparameter settings and evaluated using complementary measures of batch correction and biological conservation metrics^32–34^ (Fig. 2E), while taking into account known technical covariates and potential confounding biological covariates such as age, sex and disease (Methods; Supplementary Fig. 2A-C). Among the evaluated technical covariates, we identified the sequencing library ID as the optimal, minimally confounded integration variable (Supplementary Fig. 2A, C). Biological conservation was assessed using a harmonized version of the original authors’ annotations derived using CellHint^17^, together with a targeted, label-free Moran’s I^35^-based metric^36^ that quantifies the preservation of lineage- and disease-specific gene signatures within the integrated neighborhood graph (Methods). We reasoned that an optimal embedding should preserve well-defined immune populations, separate technical artefacts such as doublets and platelet contamination, and achieve effective mixing across studies and batch variables.

The Moran’s I metric revealed biologically meaningful differences between embeddings from different integration methods that could not be resolved by conventional, label-based integration metrics. For example, CD16 C1Q^Hi^ monocytes (Fig. 2D; Supplementary Fig. 3A), cDC2, and γδ T-cells (Supplementary Fig. 1E, F; Supplementary Fig. 3B, C) were more distinctly separated from related cell subsets using Harmony compared to scVI. Similarly, there were clear differences in how integration methods identified putative doublets and platelet contamination (Fig. 2D), with Harmony distinguishing these contaminants within the CD8 T-cell lineage. Finally, we demonstrate the usage of Moran’s I to assess batch mixing between studies (Fig. 2E), with a lower Moran’s I score in Harmony versus scVI reflecting good batch mixing. Across all benchmarking strategies, lineage-specific Harmony-integration consistently provided the best balance between batch correction and preservation of biologically meaningful immune cell subsets (Fig. 2F; Supplementary Fig. 1D). Accordingly, this strategy formed the foundation of the HARP reference atlas.

### Identification of fine-grain cell subsets

The final cell embeddings were clustered to the resolution where no cluster represented study-, technology-, or donor-specific clusters before annotation. This resulted in a final HARP atlas (8,394,584 cells) with 192 distinct cell subsets, which were defined through an iterative community cell annotation process, with each major lineage independently curated by HCA Immune Bionetwork experts located at institutes in Cambridge (UK), Boston, and Singapore (Fig. 3A, Extended Fig. 1, Supplementary Fig. 4-7). The final embeddings, fine-grain annotation, and supporting annotation rationale are publicly available through the HCA Cell Annotation Platform. Cell identities are organized into a four-level hierarchy, progressing from major immune lineage (level 1; myeloid, B and plasma, CD4 T, CD8 T and innate lymphoid, hematopoietic stem and progenitor cells [HSPC]), to canonical cell types (level 2; e.g., Monocytes, Dendritic cells [DC]), canonical cell subsets (level 3; e.g., CD14^Hi^ monocytes, CD16^Hi^ monocytes), and fine-grained cell subsets (level 4) that capture recurrent biological programs, such as activation (*FOS-JUN, NR4A2, HLA, CISH*) and cytokine response (*IL1B, IFN*). Of the 192 consensus cell subsets, 66 represent rare or newly resolved subsets, which are bolded in Extended Fig. 1.

**Figure 3:**
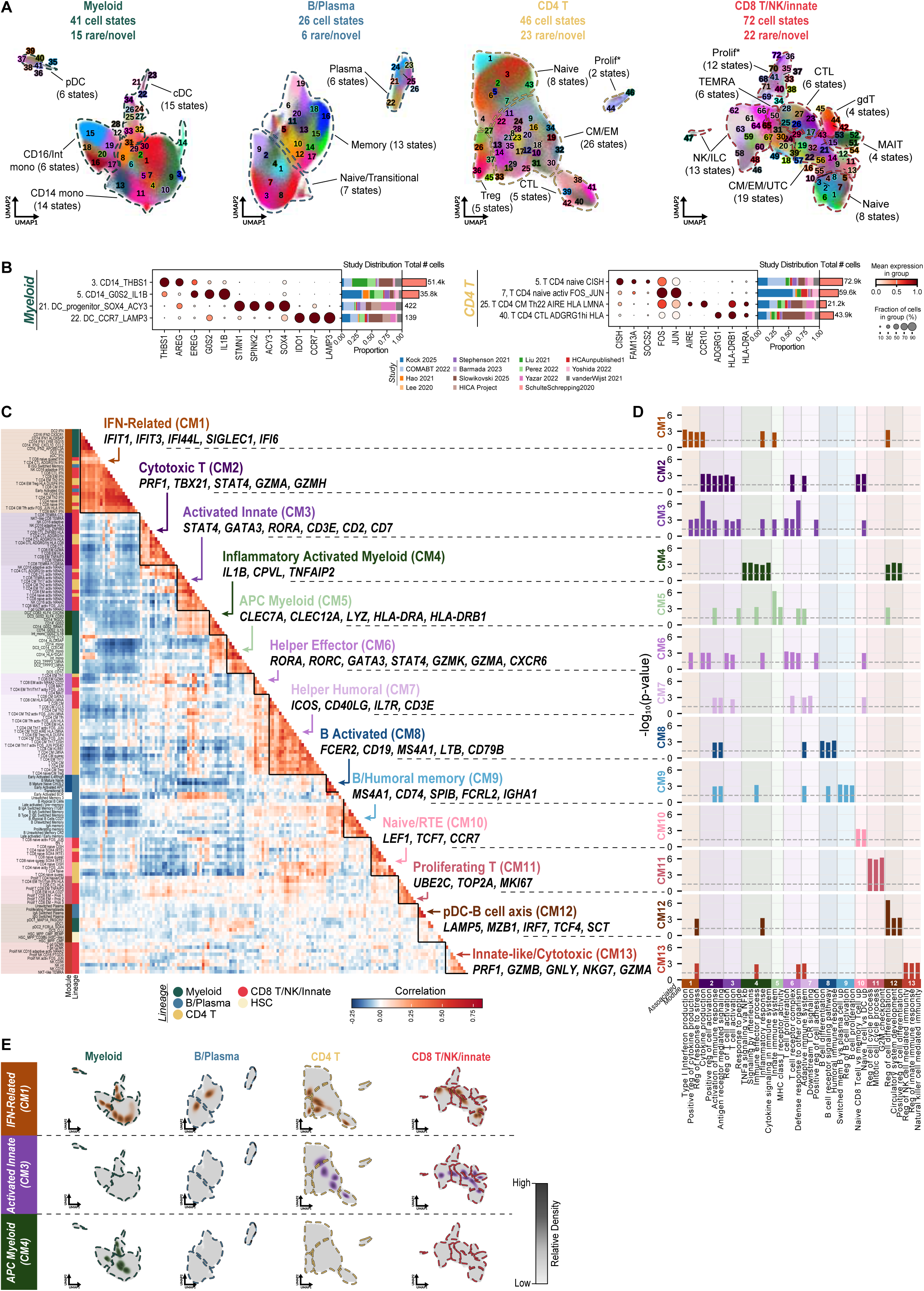
Fine-grained cell subset annotation across the HARP atlas reveals coordinated, cross-lineage co-abundance cellular modules. **(A)** UMAP embeddings of four major immune lineages, representing 192 granularly annotated cell subsets [185 after excluding technical clusters] and their respective sublineages. Total number of cell subsets and identified rare/novel subsets are indicated above each compartment. See Supplementary Figures 5-7 for sublineage UMAPs and dotplot of markers, and Supplementary Figure 7 for the 7 HSPC cell subsets. (**B**) Dot plots of mean marker gene expression levels for selected myeloid (left) and CD4 T cell (right) novel/rare cell subsets, annotated by study distribution and total number of cells. Stacked horizontal bar plots represent the study distribution proportion for each cell subset. Red horizontal bar plots indicate the total number of cells per cell subset. **(C)** Heatmap showing pairwise cell type correlations as calculated from sample-level cell type proportions. Cellular Modules (CMs) were defined using hierarchical clustering of correlation values. Genes associated with each CM were identified by applying OPLS to pseudobulked gene expression to identify a gene vector with the highest discriminatory power to distinguish cell subsets within each CM from all other cell subsets. Key representative genes for each CM from the top 40 genes with highest OPLS loading for each CM are indicated. (**D**) Bar charts showing results from msigDB over-representation analysis on top 40 highest loading genes from OPLS analysis per CM. For each CM, 3-5 significant pathways (x-axis) are shown along with their p-values (y-axis); dashed line indicates threshold p-value of 0.05. Absence of bars indicates results that did not reach statistical significance. (**E**) Density plot UMAPs of three representative CMs (Supplementary Figure 4 for other CMs). Density scales are normalized per cell subset to emphasize localized peaks.

Within the monocyte and DC compartment (2,315,268 cells), HARP resolved 41 transcriptionally distinct cell subsets spanning classical (14 subsets), nonclassical (6 subsets), and intermediate monocytes (2 subsets), together with 21 distinct DC subsets. These populations were organized by recurrent transcriptional programs related to interferon signaling (*IFN, ISG15*), TNF signaling and IL1B production (*NFKBIA, TNFAIP3*, *IL1B, G0S2:* CD14_G0S2, CD14_G0S2_IL1B, Int_mono_G0S2_IL1B), sepsis-associated response^4^ (*CLU, RETN:* CD14_ALOX5AP, CD14_IFN1_ALOX5AP, CD14_LY6E_ISG15, DC3_CLU), and proliferation (*MKI67, STMN1*), and a previously reported natural killer (NK)-like monocyte signature (CD14_MKI67_NKG7_ALOX5AP)^2^ (Extended Fig. 1, Supplementary Fig. 5A, B).

The DC compartment included canonical cDC1 and cDC2 populations expressing multiple distinct gene programs. It also revealed extensive heterogeneity within pDC related to activation state (pDC1, pDC1_MAP1A_PASCIN1, pDC1_NIBAN3_FCHSD2, pDC2_FCRLA_SOX4, pDC3_IRF4_CXCR4, pDC_IFN), as well as rare populations including circulating migratory *CCR7*^Hi^ *LAMP3*^Hi^ DC (DC_CCR7_LAMP3), and a *SOX4*/*ACY3*-expressing progenitor-like DC state (DC_progenitor_SOX4_ACY3)^2^ (Fig. 3A, B - left panel, Extended Fig. 1).

The B and plasma cell compartment (893,376 cells) resolved 26 distinct cell subsets spanning naive/transitional (7 subsets), unswitched and switched memory (13 subsets), and antibody-secreting plasma cells (6 subsets). Activation signatures followed distinct transcriptional trajectories across the B cell lineage, with early activation in naive B cells marked by *CD83* and *IL4R* expression, whereas activated memory B cells preferentially expressed *NR4A1* and *NR4A2* (Extended Fig. 1, Supplementary Fig. 5C, D). HARP also resolved heterogeneity within atypical B cells associated with chronic activation or antiviral responses^37^ (*ITGAX, TBX21*: Atypical B cells, Atypical B cells CD27).

Within the CD4 T cell compartment (2,399,830 cells), HARP identified 46 distinct cell subsets, including canonical naive (8 subsets), central memory (5 subsets), regulatory (6 subsets), helper (19 subsets), effector (2 subsets), and cytotoxic (5 subsets) populations (Extended Fig. 1, Supplementary Fig. 6A, B). Across multiple differentiation states (e.g., naive, Th1, Th2, Th17 Tfh, cytotoxic T lymphocytes (CTL)), two conserved early activation programs centered on AP-1 (*FOS*/*JUNB*) or *NR4A2* were repeatedly observed (Extended Fig. 1). The atlas further resolved a rare *CISH*-expressing naive population previously implicated in inflammaging^38^ together with a previously undescribed *AIRE-*expressing helper T cell subset (T_CD4_CM_Th22_AIRE_HLA_LMNA). *AIRE* is classically expressed by medullary thymic epithelial cells but with unclear function in the context of circulating Th22 cells (Fig. 3B - right panel).

The CD8 T cell, unconventional T cell (UTC), and innate lymphoid cell compartment (2,859,350 cells) comprised 72 unique subsets. As observed in the myeloid and CD4 T cell compartments, recurrent transcriptional programs associated with early activation (*FOS*/*JUNB*, *NR4A2),* TNF signaling (*TNFAIP3, JUND*), interferon signaling (*IFIT3, IFI44L*), MHC class II (*HLA-DRB1, HLA-DRA*), and cytotoxicity (*GZMA, GZMH*) were repeatedly observed across multiple differentiation states, including CD8 naive, central memory (CM), effector memory (EM), terminally differentiated effector memory cells re-expressing CD45RA (TEMRAs), NK, mucosal-associated invariant T (MAIT), and γδT cells (Extended Fig. 1, Supplementary Fig. 6C, Supplementary Fig. 7A). HARP recapitulated previously described heterogeneity within the TEMRA cells^39^ and HOBIT-expressing cytotoxic effector T cells^40^. We also identified rare double positive and double negative T cell populations together with innate lymphoid cell (ILC) and proliferating subsets (Supplementary Fig. 6D).

Finally, despite representing only 11,247 cells, the HSPC compartment resolved 7 transcriptionally distinct progenitor subsets (Extended Fig. 1, Supplementary Fig. 7B-C). Despite their rarity in peripheral blood, these populations recapitulated the major stages of haematopoietic differentiation typically observed in bone marrow, spanning primitive HSC-like states through lymphoid, myeloid, and erythro-megakaryocytic-primed progenitors^41–43^.

Collectively, these analyses show that HARP captures not only canonical immune cell identities but also recurrent transcriptional programs that transcended lineage boundaries, including interferon responses, inflammatory activation, proliferation, and immediate early activation (Extended Fig. 1). Rather than defining independent immune cell subsets, these conserved programs were repeatedly observed across distinct immune cell types while remaining embedded within lineage-specific differentiation states, providing a unified framework for comparing immune cell activation across health and disease. This observation suggested that coordinated biological programs extend across canonical immune cell identities, motivating a systematic analysis of cross-lineage cellular modules.

### Co-Abundance Cellular Modules (CMs) reveal coordinated immune programs

We hypothesized that changes in immune-cell abundance occur in a coordinated manner, with multiple cell subsets expanding or contracting together in response to different demographics, such as age and sex. To identify these coordinated responses, we computed a pairwise Pearson correlation matrix based on the relative abundances of all annotated cell subsets and applied hierarchical clustering, identifying 13 distinct co-abundance Cellular Modules (CMs) comprising cell subsets whose abundance consistently co-varied across donors (Fig. 3C).

To characterize the biological functions underlying each CM, we applied orthogonal partial least squares (OPLS)^44^ to identify transcriptional programs that define module membership rather than canonical immune cell identity. The 40 genes with the highest module-specific OPLS component loadings were subjected to over-representation analysis (ORA), allowing each CM to be assigned a functional biological annotation (Fig. 3C, D, Methods). These analyses assigned functional identities to individual CMs, including interferon signaling (CM1: *IFIT1, IFIT3*), inflammatory myeloid cell activation (CM4: *IL1B, TNFAIP2*), naive lymphocyte identity (CM10: *LEF1, TCF7)*, proliferation (CM11: *MKI67, TOP2A*), and T and Innate cytotoxicity (CM2 and CM13: *PRF1, GZMB*). ORA further demonstrated significant enrichment for pathways including type I interferon production (CM1), interleukins and TNFα via NF-kB signaling (CM4), cell-cycle regulation (CM11), and NK cell-mediated immunity (CM13) (Fig. 3D).

The resulting CMs varied in both lineage composition and biological scope, ranging from modules that spanned multiple immune lineages to those restricted to individual lineages. For instance, CM1 (IFN-Related) encompassed interferon-responsive subsets distributed across all four major lineages, demonstrating that shared transcriptional programs can transcend canonical immune identities (Fig. 3C, E). In contrast, other CMs displayed stricter lineage specificity: CM3 (Activated Innate) encompassed activated subsets spanning the CD4 and CD8/innate compartments, whereas CM4 (Inflammatory Activated Myeloid) was restricted to the myeloid lineage (Fig. 3C, E and Supplementary Fig. 4). Importantly, even within individual lineages, subsets belonging to the same CM occupied distinct areas of the low-dimensional embedding rather than forming a single transcriptionally defined cluster (Fig. 3A, E), indicating that these modules represent shared biological programs superimposed on canonical immune cell identities rather than artifacts of overclustering.

### Age and sex remodel immune cell composition

We next examined how age and sex remodel immune cell composition in the context of coordinated CMs identified above. Linear modelling revealed widespread remodeling of both adaptive and innate compartments with age (Supplementary Fig. 8A). Among the most significant age-associated changes, CM10 (Naive/RTE-associated populations) showed a progressive decline across the lifespan, including in *SOX4*^Hi^ recent thymic emigrants (RTEs)^45^. In contrast, CM7 (helper/humoral) and CM2 (cytotoxic) progressively increased with age (Fig. 4A, B).

**Figure 4.**
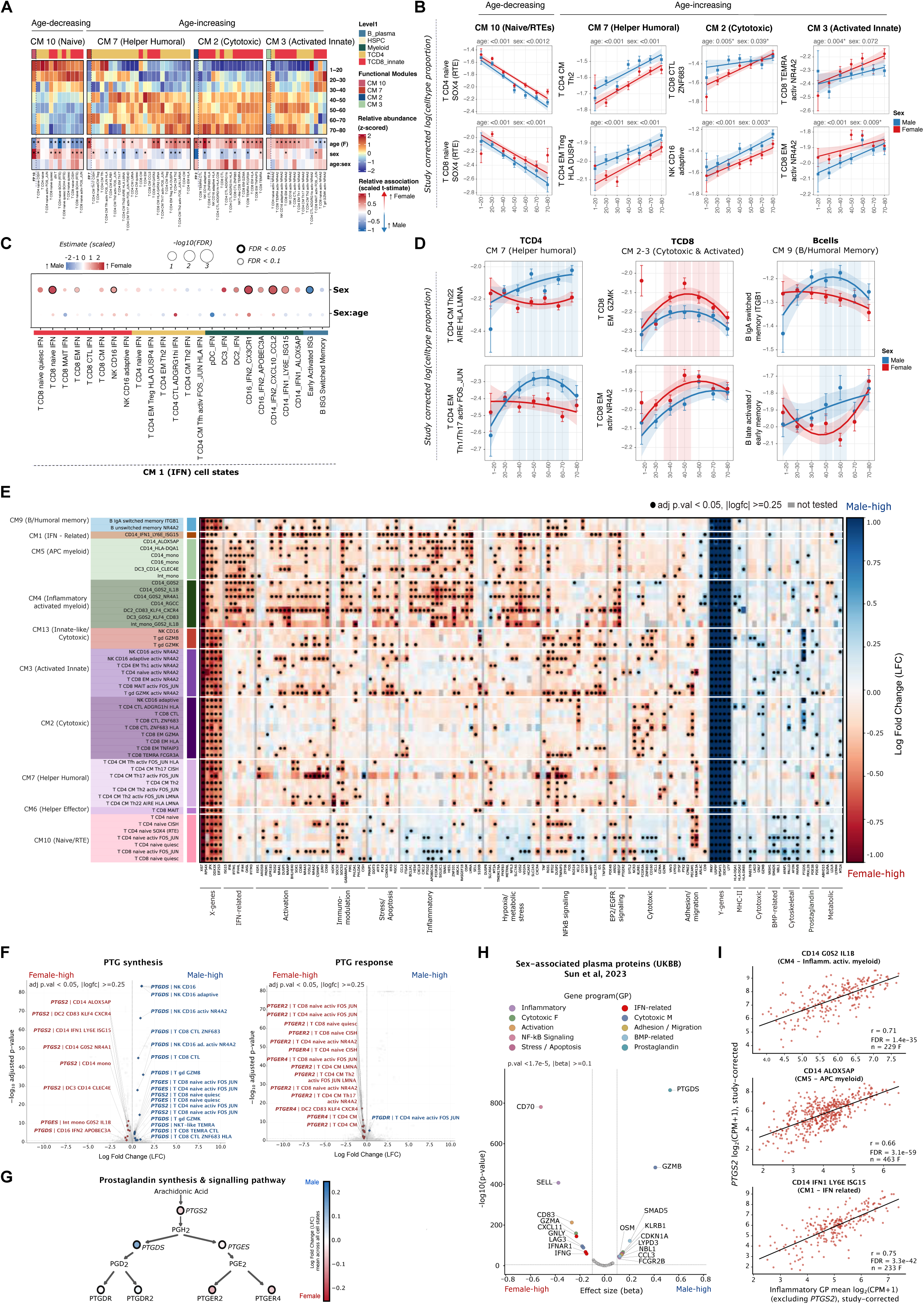
Age- and sex-associated variation in immune cell subset abundance and cell subset-specific gene expression in healthy donors. (**A**) Heatmap indicating relative shifts in cell subset abundance with age; cell subsets are categorized by CMs. Upper panel shows z-scored, study-corrected cell subset proportions averaged within age bins. Lower panel shows signed t-statistics from the linear mixed-effects model (LMM; log10(proportion) ∼ age × sex + (1 | study)), where each coefficient was tested using Satterthwaite-approximated **t**-tests. Top colour bar indicates the broad lineage (Level 1 annotation) the cell subset belongs to. Asterisks denote BH-adjusted p < 0.05. Non-significant terms (BH-adjusted p ≥ 0.10) are down-weighted (×0.25) to visually de-emphasise unconfirmed trends. (**B**) Examples of cell subsets displaying significant linear associations with age and/or sex. Points represent mean study-corrected log10(proportion) values within each age bin separated by sex; error bars indicate ± SEM. Lines and bands show linear fits with 95% confidence intervals. Reported BH-adjusted p-values are from Satterthwaite-approximated **t**-tests on the LMM coefficients (log10(proportion) ∼ age × sex + (1 | study)). (**C**) Dotplot summarising sex and sex:age interaction effects across CM1 (IFN-related) cell types. Dot color indicates scaled effect size and dot size reflects statistical significance (-log10 FDR). Black and grey borders indicate BH-adjusted p<0.05 and p<0.1, respectively. (**D**) Examples of cell subsets exhibiting significant non-linear shifts in abundance with age, plotted separately by sex. Lines and bands show quadratic fits with 95% confidence intervals indicated. Shaded regions indicate age intervals in which female and male confidence intervals do not overlap. Cell subsets were selected based on a significant improvement in fit of the quadratic-by-sex model over a sex-only null model (F-test via ANOVA model comparison, BH-adjusted p < 0.05). (**E**) Heatmap of differential expression of genes in males versus females organised into manually curated gene programs (GPs) (x axis) and CMs (y axis). Indicated genes were significantly differentially expressed in at least four cell subsets. Only cell subsets contributing ≥10 DEGs with concordant LFC direction are shown. Colour indicates log-fold change (LFC; red indicates female-high, blue indicates male-high expression). Filled circles denote BH-adjusted p < 0.05 and |LFC| ≥ 0.25; grey-coloured combinations were not tested due to their limited power. (**F**) Volcano plots showing sex-associated LFC (x axis) versus –log₁₀ adjusted p-value (y axis) for genes related to prostaglandin synthesis GP (**PTGS2, PTGDS, PTGES**; left), and response (**PTGER2, PTGER4, PTGDR**). Labelled points meet thresholds of BH-adjusted p < 0.05 and |LFC| ≥ 0.25. Female-high associations are shown in red, male-high in blue. (**G**) Schematic of prostaglandin synthesis and response pathway. Coloured circles indicate mean LFC across all cell subsets for the gene. (**H**) Sex-associated plasma proteins corresponding to recurrent sex-biased GPs. Volcano plot showing sex associations from the UK Biobank Pharma Proteomics Project Olink data for proteins matching genes from GPs, with selected IFN-related proteins included. The x-axis shows the male-female difference in protein abundance, where positive values indicate higher abundance in males and negative values indicate higher abundance in females. The y-axis shows - log10(p-value). Dotted vertical lines indicate the effect-size cutoff |beta| > 0.1, and the horizontal dotted line indicates the significance cutoff P < 1.7 × 10⁻⁵. Colored and labelled proteins pass both cutoffs and GP assignment; grey proteins do not. (**I**) Scatter plots showing the correlation between **PTGS2** expression (log2p (CPM +1)) and the mean expression of all other innate inflammatory GP genes (log2p (CPM +1), excluding **PTGS2**) for individual female donors, in three myeloid cell subsets: CD14_G0S2_IL1B (from CM4, inflammatory activated myeloid), CD14_ALOX5AP (from CM5, APC myeloid), and CD14_IFN1_LY6E_ISG15 (from CM1, IFN-related). Pearson r, FDR, and sample size (n) are shown per panel.

Focusing on individual cell subsets within these modules recapitulated multiple established features of immune aging, including: expansions of CD4 Th1 (CM3: T_CD4_EM_Th1_activ_NR4A2), Th2 (CM7: T_CD4_CM_Th2), HLA-DR+ (CM7: T_CD4_EM_HLA), and memory Tregs (CM7: T_CD4_CM_Treg; T_CD4_EM_Treg_HLA_DUSP4), together with increased CD16 monocytes (CM5: CD16_mono) and reductions in *SOX4*^Hi^ RTEs (CM10: T CD4 naive SOX4 (RTE); T CD8 naive SOX4 (RTE)), consistent with previous studies^6,46–49^. The recovery of these well-established age-associated changes demonstrates that HARP preserves known biological signals while integrating diverse cohorts.

Beyond these established patterns, HARP identified previously unrecognized age-associated remodeling across multiple immune lineages. A particularly striking finding was the coordinated expansion of an NR4A2-associated CM spanning CD4 T, CD8 T, and NK cells (CM3: T_CD8_TEMRA_NR4A2; NK_CD16_adaptive_activ_NR4A2; T CD8_EM_activ_NR4A2, NK_CD16_activ_NR4A2, T_CD8_CTL_activ_NR4A2, T_CD4_EM_Th1_activ_NR4A2). Although *NR4A2* is a transcription factor that has previously been linked to T-cell exhaustion^50^ and chronic activation^51^, its coordinated association with human immunoaging has not been described. Additional age-associated remodeling included expansion of lymphocyte subsets expressing early response genes (including *NR4A2*, *FOS*, and *JUN*), cytotoxic CD4 T cells^52^ (CM2: T_CD4_CTL_ADGRG1lo; T CD4_CTL_ADGRG1hi_HLA), CD8 TEMRA cells (CM2: T_CD8_TEMRA_CTL), and activated memory B cell populations (CM6: B proliferating memory; B late activated/pre-memory; B late activated/early memory), revealing coordinated remodeling that extends beyond canonical immunosenescence signatures.

Sex-associated differences in immune-cell abundance were similarly widespread and highly coordinated (Fig. 4B, C, Supplementary Fig. 8A). Males preferentially exhibited expansion of helper and cytotoxic lymphocyte populations (e.g., activated CD4 Tfh, Th2, Th17, Th22), together with CD8 T cell and NK cell subsets (CM2 and CM7). In contrast, females showed increased abundance of naive populations, including RTEs (CM10), together with a prominent IFN-responsive signature within CM1 driven largely by CD14 monocyte, CD16 monocyte, and DC IFN cell subsets (CD14_IFN2_CXCL10_CCL2, CD16_IFN2_CX3CR1, DC2_IFN populations; Fig. 4C). No significant age-by-sex interactions were detected, indicating that these differences are largely maintained throughout the lifespan.

Because many immune differences emerge during reproductive life, we next examined whether sex-associated abundance changes followed non-linear trajectories with age (Fig. 4D). Several CD4 helper populations, including Th1/Th17, and the newly defined Th22 AIRE^Hi^ subset, diverged predominantly later in life, with progressively higher abundance in males. In contrast, specific activated and effector-memory CD8 T-cell populations (i.e., T_CD8_EM_GZMK and T_CD8_EM_activ_NR4A2) were preferentially enriched in females between approximately 30 and 50 years of age. Similar non-linear dynamics were observed within the B-cell compartment, where males showed increased abundance of memory-associated populations during midlife, including IgA-switched memory *ITGB1*^Hi^ B cells.

### Sex-biased gene programs reveal coordinated immune signaling networks

We next explored whether these sex-associated differences extended beyond cell abundance to coordinated transcriptional programs (Supplementary Fig. 8B). Analysis of recurrent differentially expressed genes (DEGs) shared across at least four cell subsets (adjusted *P* < 0.05, |logFC| > 0.3) identified 246 genes across 108 cell subsets. Over-representation analysis of these recurrent DEGs revealed marked sexual dimorphism in immune signaling, with females enriched for inflammatory pathways, including TNF signaling via NF-kB, inflammatory response, and IFNγ response, whereas males were preferentially enriched for pathways related to antigen presentation, cell adhesion, and cyclic AMP (cAMP)-mediated signaling (Supplementary Fig. 9A).

To identify higher-order transcriptional organization, recurrent DEGs were consolidated into biologically coherent gene programs (GPs) based on shared biological functions and consistent expression patterns across cell subsets, yielding 18 GPs spanning 10 CMs (Fig. 4E, Supplementary Fig. 8C). This revealed coordinated sex-biased transcriptional programs extending beyond sex chromosome-linked genes. Female-biased GPs were enriched for inflammatory, metabolic, adhesion, and prostaglandin E2 receptor (EP2)-related pathways, whereas male-biased GPs were dominated by MHC class II antigen presentation, cytotoxicity, and prostaglandin-related genes.

One of the strongest sex-biased pathways involved coordinated differences in prostaglandin synthesis and response genes (Fig. 4F, G). *PTGS2/PTGES* was significantly higher in females across multiple myeloid subsets, including CD14/intermediate monocyte subsets and DC populations, whereas naive T populations showed higher *PTGS2/PTGES* in males. Conversely, *PTGDS* was consistently higher in male across NK and cytotoxic T cell subsets (Fig. 4F). Females additionally exhibited higher expression of prostaglandin receptors *PTGER2* and *PTGER4* across multiple naive and memory T cell populations and DCs. Collectively, these findings delineate a previously undescribed sex dimorphism across the arachidonic acid–PGH2–PGD2/PGE2 axis, characterized by female-biased myeloid *PTGS2/PTGES* expression, lymphoid *PTGER2/PTGER4* signaling, and male-biased innate/lymphoid *PTGDS* expression (Fig. 4G, Supplementary Fig. 9B).

To determine whether this prostaglandin axis generalized across populations, we examined *PTGDS* and *PTGS2* expression in the Chinese Immune Multi-Omics Atlas^49^ (Supplementary Fig. 9C, D). Consistent with our findings, *PTGDS* showed reproducible male-biased expression across NK-ILCs, unconventional T cells, CD8 T cells, CD4 T cells, and B cells, with the strongest and most significant effects observed in NK, CD8 CTL, and γδ T cell subsets (Supplementary Fig. 9C). *PTGS2* expression likewise demonstrated female-biased expression within inflammatory classical monocyte subsets (cMono IFI44L and cMono *CXCL10)* (Supplementary Fig. 9D), supporting conservation of this immunomodulatory pathway^53^ across independent cohorts.

We next asked whether these transcriptional programs were reflected at the protein-level. Mapping non-sex-chromosome GP genes onto the UK Biobank Pharma Proteomics Project Olink panel^54^ identified 19 proteins with significant sex associations (unadjusted *P* < 1.7 × 10⁻⁵, |beta| > 0.1) (Fig. 4H). Concordant protein-level differences supported several transcriptional axes, including female-biased inflammatory, interferon, and TNF-associated proteins (IFNG, IFNAR1, CXCL11, CD83, SELL, TNF, CD70) and male-biased prostaglandin, cytotoxic, and SMAD-related proteins (PTGDS, GZMB, SMAD5). A subset of proteins with smaller effect sizes showed discordant plasma associations, likely reflecting expression in distinct cell subsets that cannot be resolved by plasma proteomics. Altogether, these findings provide orthogonal protein-level validation of the core sex-biased transcriptional programs identified by HARP. Finally, to investigate whether *PTGS2* participates in the broader inflammatory program observed in females, we examined correlations between *PTGS2* expression and inflammatory GP genes across myeloid subsets exhibiting the strongest inflammatory enrichment (CM1, CM4, CM5) (Fig. 4I; Supplementary Fig. 9E). *PTGS2* expression showed strong positive correlations with inflammatory genes across multiple myeloid subsets, including CD14_G0S2_IL1B, CD14_ALOX5AP, and CD14_IFN1_LY6E_ISG15, consistent with coordinated regulation of *PTGS2* within inflammatory transcriptional programs^55^.

Together, these analyses reveal that sex is a major determinant of immune-system organization across the human lifespan. Females exhibited coordinated inflammatory, interferon, and PGE₂-associated programs spanning multiple immune compartments, whereas males preferentially exhibited antigen presentation, cytotoxicity, and PGD₂ synthesis-associated programs. These sex-biased transcriptional programs were independently validated across an external immune atlas and large-scale plasma proteomics, demonstrating that coordinated immune organization is reflected not only across individual cell subsets but also across higher-order multi-cellular immune programs.

### COVID-19 broadly remodels immune-cell composition, while specific subsets track with disease severity

Multiple large PBMC scRNAseq datasets were generated during the SARS-CoV-2 pandemic, providing an opportunity to assess biological conservation across the integrated HARP atlas while leveraging its fine-grained annotation to resolve previously unrecognized disease-associated patterns. We therefore examined changes in immune-cell abundance across CMs, separately comparing individuals with COVID-19 to healthy controls and evaluating associations with disease severity across an ordinal scale comprising mild, moderate-to-severe, and critical disease.

COVID-19 was associated with coordinated remodeling across multiple CMs. Most naive lymphocyte subsets within CM10 and helper T cell subsets within CM7 were reduced, consistent with the well-established lymphopenia associated with COVID-19^56^. Likewise, most non-inflammatory myeloid subsets within CM5 were depleted, whereas IFN-related subsets within CM1 expanded, consistent with findings from the original studies^9,10,20^ (Fig. 5A, Supplementary Fig. 10A).

**Figure 5:**
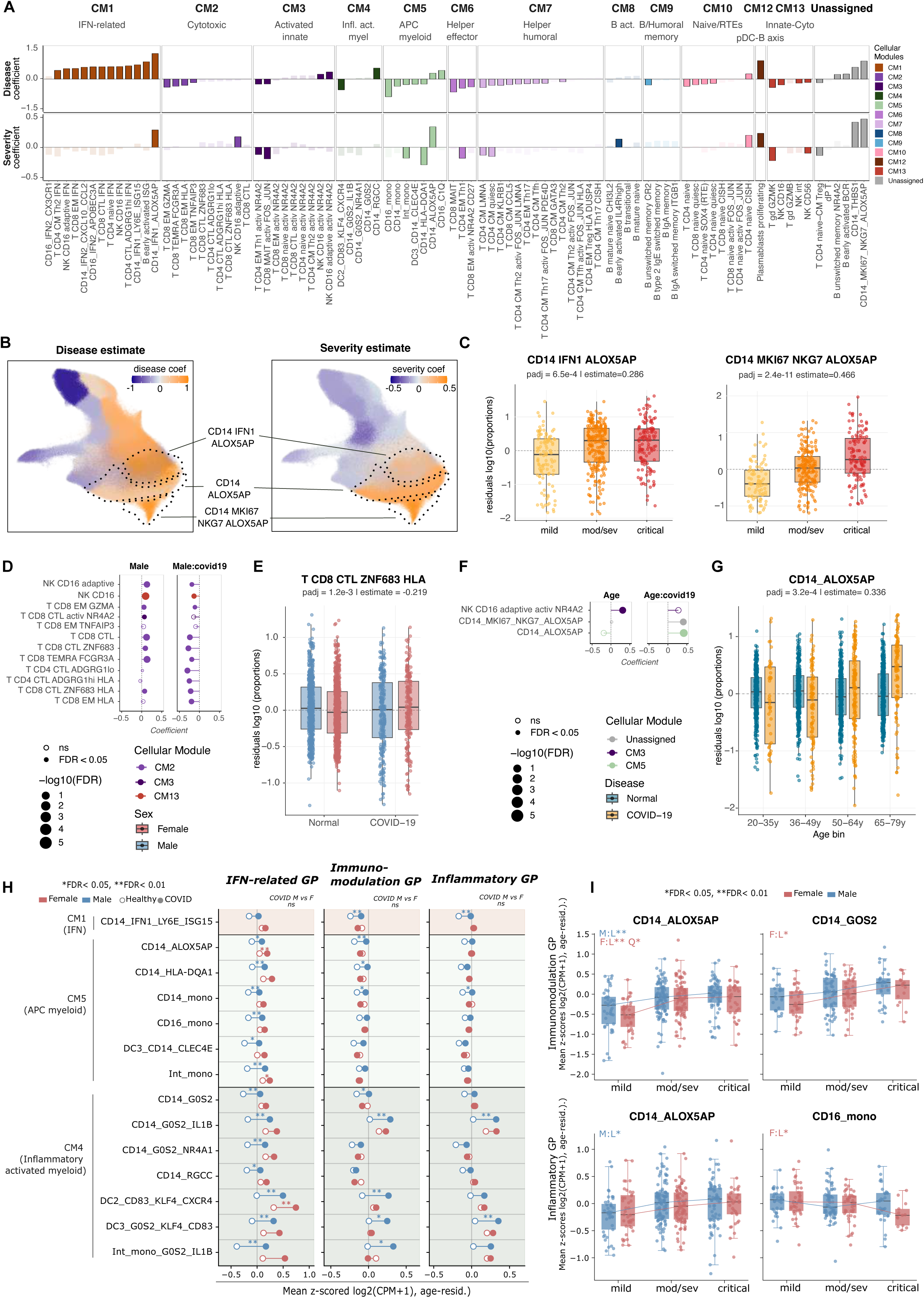
Disease, age, and sex-associated variation in immune cell-subset abundance and gene program activity during COVID-19. (**A**) Bar charts depicting per-cell subset coefficients of (top) the COVID-19 disease versus controls covariate (log_10_(proportion) ∼ age_bin + sex + COVID-19_or_control + study) and (bottom) the disease severity covariate (log_10_(proportion) ∼ age_bin + sex + disease_severity + study; disease severity modelled as ordered factors: mild < moderate/severe < critical). Bars are grouped and colored by the co-abundance cellular module (CM) membership of the respective cell subsets; solid outlines for bars indicate FDR < 0.05. (**B**) Gene expression UMAP of all monocytes and cDC2 cells, colored by the per-cell subset coefficients of (left) the COVID-19 disease versus controls covariate and (right) the disease severity covariate. Granular MS1 cell subsets are delineated by dotted lines. (**C**) Boxplots depicting residuals of log_10_(proportion) of (left) CD14_IFN1_ALOX5AP and (right) CD14_MKI67_NKG7_ALOX5AP per donor, grouped by disease severity. Residuals were computed by regressing the effects of study, age (binned), and sex. (**D**) Lollipop plots depicting coefficients of the male sex covariate and the interaction term between male sex and moderate, severe, or critical COVID-19 status (log_10_(proportion) ∼ age_bin + sex + moderate_severe_critical_COVID-19_or_control + sex:moderate_severe_critical_COVID-19_or_control + age_bin:moderate_severe_critical_COVID-19_or_control + study). (**E**) Boxplots depicting residuals of log_10_(proportion) of T_CD8_CTL_ZNF683_HLA per donor, grouped by moderate, severe, or critical COVID-19 status versus controls, as well as female/male sex. Residuals were computed by regressing the effects of study and age (binned). (**F**) Lollipop plots depicting coefficients of the age bin covariate and the interaction term between age bin and moderate, severe, or critical COVID-19 status. (**G**) Boxplots depicting residuals of log_10_(proportion) of CD14_ALOX5AP per donor, grouped by moderate, severe, or critical COVID-19 status versus controls, as well as age bin. Residuals were computed by regressing the effects of study and sex. Proportions for each cell subset per donor were computed out of all PBMCs per donor. (**H**) Lollipop plots of IFN-related, Immunomodulation, and Inflammatory GP scores across 14 myeloid cell subsets, grouped by CM (CM1: IFN monocyte; CM5: APC myeloid; CM4: Inflammatory activated myeloid). GP scores are calculated per sample as the mean of within-study z-scored log₂CPM values across GP genes (≥7 genes/GP), then residualised for age by OLS. Open circles show mean scores for healthy donors, filled circles for COVID-19 patients, for each cell subset and sex (blue: male; red: female); connecting lines indicate the direction and magnitude of change. Stars above a line denote significance of the COVID-19-vs-healthy comparison within the corresponding sex, while COVID-19 only comparison between sexes was not significant (NS). All three contrasts per cell subset (female COVID vs. healthy; male COVID vs. healthy; female vs. male within COVID) are OLS-tested with age as covariate and BH-corrected across all three contrasts for all GPs, per CM. *adj. p < 0.05, **adj. p < 0.01 (**I**) COVID-19 severity-dependent **immunomodulation** (top row) and **inflammatory** (bottom row) GP scores, illustrating male and female severity trajectories. Boxplots show age-residualised scores by sex (blue: male; red: female) across severity categories (mild, moderate/severe, critical; ≥3 samples per sex × severity required). Boxes span the interquartile range with a median bar; individual samples are overlaid as jittered points, with lines connecting group medians. BH correction is applied independently within each of the four contrasts (male linear [L], male quadratic [Q], female linear, female quadratic) across CM. Panel annotations: M/F = sex, L/Q = contrast type; significance thresholds as in (H).

In contrast to these broad changes in cell abundance between healthy individuals and those with COVID-19, relatively few cell subsets showed progressive association with disease severity. Among the subsets most strongly associated with disease severity were MS1-like monocytes, which have previously been linked to COVID-19 severity and sepsis^4,10,12,57^ and are marked by expression of genes including *ALOX5AP*, *CLU*, and *RETN*. HARP resolved the previously described MS1 population into three subsets: CD14_ALOX5AP, CD14_IFN1_ALOX5AP, and CD14_MKI67_NKG7_ALOX5AP. The three MS1-like subsets differed markedly in their association with disease severity (Fig. 5A-C), with the proliferative CD14_MKI67_NKG7_ALOX5AP population showing the strongest association with disease severity than other MS1-like subsets, demonstrating that fine-grained annotation can distinguish clinically informative heterogeneity within previously defined disease-associated cell populations.

Because age and sex are key determinants of COVID-19 outcome, with highest mortality observed among older men^58^, we next examined whether disease-associated changes in immune-cell abundance differed across these demographic variables. Although cytotoxic lymphocyte populations within CM2 increased with age in healthy individuals (Fig. 4A), impaired cytotoxic responses have been associated with severe COVID-19^59,60^. Among individuals with moderate-to-severe or critical COVID-19, cytotoxic CM2 populations were reduced significantly more in males than females (Fig. 5D). For example, the newly resolved T_CD8_CTL_ZNF683_HLA subset exhibited a significant interaction between COVID-19 and sex, with the direction of the sex-associated difference reversing between healthy controls and individuals with COVID-19 (Fig. 5E).

COVID-19 associated remodeling likewise differed with age (Fig. 5F; Supplementary Fig. 10B). For example, CD14_ALOX5AP monocytes were largely unchanged across the age range among healthy controls, but were markedly expanded in older individuals with moderate, severe, or critical COVID-19 (Fig. 5G). In sum, these findings demonstrate that HARP not only recapitulates established features of COVID-19 immune dysregulation but also resolves clinically relevant heterogeneity within disease-associated immune populations and reveals age- and sex-dependent differences that are obscured at lower cell annotation resolution.

### Sex-divergent transcriptional responses to COVID-19 across myeloid and lymphoid compartments

In healthy individuals, nine immune-related gene programs (GPs), including *IFN–related* response and *NF-kB signaling* programs, showed higher expression in females than males (Fig. 4E). We therefore sought to determine whether COVID-19 altered these programs differently by sex. The same nine GPs were scored across pseudobulk profiles from the 14 myeloid and 35 lymphoid cell subsets used to derive the programs, in both individuals with COVID-19 and healthy controls. GP scores were then compared between healthy controls and patients with COVID-19 within each sex, and between females and males with COVID-19 (Fig. 5H, I, Supplementary Fig. 10C-F, Supplementary Fig. 11). Across multiple GPs, males exhibited more substantial transcriptional responses to COVID-19, because their healthy baseline expression was lower.

This distinction between sexes was evident for the *IFN-related* GP across the APC myeloid (CM5) and the inflammatory activated myeloid (CM4) cellular modules (Fig. 4E). Males demonstrated lower baseline *IFN-related* GP expression and greater induction in response to COVID-19 than females, yet expression in males remained consistently lower during disease (Fig. 5H). A similar male-biased induction pattern was observed for the *immunomodulation and inflammatory* GPs, which were increased with COVID-19 across cell subsets spanning CM1, CM4, and CM5 in males but did not significantly change in females, consistent with their higher baseline expression in healthy individuals (Fig. 5H). The *hypoxia/metabolic stress* and *stress/apoptosis* GPs followed a similar male-biased pattern, showing greater COVID-19 associated induction across myeloid subsets spanning CM1, CM4, and CM5 in males (Supplementary Fig. 10C). These findings suggest greater metabolic stress- and apoptosis-associated transcriptional activity in the male myeloid compartment during COVID-19. By contrast, the *cytotoxic* GP showed significantly higher expression in the DC2_CD83_KLF4_CXCR4 subset for females with COVID-19 (Supplementary Fig. 10C).

We next investigated whether these GPs varied with COVID-19 severity in a sex-dependent manner. The *immunomodulation* GP increased with severity in both sexes, with the strongest associations observed in CD14_ALOX5AP and CD14_G0S2 monocytes. The *inflammatory* GP likewise increased with severity in CD14_ALOX5AP cells in both sexes but decreased specifically in CD16 monocytes in females. In contrast, the *IFN-response* GP did not show a significant association with disease severity, suggesting that IFN activation is associated primarily with COVID-19 itself rather than with progressive disease severity (Fig. 5I; Supplementary Fig. 10D-F).

The lymphoid compartment showed both shared and distinct sex-associated responses. The strongest COVID-19-associated changes were observed within the helper/humoral (CM7) and the naive/RTE (CM10) compartments and involved the *IFN-related*, *activation*, *immunomodulation*, *hypoxia/metabolic stress*, *adhesion/migration*, *stress/apoptosis*, and *NF-kB signaling* GPs (Supplementary Fig. 11). As in the myeloid compartment, these programs generally showed greater disease-associated induction in males. The *IFN-related* GP remained the dominant sex-associated signal, although both sexes showed increased expression in COVID-19. In contrast, among individuals with COVID-19, the *cytotoxic* GP was consistently higher in females across all T- and NK-cell related CMs, including CM2 and CM7.

In sum, these analyses demonstrate that sex-specific responses to COVID-19 are organized through distinct multicellular gene programs, with inflammatory and immunomodulatory responses predominating in male myeloid compartments and cytotoxic programs preferentially maintained in female lymphoid compartments.

### scTiger enables scalable label transfer of high-resolution PBMC annotations to external datasets

To facilitate broad adoption of HARP, we developed *scTiger* (*s*ingle *c*ell *T*ranscriptional *I*mmunophenotyping using *G*radient boosting and *E*xpert *R*easoning), a hierarchical label-transfer framework designed to map the high-resolution HARP cell annotation to previously unseen PBMC datasets (Methods). *scTiger* can distinguish transcriptionally similar immune cell subsets while remaining computationally efficient and scalable to millions of cells. *scTiger* combines hierarchical lineage classification with lineage-specific metric learning, enabling accurate annotation of fine-grained immune cell subsets while preserving biologically meaningful relationships between closely related populations (Fig. 6A).

**Figure 6:**
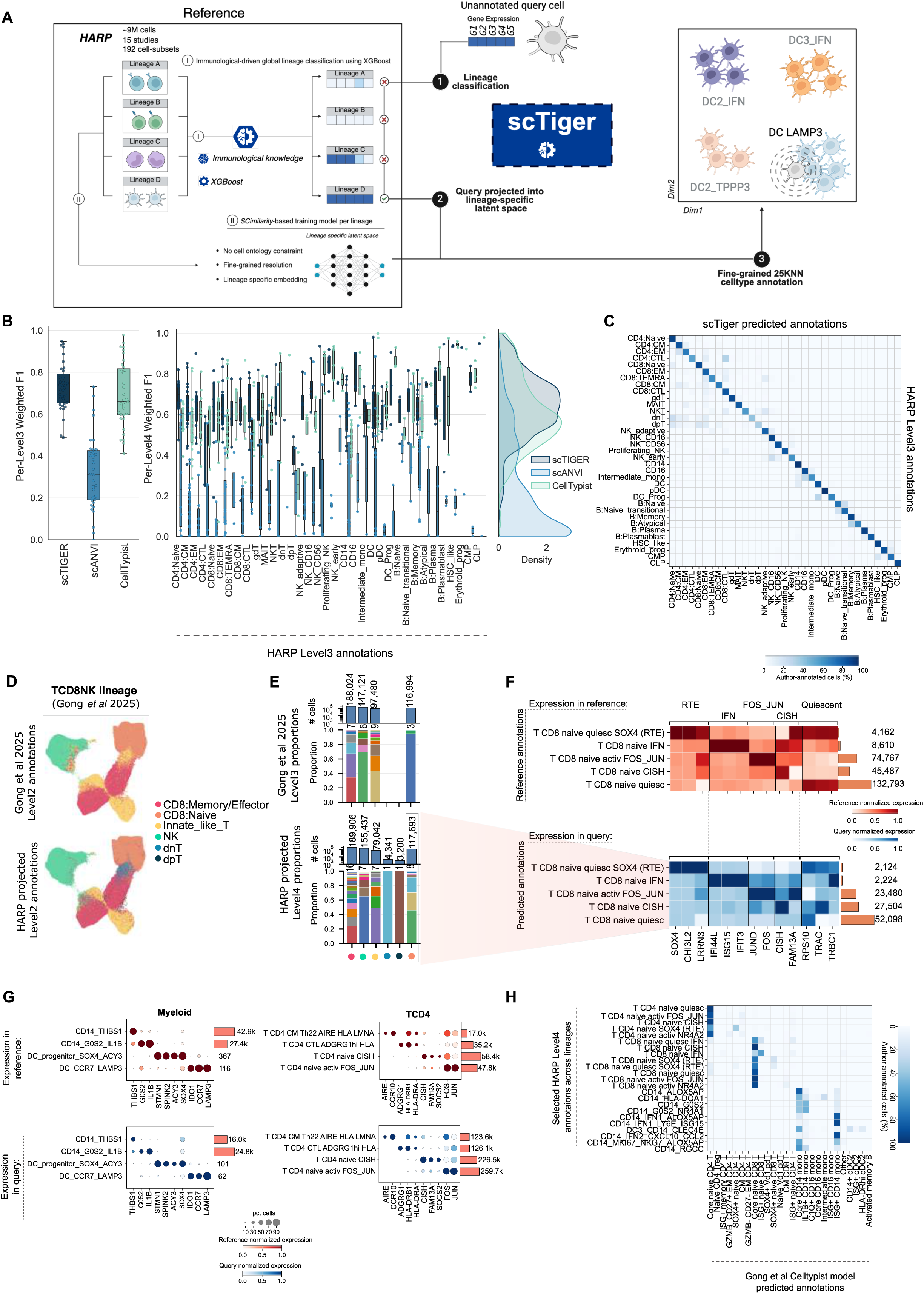
scTiger enables hierarchical, lineage-aware annotation of immune cells at fine-grained resolution. **(A)** Overview of the scTiger framework. HARP is first used to train a hierarchy of XGBoost classifiers for major immune lineage assignment (CD4 T cells, CD8 T cells, NK cells, B cells, myeloid cells, and HSPCs). These lineage annotations are then used to train independent lineage-specific neural-network encoders, generating reference latent embeddings for each lineage. During annotation, query cells are first assigned to a lineage using the trained XGBoost hierarchy before being projected onto the corresponding lineage-specific latent space. Fine-grained cell-subset labels are assigned by majority voting amongst the **25** nearest reference neighbors within each lineage-specific embedding. **(B)** Performance comparison of scTiger, CellTypist, and scANVI on held-out HARP unseen batches. Left: boxplots of weighted F1 scores for Level 3 immune lineages. Each dot represents a Level 3 category. Center: boxplots of per-cell-subset weighted F1 scores for Level 4 cell subsets grouped by their Level 3 annotation. Each dot represents a Level 4 cell subset. Right: density distributions summarizing classifier performance across all evaluated cell subsets in the boxplots at the center. **(C)** Confusion matrix comparing scTiger predicted annotations and original HARP Level 3 annotations. Color intensity represents the percentage of predicted cells correctly assigned to their corresponding original annotation. **(D-G)** Application of scTiger to the Gong **et al.** PBMC dataset^6^. **(D)** UMAP visualizations of original author-provided Level 2 annotations (top) and scTiger-transferred HARP annotations (bottom) are shown for the CD8 T and NK population. **(E)** Stacked bar plots summarizing the composition of the Gong **et al.** author-provided Level 3 annotations (top) and the scTiger-transferred HARP Level 4 annotations (bottom) within each original Gong **et al.** Level 2 population. Grey box highlights the CD8:Naive population, originally annotated as a single cell type by Gong **et al.**, which is resolved into multiple transcriptionally distinct naïve CD8 T-cell subsets following HARP reference mapping. **(F)** Heatmaps comparing expression profiles of the newly resolved cell subsets in the original Level 2 CD8:Naive population from the Gong **et al.** query dataset (bottom) and the corresponding cell subsets in HARP (top). Only cell subsets representing >1% of the original Gong **et al.** Level 2 CD8:Naive population are displayed. Horizontal bars indicate numbers of each cell subset per dataset. **(G)** Dotplots showing marker gene expression in HARP (top) and in the Gong **et al.** dataset for the scTiger-transferred HARP annotations (bottom) for selected rare and fine-grained myeloid and dendritic cell (left) and CD4 T-cell (right) populations. Dot size indicates the fraction of expressing cells and color intensity denotes mean expression. Horizontal bars indicate numbers of each cell subset per dataset. **(H)** Confusion matrix comparing HARP Level 4 annotations with cell-type annotation predictions generated using the pretrained Gong **et al.** CellTypist model.

To create the reference embedding space, independent neural-network encoders were trained for each major immune lineage using a fine-tuned version of *SCimilarity*^61^ with HARP serving as the reference atlas. Unlike the original *SCimilarity*, which learns a single global embedding using an encoder-decoder architecture constrained by Cell Ontology (CL) relationships, *scTiger* employs lineage-specific encoder-only models trained directly on the HARP annotation hierarchy. Here, each encoder is optimized using a joint objective combining supervised triplet loss, cross-entropy classification loss, and center loss, encouraging cells sharing the same fine-grained annotation to cluster together while improving class discrimination and reducing intra-class variability. Importantly, because CL relationships incompletely capture the transcriptional programs distinguishing closely related immune subsets within individual lineages, *scTiger* removes ontology-guided triplet selection and instead learns directly from the HARP annotation hierarchy. This preserves separation among transcriptionally similar and rare immune cell populations that would otherwise be compressed together into common embeddings. To improve representation of rare immune subsets, lineage-specific training also incorporates oversampling of low-frequency populations. Following training, latent embeddings for every HARP reference cell are precomputed and stored together with their annotations, enabling rapid nearest-neighbor label transfer.

Label transfer proceeds through two hierarchical stages. First, query cells are assigned to one of six major immune lineages (CD4 T cells, CD8 T cells, NK cells, B cells, myeloid cells, or hematopoietic progenitor) using a hierarchy of XGBoost classifiers trained on canonical lineage marker gene expression. Second, each query cell is projected into the corresponding lineage-specific reference latent embedding, where its identity is assigned by majority voting among the 25 nearest reference neighbors. Annotation can be performed at any level of the HARP annotation hierarchy (levels 1-4), allowing users to tailor annotation granularity against confidence depending on the biological question being addressed. Restricting nearest-neighbor searches to lineage-specific embeddings substantially reduces computational complexity while improving discrimination amongst transcriptionally similar immune cell subsets. Runtime and memory benchmarking demonstrated competitive scaling relative to existing approaches, including *CellTypist*^62^ and *scANVI*^14^, enabling efficient annotation of datasets containing up to one million cells (Supplementary Fig. 12C, D).

We first evaluated *scTiger* at the Level 3 annotation hierarchy using held-out HARP batches (20% of the dataset), strictly withheld from training. At this level, which provides sufficient biological resolution for many downstream single-cell analysis, *scTiger* achieved a weighted F1 score of 0.94. The method reliably distinguished major immune cell subsets (e.g., naive and memory lymphocytes), while also resolving rare populations such as double-negative or double-positive T cells and HSPC subsets (e.g., Common Myeloid Progenitor [CMP] and Common Lymphoid Progenitor [CLP]; Fig. 6C).

We next evaluated performance across the complete set of 192 fine-grained Level 4 cell subsets. Despite the substantially greater annotation complexity, *scTiger* continued to outperform existing approaches, achieving a weighted F1 score of 0.66 compared with 0.55 for CellTypist and 0.40 for scANVI (Fig. 6B; Supplementary Fig. 13A, B). Importantly, this average performance masked considerable variation across individual cell subsets. Many level 4 populations were recovered with high recall (>0.80), including Prolif_NK_activ_FOS_JUN, T_CD4_EM_Treg_HLA_DUSP4, CD14_IFN1_MT1G, DC5_IL3RA, and HSC_MPP_CLP. In contrast, most populations with recall below 0.50 reflected confusion between closely related transcriptionally defined subsets rather than misclassification across major subsets. This was particularly evident for highly transcriptionally similar CD4 cytotoxic *ADGRG1* subsets and closely related pDC cell states.

These results demonstrate that *scTiger* provides highly accurate annotation at Level 3 intermediate hierarchical resolution while maintaining substantially greater resolution than existing methods at the finest annotation level. Importantly, the remaining Level 4 ambiguities predominantly occur among transcriptionally adjacent immune subsets, suggesting that they reflect the challenge of distinguishing closely related transcriptional programs of cell states existing on a biological continuum within otherwise well-defined immune subsets, rather than failures to recover canonical immune identities. For such fine-grained annotations existing on a cellular continuum, we envision *scTiger* to be used as a fast tool for initial label transfer, where users can quickly annotate their clusters and then verify against the provided marker genes to confirm or adjust any transcriptionally similar cell subtypes.

To determine whether *scTiger* generalized beyond the HARP training data, we applied it to two large independent published scRNAseq PBMC datasets comprising approximately 18 million cells: (1) Gong *et al.*^6^, (labeled as *Gong*; ∼16M PBMCs) and (2) Terekhova *et al.*^48^ (labeled as *Terekhova*; ∼2M PBMCs). At coarse annotation levels, *scTiger* showed strong concordance with the published annotations across all major lineages (Fig. 6D). At the finer annotation level (level 4), however, HARP resolved substantially greater immune cell subset diversity (Supplementary Fig. 12). Within the CD8/NK lineage, for example, HARP distinguished 75 immune subsets (Fig. 6E - bottom panel) compared with 26 reported by Gong *et al.* (Fig. 6D,E - top panel). This difference in cell subsets recovered may partly reflect the absence of COVID-19 or other disease states in the Gong *et al.* dataset, as well as potential undersampling of certain populations. Importantly, when the five HARP-defined naive CD8 T cell subsets were projected onto the *Gong* dataset, they retained the same defining marker gene expression patterns used for their original annotation in HARP (Fig. 6F), demonstrating that these populations represent reproducible biological subsets rather than dataset-specific transcriptional clusters and that *scTiger* can recover this additional biological resolution in existing PBMC datasets.

We next asked whether *scTiger* could accurately recover very rare immune populations. Within the myeloid compartment, two of the rarest HARP subsets – DC_progenitor_SOX4_ACY3 (387 cells; 0.004% of PBMCs) and DC_CCR7_LAMP3 (116 cells; 0.001% of PBMCs) – were identified in the *Gong* dataset at comparable frequencies while preserving their transcriptional marker gene signatures that originally defined those subsets in HARP (Fig. 6G - left panels). Similarly, *scTiger* recovered rare CD4 T cell populations, including Th22 cells (T_CD4_CM_Th22_AIRE_HLA_LMNA; 17,000 cells, 0.2% of PBMCs) and *ADGRG1*^hi^ CD4 T cells (7,106 cells, T_CD4_CTL_ADGRG1^hi^_HLA; 0.4% of PBMCs), together with their characteristic marker genes (Fig. 6G - right panels). Integrated-gradient feature-attribution scores further demonstrated that *scTiger* learns biologically interpretable representations rather than opaque classification rules, with subset-defining marker genes consistently ranking as features with the highest attribution scores and emerging among the strongest contributors to classification decisions. For example, *CCR7* and *LAMP3* emerged among the highest-ranking features contributing to classification of DC_CCR7_LAMP3 cells (Supplementary Fig. 13E–G);

Finally, we assessed whether these HARP annotations could be recovered using existing PBMC annotation tools. Annotation of HARP using the CellTypist model provided by Gong *et al.*^6^ (https://apps.allenimmunology.org/aifi/resources/imm-health-atlas/) collapsed multiple well-defined HARP immune subsets into a small number of broad categories, particularly within the naive CD4 and CD8 T cell compartments (Fig. 6H). Multiple transcriptionally distinct naive T-cell subsets were merged into common naive T-cell labels despite retaining distinct molecular signatures. These results illustrate that the expanded HARP annotation captures biologically meaningful cell subset diversity that is not recovered by existing PBMC annotation tools. Together, these analyses demonstrate that *scTiger* accurately transfers the high-resolution HARP annotation across independent datasets while preserving biologically meaningful immune cell diversity, including rare immune populations and fine-grained immune subsets that are frequently collapsed together by existing automated label-transfer annotation methods.

## Discussion

Here we present HARP, an HCA peripheral blood reference atlas integrating 15 scRNAseq studies from four continents, comprising a core reference of ∼8.4 million cells and 192 immune cell subsets. HARP complements existing large-scale PBMC atlases^6,49^ by substantially increasing the number of donors, geographic representation, and annotation resolution through harmonized multi-center integration. Together with scTiger, a hierarchical metric-learning framework for label transfer, HARP provides a scalable and transferable reference for high-resolution immune cell annotation. Using scTiger, we successfully projected and annotated an additional ∼18 million PBMCs from independent studies, more than tripling the number of cells analyzed within a unified reference framework and demonstrating the scalability and generalizability of HARP as a community reference atlas.

The rapidly expanding volume of single-cell data generated across tissues and disease contexts presents unprecedented opportunities for integrative analyses and the development of foundation models capable of predicting clinically relevant outcomes^63,64^. However, our study also highlights important challenges associated with large-scale, unsupervised data integration. We found that many published datasets lack canonical marker genes of key immune cell populations or contain contamination from low-quality cells, introducing technical artefacts that obscure biological signals and propagate into downstream analyses. To address these issues, we present a systematic data integration framework, supported by qualitative and objective quantitative metrics for evaluating data filtering, dimensionality reduction, and batch correction during atlas assembly.

A second major challenge is the absence of standardized cellular annotation. Although blood is among the best-characterized human tissues, substantial variation in nomenclature persists across studies, limiting reproducibility and cross-study comparison. The hierarchical annotation presented here was developed collaboratively by members of the HCA Immune Bionetwork members across four centers and is grounded in established immunological knowledge. Rather than representing a final fixed cellular taxonomy, we envision this annotation as a living community resource that will continue to evolve through collaborative review and community engagement via the HCA Cell Annotation Platform (CAP; https://celltype.info/), through which the community is invited to contribute feedback as we work together towards a consensus reference annotation for human PBMCs.

Beyond resolving fine-grained immune subsets, HARP revealed higher-order multicellular organization through coordinated Cellular Modules (CMs) and shared transcriptional gene programs. These complementary levels of organization expose coordinated immune cell relationships that are not apparent when individual cell subsets are considered in isolation, providing a systems-level framework for understanding immune remodeling across aging, sex, and disease.

This systems-level framework revealed conserved transcriptional programs that recur across immune cell types, including interferon and TNF-response pathways, and coordinated CMs that remodeled in response to demographic and disease-associated factors. These CMs recapitulate established features of immune ageing, including declining naive and recent thymic emigrant populations, together with expansion of helper, humoral, and cytotoxic compartments, consistent with thymic involution and cumulative antigen exposure^48,49,65,66^. One particularly notable finding was a previously unrecognized age-associated module characterized by high expression of *NR4A2*. As *NR4A2* is induced by persistent TCR signaling and regulates chronic antigen adaptation and T-cell exhaustion, expansion of *NR4A2*^Hi^ lymphocyte subsets is consistent with the progressive accumulation with age of chronically stimulated, antigen-experienced immune cells^50,51^.

Sex emerged as another major determinant of immune organization, influencing both immune cell abundance and coordinated transcriptional programs. For example, males consistently exhibited higher cellular abundance of Tfh, Th2, Th17, Th22, cytotoxic T-cell, and CD16^Hi^ NK-cell populations, whereas females showed higher expression of interferon-response, NF-kB signaling, and cytotoxic gene programs. These findings extend previous reports of sex-specific interferon responses^67,68^ by resolving both the underlying transcriptional programs and their cellular context. Enhanced basal interferon signaling in females is consistent with the effects of X chromosome-encoded immune regulators and estrogen on type I/II interferon production^69–72^ and may contribute to the increased susceptibility of women to autoimmune disease^72–74^.

Among the most reproducible sex-associated transcriptional differences, we identified coordinated regulation of prostaglandin pathway genes. Female immune cells showed higher expression of *PTGS2* together with *PTGER2* and *PTGER4*, consistent with enhanced capacity for inducible PGE₂ synthesis and signaling. In contrast, males preferentially expressed *PTGDS*, suggesting relatively greater engagement of the PGD₂ pathway, which promotes type 2 immune responses through CRTH2-dependent recruitment of Th2 cells, eosinophils, and basophils^75,76^. Although these observations require validation in additional tissue contexts where prostaglandins exert their primary effects, they suggest previously unappreciated sex-specific organization of prostaglandin signaling in human immunity.

Because COVID-19 has been extensively profiled by single-cell genomics, it provided an ideal benchmark for evaluating whether HARP could both recapitulate established features of immune dysregulation and reveal additional biological insights by leveraging the higher-resolution annotation. Our fine-grained annotation enabled us to distinguish widespread immune remodeling associated with COVID-19 disease versus controls from a relatively small number of immune subsets that tracked progressive disease severity. For example, coordinated remodeling of interferon-responsive cell subsets (CM1) reflected the broad immune response to infection, whereas progression to severe disease was associated with changes in only a limited number of immune subsets. At the transcriptional level, males generally mounted larger inducible responses relative to their healthy baseline; however, these responses frequently remained below the absolute expression levels observed in females, consistent with previous hypotheses^77^. Together, these findings suggest that both baseline immune organization and inducible transcriptional capacity contribute to sex-specific immune responses during infection.

We also refined the characterization of monocyte-state 1 (MS1) monocytes, an immunosuppressive population associated with sepsis and severe COVID-19^4,57^. HARP resolved this population into multiple transcriptionally distinct subpopulations, including a cycling subset that showed the strongest association with COVID-19 disease severity. This finer resolution suggests that the biological function previously attributed to MS1 monocytes may arise from distinct underlying subpopulations rather than a single cellular state.

Finally, we developed *scTiger* to extend the utility of HARP beyond the reference atlas itself by enabling robust annotation transfer to independent datasets while preserving biologically meaningful fine-grained immune subset annotations. *scTiger* consistently outperformed existing methods when transferring the 192-cell-subset HARP hierarchy to independent datasets while remaining scalable to millions of cells. Successful projection onto two independent PBMC atlases demonstrated that both common and rare immune cell subsets, including DC_progenitor_SOX4_ACY3 and DC_CCR7_LAMP3 (0.001-0.004% of PBMCs), can be reproducibly identified across studies, establishing HARP and its accompanying annotation framework as a scalable community reference for harmonizing future PBMC datasets.

Our study has several limitations. We prioritized robust cell-subset annotation through construction of a shared reference embedding rather than modelling patient-level variation. Future integration approaches designed to preserve donor-specific heterogeneity across even larger datasets may provide complementary biological insights^78,79^. In addition, our analyses are cross-sectional, restricted to circulating PBMCs rather than tissue-resident immune cells, and encompass a limited range of diseases, underscoring the need for longitudinal studies and broader disease representation in future versions of the atlas.

In summary, HARP provides a harmonized, high-resolution, and transferable reference atlas of human peripheral immunity. Beyond serving as a community resource, HARP demonstrates that combining biologically informed data integration, community-driven annotation, multicellular organization through cellular modules and gene programs, and scalable label transfer can enable the discovery of coordinated immune programs associated with age, sex, and disease. We anticipate that HARP and *scTiger* will provide a foundation for future studies of human immune variation and a framework for community-scale reference atlas construction across the Human Cell Atlas and beyond.

## Data and code availability

Data and code availability will be released upon publication. Processed lineage objects will be available via an interactive website and on the HCA Data Portal. Code will be made available via GitHub and Zenodo. Annotation information will be available via the Cell Annotation Platform.

## Methods

### Data sources

#### De novo generated scRNAseq data

For the unpublished study (*HCAunpublished1*), peripheral blood was collected under the Tasmania Health and Medical Human Research Ethics Committee (H0012902)^8^. Peripheral blood was collected into 8 mL CPT™ tubes (FICOLL™/sodium heparin; BD 362753) and processed within 4 hours of collection. Cells were washed, resuspended in freezing medium and stored in liquid nitrogen. CITE-seq antibodies were centrifuged and a staining mastermix was prepared by combining CITE-seq antibodies and isotype controls (1:100). Thawed PBMCs were dispensed into a 96-well plate, washed, and resuspended in mastermix. Samples were incubated and cell concentration and viability were assessed by trypan blue exclusion. Stained single-cell suspensions were pooled and loaded onto the Chromium Single Cell Chip A (10x Genomics) targeting capture of 20,000 cells per well, and partitioned and barcoded using the Chromium Controller with the Single Cell 3a Library and Gel Bead Kit (PN-120237). GEM generation, barcoding, cDNA amplification, and library construction were performed according to the 10x Genomics Chromium User Guide. Gene-expression and antibody-derived tag (ADT) libraries were generated with unique sample indices, multiplexed, and sequenced on an Illumina NovaSeq 6000 across 2 × 150 cycle flow cells (26 bp Read 1, 8 bp Index, 98 bp Read 2). As the 35-antibody CITEseq panel used for this dataset did not fully overlap with the 192-antibody panel used in the COMBAT^12^, Stephenson^9^, and Slowikovski^10^ datasets, only the gene expression data were used for the analyses presented in this study.

#### Published scRNAseq data

The original single-cell RNA-seq data count matrices from source studies used in HARP is available via the original studies and the remapped count matrices will be available via the HCA Data Portal.

### scRNAseq analysis

#### UMI quantification

Raw FASTQ files were downloaded and processed using scripts from https://github.com/cellgeni/reprocess_public_10x. Briefly, a series of metadata for each dataset was collected using the GEO soft family file. Following this, ENA web API was used to obtain the information about the format in which raw data is available for every run (SRR/ERR), as well as to infer the sample-to-run relationships. Raw read files were then downloaded in one of the three formats: (1) SRA read archive; (2) submitter-provided 10x BAM files; (3) gzipped paired-end FASTQ files. SRA archives were converted to fastq using fastq-dump utility from NCBI SRA tools v2.11.0 using “-F -- split-files” options. BAM files were converted to fastq using 10X bamtofastq utility v1.3.2. Following this, raw sequencing reads were mapped and quantified using the STARsolo algorithm. STAR version 2.7.10a_alpha_220818 compiled from source files with the “-msse4.2” flag was used for all samples. Wrapper scripts documented in https://github.com/cellgeni/STARsolo/ were used to auto-detect 10x kit versions, appropriate whitelists, and other relevant sample characteristics. Human reference genome and annotation exactly matching Cell Ranger 2020-A was prepared as described by 10x Genomics: https://support.10xgenomics.com/single-cell-gene-expression/software/release-notes/build#header. The STARsolo command was optimized to generate the results maximally similar to the Cell Ranger v6 software from 10x Genomics. Namely, “--soloUMIdedup 1MM_CR –soloCBmatchWLtype 1MM_multi_Nbase_pseudocounts --soloUMIfiltering MultiGeneUMI_CR --clipAdapterType CellRanger4 -- outFilterScoreMin 30” were used to specify UMI collapsing, barcode collapsing, and read clipping algorithms. For paired-end 5’ 10x samples, options: `“-- soloBarcodeMate 1 --clip5pNbases 39 0”` were used to clip the adapter and perform paired-end alignment. For cell filtering, the EmptyDrops algorithm employed in Cell Ranger v4 and above was invoked using `“--soloCellFilter EmptyDrops_CR”` options. Options `“--soloFeatures Gene GeneFull Velocyto”` were used to generate both exon-only and full length (pre-mRNA) gene counts, as well as RNA velocity output matrices.

### Metadata harmonization

#### Tier 1 metadata harmonization

Cell-level tier 1 metadata comprising technical information required for integration (as defined by HCA^81^) was collected from the CZI CELLxGENE^82^ data objects^3,7–12,18–23^, from the study publications’ supplementary information^20^, as well as directly from the study authors^10^. All metadata was matched by barcode and mapped on a per-study basis to the remapped matrices. Sex metadata was harmonized using the original sex information where available, with missing values inferred using the chromosome Y non-pseudoautosomal region (PAR)-to-PAR expression^7^. Missing values were imputed at the donor level when unambiguous, and prediction-guided updates were restricted to high-confidence assignments.

#### Tier 2 metadata harmonization

The harmonization of tier 2 metadata, which comprises biological information required for downstream analysis^81^, was performed in two phases. For studies where tier 2 metadata fields were not publicly available (sex, age, disease status), metadata was collected manually from study authors. For cases where development stage (i.e., age) annotations were available from CZ CELLxGENE^82^ published studies, development-stage ontology terms and free-text age fields were normalized to a to numeric midpoints (*age_midpoint*), which were either the actual age, if a numeric age was present, or the average of the age interval. All *age_midpoints* were aggregated into fixed age bins schemes (5-year, 10-year, and custom harmonized [0-1y, 1-19y, 20-35y, 36-49y, 50-64y, 65-79y, >80y] bins). For COVID-19, the original disease severity metadata were harmonized at the sample level according to the following criteria: *mild*: asymptomatic; viral RNA detected; symptomatic and independent; symptomatic and requiring assistance; hospitalized without oxygen therapy. *moderate/severe:* hospitalized and receiving oxygen by mask or nasal prongs; hospitalized and receiving non-invasive ventilation (NIV) or high-flow oxygen. *critical*: intubation and mechanical ventilation; mechanical ventilation or vasopressor support; mechanical ventilation with vasopressor support, dialysis, or extracorporeal membrane oxygenation (ECMO).

### Semi-automated, biologically stratified QC harmonization

Quality control was performed independently for each lineage within each study to account for lineage- and study-specific differences in transcriptional complexity and sequencing depth. First, we grouped cells into seven manually harmonized broad immune cell lineage (T+NK+ILC, B, plasma, monocyte, dendritic cells, neutrophil, and unassigned) based on Cell Ontology^83^ labels collected by CZI CELLxGENE^82^, which represent standardized mappings of the original author annotations. Secondly, we applied sctk scAutoQC^84^ to automatically generate density-based thresholds for each cell lineage per study. QC metrics used for scAutoQC’s Gaussian fitting step included total counts (n_counts), numbers of detected genes (n_genes) and fraction of mitochondrial reads (percent_mito). Diagnostic plots for QC metrics and automated thresholds were generated and stratified by author annotation, disease, and harmonized labels for a total of 105 separate QC subsets, which we inspected and adjusted manually for cases where the automated thresholds were too strict or permissive. Cells that did not pass the semi-automated QC thresholds were excluded from integration and downstream analyses.

### Author’s original cell label harmonization

In order to assess cell-type biological conservation during integration benchmarks, we further harmonized the original author cell annotations. Author-provided cell-type annotations were reconciled across studies using CellHint v1.0^17^, which aligns cell labels between datasets in a shared low-dimensional space. CellHint was run with the study identity as the batch variable and the original author annotation as the input label, using an scVI^30^ embedding as the alignment space. Resulting harmonized labels were validated by inspection of lineage-specific marker gene expression and grouped into immune cell lineages (i.e., CD4 T cells, CD8 T cells, NK cells, B and plasma cells, monocytes, dendritic cells). All author-provided annotations were manually curated into 32 cell-type categories, informed by CellHint’s alignment output. Labels that were too coarse (e.g. “B cell”, “PBMC”) were kept as unannotated. For these labels, and where author labels were missing or could not be harmonized (different resolution across datasets), we used CellTypist^62^ with the Healthy_COVID-19_PBMC model (https://www.celltypist.org/models) for cell type prediction. These annotations were solely used for assessing biological conservation through the ARI, NMI, and kBET scores^32^, and lineage definition during global vs lineage integration benchmarks, which happened prior to cell type annotation.

### Assessment of batch variables to use for integration

To determine the optimal batch variable for integration (e.g., sequencing library, study, sequencing chemistry), the merged data objects as well as per study datasets were pre-processed with the default Scanpy highly variable gene selection (with 2,000 top genes) and PCA with default parameters. These PCA results were used as input for the principal component regression (PCR) analysis. PCR scores were computed for each covariate to quantify the linear variance contribution (Σa wa·R²a, weighted by per-PC variance ratio), as previously described^32,85^. We assessed each score against an empirical null distribution from 1,000 permutations. Categorical covariates were permuted at the sample level, not the cell level, then broadcast back to cells, preserving within-sample composition, while numerical covariates were permuted per cell. This gave each covariate a z-score and an empirical P-value:

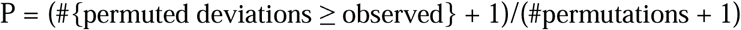

To identify confounding of technical and biological covariates, we used the Theil’s U^86^ statistic (uncertainty coefficient). For two variables X and Y, the uncertainty U is calculated as:

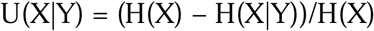

Where H is the entropy function. Theil’s U is non-symmetric, quantifying how much information in X is explained by Y, with larger values indicating stronger confounding of Y. We computed this statistic for all pairwise combinations of technical and biological covariates (Supplementary Fig. 3).

Cells were further aggregated into pseudobulk profiles by composite of sample, donor and library IDs (smallest entity of a potential batch), and dimensionality-reduced with PCA for visual analysis of technical and biological effects. All analyses were computed using the scAtlasTb framework^36^.

### Feature selection approach for optimized embeddings

Highly variable genes (HVGs) were selected using the highly_variable_genes function in Scanpy^87^, utilizing library_id, determined above as optimal batch variable, for batch-aware feature selection. To ensure the inclusion of disease-associated transcriptional programs while enabling robust batch correction, HVGs were selected as the union of genes in healthy and COVID-19 donors. Specifically, we calculated the top 3,000 HVGs independently for healthy and COVID-19 cohorts, and considered the union of both gene sets.

To avoid clustering driven by spurious signals that were not conducive to cell type annotation, we excluded specific genes from the final feature space selection. For example, we excluded B and T cell receptor (BCR, TCR) genes, which reflect clonal identity and are known to cause spurious clusters^88^. These included TCR (e.g. *TRAV, TRAJ, TRBV, TRBD, TRBJ*) and immunoglobulin heavy/light chain (e.g. *IGLV3-1, IGHJ1, IGHV1-2*) V(D)J genes to eliminate donor-specific signals driven by individual V(D)J gene usage. Specifically, we filtered out variable, diversity, and joining segment genes, while retaining constant-region genes. TCR variable genes encoding canonical invariant or semi-invariant T-cell receptors (TRAV1-2, TRDV1, TRDV2, TRGV9, TRAV10, TRBV25-1) were retained as exceptions given their utility as marker genes beyond clonotype identity. We further refined our feature set by removing sex-linked genes (e.g. *XIST, RPS4Y1, DDX3Y*)^7^ to minimise sex-specific artifacts.

### Quantitative benchmarking

We benchmarked scVI^30^, Harmony^29^, and scPoli^31^ at global and per lineage resolutions, as well as different preprocessing and hyperparameter choices, such as different feature selection approaches, count scaling or model architecture choices. We evaluated the different models with scib metrics^32^ for biological conservation (ARI, NMI, cLISI, cell_cycle conservation) and batch correction (PC regression, iLISI, kBET^85^, bras_batch^34^). Biological conservation metrics ARI, NMI and kBET were calculated against CellHint harmonized author-annotated clusters (see above).

We designed additional label-free biological conservation metrics to quantify cell type separation in the embedding, independent of any cluster-based annotations, using the Moran’s I^35,87^ score to assess autocorrelation of composite gene scores for canonical cell types and signatures using Scanpy’s gene scoring implementation^87,89^ in the integrated kNN graph. All metrics were computed per lineage, with lineage-specific cell type marker-based Moran’s I scores, including for example DC1 (*CLEC9A*, *XCR1*), γδ T cells (*TRGV9*, *TRDV2*), Tregs (*FOXP3*, *IKZF2*), and plasmablasts (*MKI67*, *CDK1*). In addition to cell type marker genes, we also used a canonical interferon signature^90^ to estimate preservation of disease effects.

All metrics are min-max scaled across all entries and aggregated to overall batch correction and bio conservation scores as previously described^32^. An overall integration performance score was computed as the weighted average of batch and bio scores:

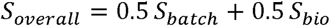

Benchmarks were set up sequentially to test different aspects at a time. We tested different feature selection strategies per lineage on a fixed Harmony model (Figure 2C) and a comparison of integration methods per lineage (Fig. 2F), as well as a comparison of lineage vs. global integration (Supplementary Fig. 1D). All benchmarks were implemented using the scAtlasTb framework^36^ and documented YAML files.

### Optimization of per-sublineage integrations

Following the preceding stages of optimization, we generated the final lineage embeddings. Using CellHint-harmonized author annotations, the full count matrix was partitioned into major lineage objects (Myeloid/dendritic cell [DC], B/Plasma, CD4 T, CD8 T/NK/Innate). We further stratified these objects by sublineage and performed integration on these objects independently to better enable novel/rare cell subset discovery.

Highly variable genes (HVG) selection was performed on the sublineage objects; we separately calculated the top 3000 HVGs for healthy and COVID-19 donors to ensure representation of disease-associated features. The union of the two sets was taken forward for dimensionality reduction.

Normalized gene expression counts for each sublineage object were scaled and underwent PCA for dimensionality reduction. The top 30 PCs were used for Monocyte/DC and B/Plasma sublineages and top 50 PCs were used for CD4 T and CD8 T/NK/Innate sublineages. Harmony (harmony-pytorch), our choice of integration strategy based on the optimization above, was then performed with library_id as batch key. Clustering was performed with the Leiden algorithm on the kNN graph of the PCA space, with multiple resolutions (1, 2, and 3) assessed. The kNN graph within each cluster was further subclustered using the Leiden algorithm at resolutions of 0.25 and 0.5 for rare cell subsets discovery. Suspected doublets, low-quality cells, and misclassified cells were filtered or reassigned to their correct sublineage and the above process repeated before arriving proceeding to final cell annotation.

### Consensus annotation

Cells were annotated at level 4 resolution across 11 sublineage objects: Myeloid/DC (Mono_DC2, DC_DC2); T/NK (T_CD4_naive, T_CD4_effector/nonnaive, T_CD8_naive, T_CD8_effector/nonnaive+ILC(NK), T/NK_proliferating); B/Plasma (B_naive, B_memory, Plasma_cells); and HSC (HSCs). For each object, 2-3 annotators from different research centers assigned cell annotations independently, which were collated into a shared spreadsheet and reconciled through iterative discussions. Marker genes were calculated using all-versus-all and one-versus-all comparison approaches using limma^91^ on sample pseudobulked gene expression of Leiden clusters; these marker genes were cross-checked at each cell annotation round via examination of feature plots and cluster gene expression. This process defined 192 consensus immune cell subsets across the 11 lineage-specific objects. Full annotations, marker gene rationale, annotation metadata, and the references used will be publicly available via the HCA Cell Annotation Platform (CAP).

### CITEseq analysis

Original CITE-seq data from three source studies^9,10,12^ were downloaded as separate AnnData objects and restricted to their shared feature set. Antibody names were harmonized manually to obtain a consistent mapping of 192 antibodies across the three datasets^9,10,12^, spanning 2,921,682 cells. ADT (protein) counts were split from GEX into a separate *obsm[’protein’]* matrix for each dataset, and the three datasets were then concatenated, with a *batch_dataset* metadata field tracking dataset of origin.

Mmochi v0.3.5^92^ was used to align protein peaks across the three datasets. Specifically, *mmc.utils.preprocess_adatas()* jointly preprocessed the protein-only objects by intersecting features and log-normalizing ADT counts (log(counts/total × 1000 + 1)). *mmc.landmark_register_adts()* was then run with *data_key=’protein’* to landmark-register the ADT distributions across batches, correcting for inter-study/inter-batch shifts in antibody staining intensity.

Cells were matched by barcode across the protein and gene expression modalities, yielding a final object of 2,705,361 cells with confirmed dual protein and gene expression. This object was used in the annotation process to delineate mixed clusters, confirm gene-label identity, and map CD45RA⁺ versus CD45RA⁻ T cell subsets.

### Downstream analysis

#### Data preparation for downstream analysis

All cells that passed QC were used for cell-type annotation and were aligned to a unified hierarchy (Level 1-4 annotation). For downstream analyses, we restricted to healthy control and COVID-19 cells with complete biological (age, sex, sample and donor ID) and technical (batch/library identifier) metadata. We focused exclusively on COVID-19 because using a single biological perturbation across multiple independent datasets enabled us to systematically evaluate and separate true biological variation from technical batch effects.

### Co-Abundance Cellular Modules (CMs)

#### Data Preprocessing and Quality Control Filtering

To eliminate technical noise and low-resolution data, filtering was applied to the cell-type annotation dataframe. Rare cell populations present in fewer than 300 samples were removed. At the sample level, samples which had fewer than 500 cells were excluded from this analysis. The remaining dataset was further filtered to retain only healthy and COVID-19 donors.

#### Compositional Transformation and Batch Correction

For the filtered dataset, a cell-type proportion matrix was generated by calculating the relative abundance of each cell type per sample. To stabilize variance, relative proportions *x* were log-transformed using the function *log(1+x)*. To explicitly control for study-specific batch effects, a within-study normalization was performed by calculating the Z-score of the log-transformed proportions independently for each individual study.

#### Co-Abundance Cellular Module (CM) Generation via Hierarchical Clustering

To identify robustly coordinated cell-type shifts, a pairwise Pearson correlation matrix was calculated across all cell-type Z-scores. To isolate tightly co-varying networks, a correlation filter was applied; cell types whose maximum absolute correlation coefficient with any other subset fell below a threshold of 0.3 were classified as non-correlating and excluded. The remaining filtered correlation matrix was converted into a distance matrix using a pairwise distance metric. Hierarchical clustering was then performed using the average-linkage method. The resulting dendrogram tree was cut into discrete, non-overlapping clusters. Clusters with <3 cell types were removed. The resultant clusters represented the 13 final co-abundance cellular modules (CMs).

#### Transcriptional Characterization of CMs

To characterize the molecular programs defining each CM, cell-type-specific, per-sample pseudobulk expression profiles were utilized. To extract gene expression signatures uniquely specific to individual CMs, an Orthogonal Partial Least Squares (OPLS) regression framework was deployed iteratively for each defined CM^44^. For a target CM, a binary indicator vector was established, assigning a value of 1 to all constituent cell types within that CM and 0 to all remaining cell types across the dataset. The OPLS model was trained on the pseudobulk expression matrix to isolate and filter out systematic transcriptomic variation orthogonal to the binary target class.

Model performance and generalization were rigorously evaluated using a 5-fold stratified cross-validation scheme coupled with a single-component Partial Least Squares (PLS) regression model. Following cross-validation, a final model was fitted on the fully OPLS-processed dataset, and the resulting feature weights (gene loadings) from the first predictive component were extracted to quantify each gene’s relative contribution to CM segregation.

To decode the biological pathways governed by these transcriptional signatures, a functional annotation approach was executed using the decoupler framework^93^ against the Molecular Signatures Database (MSigDB). First, discrete CM signatures were captured by isolating the top 40 genes with the highest positive predictive OPLS loadings. These gene sets were then evaluated via over-representation analysis (ORA) across several curated gene set collections, including Gene Ontology (GO) Biological Processes, Molecular Functions, and Cellular Components, alongside KEGG, Reactome, Hallmark, and ImmuneSigDB. To ensure robustness, pathways with fewer than 5 overlapping genes or an over-representation significance of p>0.05 were filtered out. The resulting statistical estimates and significance values directly informed the ultimate biological nomenclature assigned to each individual CM.

#### Calculation of Sample-Level CM Activity Scores

To quantify the activity of these cellular programs across donors, sample-specific CM activity scores were computed. First, the log-transformed per-sample cell type proportions were standardized globally across the entire dataset by computing the Z-score for each cell type. For a given sample, the activity score for each CM was defined as the mean of the standardized z-scores of its constituent cell types. To mitigate the influence of extreme outliers, the final sample-level activity scores were capped at an upper threshold of z = 3.

### Cell-subset compositional changes in healthy individuals

#### Data processing and sample filtering

Per-donor cell-type proportions were computed as the fraction of cells belonging to each annotated granular cell subset relative to all cells within the broad lineage compartment it belongs to (e.g., CD4 T, CD8 T, B cell, Myeloid) for each donor. Donors lacking age or sex metadata were excluded. Donors younger than 1 year or older than 80 years were removed owing to sparse representation across studies. Cell types were retained for downstream analyses only when data were available from at least 15 donors across a minimum of 2 independent studies.

#### Study-effect correction and computation of corrected residuals

To reduce study-specific batch effects while preserving biological variation attributable to age and sex, log10-transformed proportions were regressed on study identity independently for each cell type using an ordinary least-squares (OLS) linear model:

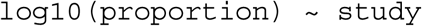

Residuals from this model were retained as study-corrected values (resid_study). A grand-mean shifted version (resid_study_scaled = resid_study + mean(log10(proportion))) was used for visualization to restore values to the original log10(proportion) scale. For heatmap visualization, study-corrected residuals were averaged across donors within each age bin and subsequently z-scored per cell type to emphasize relative temporal trajectories independent of absolute abundance differences.

#### Linear mixed-effects modelling of age and sex effects

To assess the effects of age, sex, and their interaction on cell type abundance, linear mixed-effects models (LMMs) were fitted independently for each cell type using the lmerTest R package:

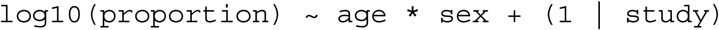

where age was treated as a mean-centered continuous variable and study was included as a random intercept to account for cohort-level variation. The significance of each fixed-effect term (`age, sex, age × sex interaction`) was evaluated using Satterthwaite-approximated *t*-tests, as implemented in lmerTest^94^ (coef(summary(fit))). P-values for each term were then corrected across cell types using the Benjamini–Hochberg (BH) procedure. For heatmap visualization, signed *t*-statistics were used to display effect direction and magnitude; non-significant terms (BH-adjusted *p* ≥ 0.10) were attenuated by a factor of 0.25 to visually distinguish confirmed from unconfirmed trends.

#### Non-linear age trajectories and identification of sex-divergent cell types

To identify non-linear and sex-divergent abundance trajectories, a two-step quadratic modelling approach was applied to study-corrected residuals. A quadratic model was selected as higher order models risk overfitting noise across the seven age bins. This quadratic fit provided the minimal parameterization needed to detect non monotonic trends such as midlife surges or late life divergence.

Age was discretized into seven bins (1–20 through 70–80 years) and parameterized as a continuous integer index (1–7). In the first step, a full quadratic-by-sex interaction model was compared against a sex-main-effect null model (retaining sex as a covariate) using OLS regression:

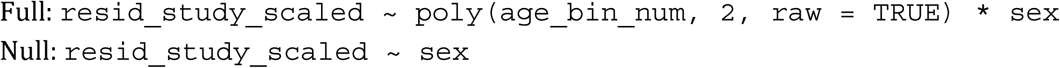

Statistical significance was assessed using a partial F-test (via ANOVA model comparison) between the two models. P-values were BH-corrected across all tested cell types, and only cell types with adjusted p < 0.05 were retained for reporting and visualization.

In the second step, sex-stratified quadratic models were fitted independently within each cell type:

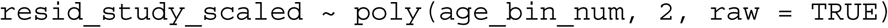

Fitted values and 95% confidence intervals were estimated across age bins for visualization. Sex divergence at a given age bin was defined by non-overlapping 95% confidence intervals between the female and male fitted trajectories at that bin’s exact position, with the direction of divergence (female-higher or male-higher) recorded accordingly.

### Cell-subset compositional changes in healthy and COVID-19 individuals

Only a single time point (the initial sample collection time point, if multiple samples were available) was analysed per donor. Cell subsets from three major cell lineages (B and plasma cells; myeloid and dendritic cells; and ILC, NK, and T cells) were analysed. Depending on which major cell lineage a cell subset belonged to, only donors with at least 50 B or plasma cells, or 50 myeloid or dendritic cells, or 500 ILC/NK/T cells, respectively, were included for analysis. Following this donor-level filtering, only cell subsets with, on average, at least 10 cells per donor were included in the downstream analysis. Donors with any missing sex or age bin metadata were then excluded from the analysis.

The log_10_(proportion) of each cell subset was computed per donor. We checked for robustness of our findings across two types of denominators: either all PBMCs per donor, or all cells of a major cell lineage that a cell subset of interest corresponded to (i.e., B and plasma cells; or myeloid and dendritic cells; or ILC, NK, and T cells) per donor. Donors from 20 to 79 years old were analysed for association with COVID-19 disease, and, separately, for correlation with disease severity, and for interactions between age or sex and moderate/severe/critical COVID-19 disease status. Given the available donor metadata, donors were categorised across four age bins. These age bins were designated as ordered factors: 20-35 < 36-49 < 50-64 < 65-79 years old, with up to cubic relationships considered for age bins. Only cell subsets that were present in at least 10 donors, each across at least 3 studies, were retained for downstream analysis, and only studies with at least 10 donors were retained for each cell subset.

For association with COVID-19 disease alone, healthy / control donors and donors with mild, moderate/severe, or critical COVID-19 disease status were analysed. A linear model was implemented using R:

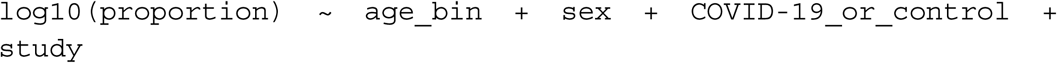

For correlation with COVID-19 disease severity, donors with mild, moderate/severe, or critical COVID-19 disease status were analysed. Disease severity was designated as ordered factors: mild < moderate/severe < critical, with up to quadratic relationships considered for disease severity. A linear model was implemented using R:

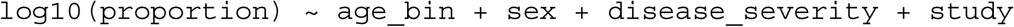

For interaction terms involving age and sex, healthy / control donors as well as donors with moderate/severe or critical COVID-19 disease status were analysed. A linear model was implemented using R:

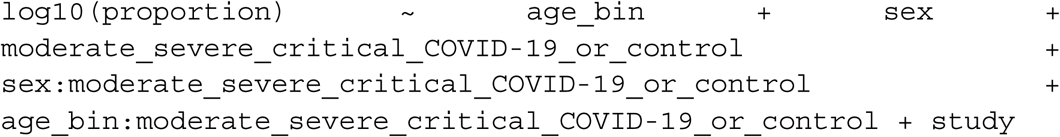

False discovery rate (FDR) values were computed for each of the above models separately, by applying BH correction across the p-values for all terms in the model output across all cell subsets analysed.

For plotting individual examples, the effect of study, as well as any age bin or sex demographic not depicted in the plot, was regressed out from the log_10_(proportion) values, leaving the residual values. For heatmap visualization, only cell subsets with at least one coefficient of interest showing FDR < 0.05 were retained. Each coefficient of interest was Z-scored separately across all cell subsets, and cell subsets were hierarchically clustered based on their Z-scored coefficients.

### Differential gene expression

#### Differential gene expression analysis in healthy individuals

To assess differential gene expression changes with sex and age in control donors, we computed pseudobulk profiles at the Level 4 cell subset level. Samples with missing age, sex, disease status were excluded prior to aggregation, and pseudobulk profiles were generated for each cell subset by summing counts within each donor sample. Only profiles derived from at least ten cells, from donors aged 1–79 years, and with at least 500 detected genes were retained. Studies with >80% representation of a single sex were excluded to minimise confounding between sex and study. Differential expression was performed on these pseudobulk profiles using limma^91^. Library size normalization was performed using the trimmed mean of M-values (TMM) method implemented in edgeR^95,96^, followed by mean–variance modelling with voomWithQualityWeights (limma). Differential expression was tested using linear mixed-effects models implemented in dream (variancePartition)^97^, including sex and age (modelled as a second-order polynomial of age_midpoint) as fixed effects and study as a random effect: or

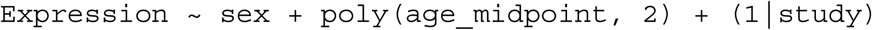

Raw p-values obtained from the linear mixed-effects models were moderated using the empirical Bayes framework implemented in limma (eBayes). Multiple-testing correction was performed using the BH procedure. Genes with an FDR < 0.05 and |logFC| ≥ 0.25 were considered differentially expressed.

#### Gene programs (GPs)

To identify gene programs (GPs) shared across cell subsets within coabundance-associated cell modules (CMs) that vary with sex and age, we investigated recurrent differentially expressed genes (DEGs) across the 131 cell subsets with significant sex-associated DEGs from the additive model (Expression ∼ sex + poly(age_midpoint, 2) + (1|study)). TCR, BCR, long non-coding/unannotated, open-reading-frame genes, and c1orf56^HI^ cell subsets were removed prior to gene program (GP) identification. To identify coordinated GPs associated with sex, we retained DEGs (FDR < 0.05, |logFC| ≥ 0.3) that were recurrent in at least 4 cell subsets, applying a per-cell-subset threshold of at least 10 significant DEGs, resulting in 246 DEGs across 108 cell subsets. These DEGs were separated by direction (logFC > 0 (male) vs logFC < 0 (female) and jointly assigned to pathway themes identified by over-representation analysis (ORA; Enrichr, against MSigDB_Hallmark_2020, KEGG_2021_Human, GO_Biological_Process_2023, and Reactome_2022). DEGs were then manually grouped into GPs based on shared expression patterns across cell subsets and CMs, and were retained only when at least three genes converged on a biologically coherent program. DEGs were subsequently curated such that each DEG was assigned to a single, non-overlapping GP with shared biological function and similar sex direction across CMs. A full heatmap of all the cell subsets prior to filtering that belong to CMs is shown in Supplementary Fig. 8. This strategy yielded a final curated set of 141 DEGs across 50 cell subsets, with FDR < 0.05 and |logFC| ≥ 0.25.

#### GP scoring in COVID-19

To assess how healthy/control-associated GPs shift with COVID-19 across sexes, we further scored GP activity in disease condition. After retaining GPs with ≥7 genes, we computed sample-level module scores restricted to Day-0/cross-sectional timepoints (D0, T0, or no timepoint annotation - the latter corresponding to single sample per donor), avoiding non-independent repeated measures. Pseudobulk gene expression was generated by summing raw counts per sample per cell subset, converted to log2(CPM+1), and Z-scored within each study to remove study-level batch effects. GP scores were calculated as the mean Z-scored expression across GP genes per sample. For each cell subset within a CM, module scores were compared between healthy males versus COVID-19 males, healthy females versus COVID-19 females, and between male and female COVID-19 donors, using ordinary least squares (score ∼ group + age_midpoint), with age included as a covariate to account for age-related variation independent of disease status. P-values were adjusted using the BH procedure across all cell subsets and contrasts within each CM.

For COVID-19 only donors, GP scores were further tested for association with COVID-19 severity (mild, moderate/severe, critical), after restricting cell subsets with at least 3 samples per severity level. Severity dependence was tested separately per sex using OLS (`score ∼ sev_lin + sev_quad + age_midpoint`), where `sev_lin (mild = −1, moderate/severe = 0, critical = +1)` codes a monotonic linear trend and `sev_quad (mild = +1, moderate/severe = −2, critical = +1)` codes a non-monotonic (U-shaped) deviation from linearity. As above, p-values for the linear and quadratic terms in each sex were adjusted using the BH procedure across all cell subsets and contrasts within each CM.

#### Pearson correlations

Pearson correlations were computed using female sample-level pseudobulk data. For each sample–cell subset combination, *PTGS2* expression was quantified as log2p(CPM + 1) from pseudobulk counts, retaining only pseudobulk profiles derived from a minimum of 10 cells. The innate inflammatory GP score was defined as the mean log2p(CPM + 1) expression across GP member genes, excluding *PTGS2* itself (*ABCA1, APOBEC3A, BCL2A1, CCL3, CXCL2, CXCL8, FCGR2B, HES1, HES4, ITGAX, LMNA, OSM, RASGEF1B, S100B, SELL, SIGLEC1, SNAI1, ZNF331*). Donor–cell subset observations with zero *PTGS2* or a mean GP score of zero were excluded. Correlations were computed across all myeloid cell subsets from CM1, CM4, and CM5, and p-values were adjusted across all tested cell subsets using the BH procedure.

### External dataset validation of transcriptional changes associated with sex

#### Prostaglandin gene expression in the Chinese Multiome Atlas

To cross-check the prostaglandin sex-associated axis in a separate cohort, we investigated the Chinese Immune Multi-Omics Atlas (CIMA)^49^. Normalized single-cell RNA expression values were extracted for *PTGDS* and *PTGS2*. For each sample, expression was averaged within each annotated cell type at Level 1 and Level 4, generating sample-by-cell-subset expression matrices. Sex differences in *PTGDS* and *PTGS2* expression were then tested within each cell type/subset using two-sided Mann-Whitney U tests comparing female and male samples, followed by BH FDR correction across the tested cell subsets within each analysis. Level 1 plots were generated across all broad cell types for each gene. At Level 4, *PTGDS* was tested in 13 cytotoxic lymphocyte subsets spanning NK, CD8 cytotoxic T, and gamma-delta T populations, while *PTGS2* was tested in 13 myeloid subsets spanning monocytes, dendritic cells, and pDCs.

#### Plasma protein inspection in the UKBB samples

For protein-level validation, we used Supplementary Table 5 (ST5) from the UK Biobank Pharma Proteomics Project Olink dataset^54^, which reports multivariable linear regression associations between protein abundance and age, sex, and BMI. We focused on the sex association statistics, using the reported male beta coefficient as the effect-size estimate and the corresponding two-sided unadjusted *P* value, represented in ST5 as −log10(*P*). Protein IDs were mapped to gene symbols using the Olink protein identifier prefix, and these were intersected with non-sex-chromosome genes from the recurrent sex-associated gene programs (GPs) identified in our HARP transcriptomic analysis. IFN-related proteins were inspected manually and included IFNAR1, IFNG, IFNGR1, CXCL9, CXCL10, and CXCL11. Volcano plots were generated with the male-female difference in protein abundance on the x-axis and −log10(*P*) for the sex association on the y-axis. Proteins were highlighted and labelled if they passed both the ST5 significance threshold (*P* < 1.7 × 10⁻⁵) and an effect-size threshold of |beta| > 0.1; all other matched proteins were shown in grey.

### scTiger: transfer of HARP annotations to external datasets

#### Overview of scTiger

To enable transfer of our HARP reference annotations to external datasets, we developed scTiger, a hierarchical reference-mapping framework that combines lineage classification with lineage-specific metric learning. Our framework was designed to enable accurate and scalable annotation of unseen PBMC datasets with the capacity to resolve rare immune populations. Rather than learning a single representation for all PBMCs, scTiger first assigns cells to broad immune lineages using an XGBoost-based classifier, then subsequently projects cells on to independently learned lineage-specific latent spaces to perform annotation. This hierarchical, lineage-based design restricts comparisons to transcriptionally related populations and reduces the associated computational complexity.

#### Training the XGBoost-based models for lineage assignment

HARP cells were first partitioned into broad lineages: CD4 T cells; CD8 T cells and NK cells; B and plasma cells; monocytes and DCs; and hematopoietic stem and progenitor cells (HSPC) on the basis of their HARP annotations. These assignments were used as ground-truth, supervised labels for training the XGBoost models. Separate binary classifier models were trained for each lineage decision (e.g. CD4 T cell vs non-CD4 T cell) rather than assigning cells using a single multiclass classifier. This hierarchical design was chosen to provide explicit control over the transcriptional features used at each decision point. Each binary classifier was trained using a curated marker gene set selected for their ability to distinguish the populations of interest while limiting the contribution of unrelated or potentially confounding transcriptional signals. In testing this produced more robust results than a single multiclass classifier which learns features jointly across all populations.

Each model was constructed using the “XGBoost” package with a binary objective function. The parameters were set as a maximum depth of 8 (max_depth), learning rate of 0.05, and the number of iteration rounds 100 (num_boost_rounds). Histogram-based tree construction (tree_method = “hist”) was used with a maximum of 256 histogram bins (max_bin = 256), and area under the receiver operating characteristic curve (auc) was specified as the evaluation metric. Predicted probabilities were converted to binary class assignments using a probability threshold of 0.5. Cells remaining unassigned following hierarchical XGBoost classification were assigned using k-nearest-neighbor classification in the Harmony-corrected PCA space (X_pca_harmony). Each unassigned cell was assigned to the most frequent lineage among its 50 nearest classified neighbors

#### Training lineage-specific metric-learning models

Following lineage partitioning, independent fully connected neural-network encoders, optimized using a triplet-loss function as described previously^61^, were trained for the CD4 T-cell, CD8 T/NK-cell, B-cell/plasma-cell, myeloid/dendritic-cell, and HSPC compartments. 31,908 genes were common to HARP, the Gong *et al.* dataset^6^, and the Terekhova *et al.* dataset^48^; all models were trained on these intersecting genes. Raw counts were normalized to a total of 10,000 counts per cell and log1p-transformed, after which expression values were standardized gene-wise prior to model training. The resulting 31,908-dimensional expression profiles were provided as input to each lineage-specific encoder. Each model projected these expression profiles through two fully connected hidden layers into a 128-dimensional latent embedding. The standard encoder architecture consisted of hidden layers containing 512 and 256 units, respectively, with ReLU activation, followed by a 128-dimensional embedding layer. An alternative architecture used for selected lineage-specific models incorporated batch normalization, ReLU activation and dropout following each of two hidden layers, with the second hidden layer containing half the number of units of the first. The architecture and dimensionality associated with each trained model were retained in the corresponding model checkpoint. Embeddings generated by the final encoder layer were L2-normalized, constraining cells to a common normalized latent space for metric learning and subsequent similarity-based reference mapping. A linear classification head operating on the normalized embedding was additionally used to provide direct supervision of fine-grained cell identity during model training. scTiger employs an encoder-only architecture optimized specifically for cell-type annotation. No decoder was used and the models were not trained to reconstruct the input gene-expression profile. Instead, the encoder was optimized to organize cells in latent space according to their curated fine-grained HARP annotations. This design reduces model complexity and focuses the learned representation on transcriptional features that discriminate cell identities within each immune lineage.

#### Model optimization

Model parameters were learned using a joint optimization objective comprising supervised triplet loss, cross-entropy classification loss, and center loss. Triplet loss was used to structure the latent embedding by encouraging cells with the same fine-grained HARP annotation to occupy nearby positions while increasing separation between cells belonging to different annotations. Training examples were therefore defined using an anchor cell, a positive cell sharing its fine-grained annotation, and a negative cell belonging to a different annotation. Cross-entropy loss was calculated from the classification head and provided direct supervision of fine-grained cell identity. Center loss was additionally used to reduce intra-class variability by encouraging embeddings from cells belonging to the same annotation to remain close to their corresponding class center. Together, these objectives encouraged the formation of compact cell-type-specific regions in latent space while maintaining separation between transcriptionally related populations.

Because the abundance of individual immune populations within HARP varied substantially, weighted oversampling of rare cell types was applied during model training. This increased the representation of low-frequency annotations during optimization and reduced the tendency for abundant populations to dominate model updates. Each lineage-specific model was trained independently using only cells and fine-grained annotations belonging to its corresponding immune lineage.

#### Construction of the scTiger reference

Following training, HARP cells were projected through their corresponding lineage-specific encoders to generate the final reference latent spaces. Each reference cell was represented by a 128-dimensional L2-normalized embedding together with its manually curated fine-grained HARP annotation. The resulting reference embeddings were precomputed and stored together with their associated annotations, ordered reference gene sets, model metadata, and trained encoder parameters. This allowed the same trained HARP reference to be reused for annotation of additional datasets without retraining the metric-learning models or recomputing the reference embeddings.

Reference embeddings were generated separately for each major immune lineage. Consequently, after broad lineage assignment, query cells were compared only with HARP cells belonging to the corresponding lineage rather than with the complete ∼9-million-cell reference atlas. This lineage-restricted nearest-neighbor search both reduced the number of reference comparisons required for annotation and restricted candidate annotations to biologically related immune populations.

#### Reference mapping query datasets to HARP annotation

Query gene-expression matrices were first aligned to the 31,908-gene feature space defined during model development, with genes reordered to match the input feature order of the corresponding trained model. Raw query counts were normalized to a total of 10,000 counts per cell and log1p-transformed. These normalized expression values were used as input to the hierarchical XGBoost classifiers for lineage assignment. Following lineage assignment, expression profiles were further standardized gene-wise using the mean and standard deviation calculated across the query dataset and reordered to match the input gene order of the corresponding lineage-specific encoder. The resulting standardized expression profiles were then used for projection into the lineage-specific scTiger latent spaces.

Query embeddings were compared with the precomputed HARP reference embeddings from the corresponding lineage using cosine similarity. Because both query and reference embeddings were L2-normalized, cosine similarity was calculated directly from the dot product between embedding vectors. For each query cell, the 25 reference cells with the highest similarity were identified. Fine-grained cell identity was then assigned by unweighted majority voting among the HARP annotations of these 25 nearest reference neighbors. Specifically, the fraction of neighbors belonging to each candidate annotation was calculated, and the annotation receiving the greatest fraction of neighbor votes was assigned to the query cell. The fractions supporting the highest- and second-highest-ranked annotations were additionally retained to quantify the local support for each annotation.

This inference procedure requires only preprocessing and alignment of the query expression matrix, hierarchical lineage assignment, forward projection through the appropriate trained encoder, and lineage-restricted nearest-neighbor search. Retraining or fine-tuning of the HARP reference models is therefore not required when annotating a new dataset. Using this framework, annotations from the ∼9-million-cell HARP reference atlas were transferred to independent datasets, including the ∼16-million-cell Gong *et al.* Allen Institute Human Immune Health Atlas^6^ and the ∼2-million-cell Terekhova *et al.* dataset^48^, enabling harmonized high-resolution annotation across large-scale PBMC resources.

#### Software implementation and code availability

scTiger was implemented in Python, using XGBoost for hierarchical lineage classification and PyTorch for implementation and inference of the lineage-specific neural-network encoders. The software provides workflows for query preprocessing and reference-gene alignment, hierarchical lineage assignment, generation of lineage-specific latent embeddings, nearest-neighbor reference mapping and assignment of fine-grained HARP annotations. Trained model checkpoints contain the parameters required to reconstruct the corresponding lineage-specific encoder together with reference embeddings, reference annotations, ordered gene sets, and model metadata, allowing the trained HARP reference to be applied to new datasets without retraining.

### Benchmarking of scTiger celltype annotations

To evaluate the accuracy of scTiger’s cell-type annotations against established single-cell annotation frameworks, we compared scTiger’s performance with scANVI^14^ and CellTypist^62^. Benchmarking was performed using the HARP reference dataset and ground-truth cell-type labels were derived from the HARP Level 3 and Level 4 annotations. Performance was evaluated by assessing how accurately each method predicted ground-truth annotations across held-out evaluation batches. scANVI was trained using the standard scvi-tools workflow (docs.scvi-tools.org). Due to memory constraints when scaling scANVI to the full HARP reference, it was trained on a randomly sampled 50% subset of the HARP training data. CellTypist was trained on the full HARP reference following its recommended custom model-training workflow (github.com/Teichlab/celltypist). Predictions for both comparison tools were evaluated on the exact same held-out batches as scTiger.

To quantify classification accuracy, predicted annotations from scTiger, scANVI, and CellTypist were compared against the ground-truth HARP annotations on the held-out test set. Model performance was evaluated across all matching cell types using F1 and balanced accuracy scores.

### Assessment of external CellTypist model for HARP dataset annotation

To assess the correspondence between HARP annotations and those from existing PBMC annotation tools, we applied the pretrained CellTypist model provided by Gong et al.^6^ to the HARP reference dataset. Cell-type labels were assigned using the external model without retraining or fine-tuning on HARP annotations. A confusion matrix was generated comparing HARP Level 4 annotations with predictions from the pretrained Gong et al. CellTypist model. Patterns of correspondence between fine-grained HARP cell subsets and predicted CellTypist categories were qualitatively assessed to evaluate differences in annotation granularity, including the extent to which distinct HARP subsets corresponded to shared CellTypist labels.

## Extended figures

**Extended Figure 1. Summary of hierarchical organisation of annotated cell subsets and cell subset abundance (related to Main** Figure 3). Cell subsets are grouped by Level 1, Level 2, and Level 3 annotations, with Level 4 cell subsets shown as row labels. Colors indicate lineage/sublineage groupings within the annotation hierarchy. Bold Level 4 labels denote cell subsets we identified as novel/rare populations. Barplots show (from left to right) the average frequency per donor, the frequency across all cells, and the total number of cells per Level 4 cell subset represented as log10-transformed cell counts. Cell subsets are ordered within each Level 1 annotation according to their numbered Level 4 annotation in Figure 3A. Abbreviations: Int. mono = intermediate monocytes, plasmab. = plasmablasts; atyp. = atypical B cells; eff. = effector; trans. = transitional, HSC = Hematopoietic Stem Cell, MPP = Multipotent Progenitor, CLP = Common Lymphoid Progenitor, CMP = Common Myeloid Progenitor, BEMP = Basophil/Eosinophcil/Mast Cell Progenitor, ERYP = Erythroid Progenitor, MEP = Megakaryocyte–Erythroid Progenitor.

## Supplementary figures

**Supplementary Figure 1. Assessment of data integration strategies (related to Main** Figure 2**). (A)** Linear variance contributions of technical covariates towards merged count matrix from study-provided count matrices published on CZ CELLxGENE^82^ compared to merged count matrix from remapped studies. **(B)** Overview of fraction of cells removed per study and per cell lineage using semi-automated QC approach. **(C)** Schema of global vs lineage-specific integration and evaluation workflow as implemented with scAtlasTb. **(D)** Benchmark of lineage-specific and global integrations across different integration methods evaluated per lineage. **(E,F)** UMAP of additional Moran’s I examples for **(E)** cDC2, T cell doublets, and **(F)** γδ T cells highlighting differences in biologically relevant gene set score correlation with kNN graph between Harmony (top) and scVI (bottom) integrations.

**Supplementary Figure 2. Batch analysis for determining optimal batch variable (related to Main** Figure 2**). (A)** Principal component regression analysis for technical and biological covariates. Significance is calculated based on a permuted background distribution per covariate (Methods). **(B)** Confounding analysis of technical and biological covariates using Theil’s U. Lower values indicate less confounding between two covariates; Theil’s U is non-symmetric; given one variable, it represents how much information it explains in another. **(C)** PCA plots of pseudobulk profiles (per smallest entity of a batch; Methods) for full dataset, colored by library ID, sampled site condition, assay, and study.

**Supplementary Figure 3. UMAPs with gene set scores and marker genes (related to Main** Figure 2**). (A)** Gene set score and gene expression for CD16 C1Q^Hi^ monocytes in myeloid lineage. **(B)** Gene set score and gene expression for cDC2 cells in myeloid lineage. **(C)** Gene set score and gene expression for γδ T cells in CD8 T, NK, and unconventional T cell (UTC) population.

**Supplementary Figure 4 (Related to Main Figure 3). Contributions of different lineages to co-abundance Cellular Modules (CMs).** (**A**) Density plot UMAPs of all 13 CMs (extension of Main Fig 3E), illustrating the varying degrees of multi-lineage cell subset composition across CMs. Density scales are normalized per cell type to emphasize localized peaks across low- and high-abundance populations.

**Supplementary Figure 5 (Related to Main Figure 3). Annotation evidence for monocyte, dendritic cell, B, and plasma cell subsets.** (**A**) Dotplot of marker gene expression, annotated by cell numbers for each of the monocyte and dendritic cell subsets (horizontal bars). The background shading of the cell subset labels on the y-axis is color-coded to match the colors of their respective sublineage titles in (B). (**B**) UMAP embeddings of monocyte and dendritic cell populations from Mono/cDC and cDC/pDC sublineages. *Note that cDCs were included in the Mono/cDC embedding for additional context to annotate Monocyte subpopulations; however, cDC cells were ultimately annotated using the cDC/pDC embedding. (**C**) UMAP embeddings of B/Plasma cell populations from B Naive, B Memory, and Plasma sublineages. (**D**) Dotplot of marker gene expression, annotated by cell numbers for each of the B/Plasma cell subsets (horizontal bars). The background shading of the cell-type labels on the y-axis is color-coded to match their respective sublineage title color in (**C**). Numerical identifiers for all cell subsets correspond directly to those in the full lineage UMAPs in Figure 3A.

**Supplementary Figure 6 (Related to Main Figure 3). Annotation evidence for CD4 and CD8 T cell subsets.** (**A**) Dotplot of marker gene expression, annotated by cell numbers for each of the CD4 T cell subsets (horizontal bars). The background shading of the cell-type labels on the y-axis is color-coded to match their respective sublineage title color in (**B**). (**B**) UMAP embeddings of CD4 T cell populations from T Naive/Central Memory and CD4 T Effector sublineages. (**C**) UMAP embeddings of CD8 T/innate populations from CD8 T Naive/Central Memory and CD8 T Effector/innate lymphoid cells (“Innate”) sublineages. (**D**) UMAP embedding of Proliferating T and NK cells which span CD4 T, CD8 T, and NK lineages.

**Supplementary Figure 7 (Related to Main Figure 3). Annotation evidence for CD8 T, gdT, innate lymphoid cell, and hematopoietic stem and progenitor cell (HSPC) subsets.** (**A**) Dotplot of marker gene expression, annotated by cell numbers for each of the CD8 T, gdT, and innate lymphoid cell (including NK cell) subsets (horizontal bars). The background shading of the cell-type labels on the y-axis is color-coded to match their respective sublineage title color in Supplementary Fig. 6C. Numerical identifiers for all cell subsets correspond directly to those in the full lineage UMAP in Figure 3A. (**B**) UMAP embeddings of HSPC cell subsets (**C**) Dotplots of marker gene expression for the HSPC cell subtypes. HSC = Hematopoietic Stem Cell, MPP = Multipotent Progenitor, CLP = Common Lymphoid Progenitor, CMP = Common Myeloid Progenitor, BEMP = Basophil/Eosinophil/Mast Cell Progenitor, ERYP = Erythroid Progenitor, MEP = Megakaryocyte–Erythroid Progenitor.

**Supplementary Figure 8. Age and sex-associated variation in immune cell-subset abundance and gene program activity (related to Main** Figure 4). (**A**) Summary heatmap showing age-associated cell subset abundance trajectories and linear mixed-effects model (LMM) coefficients across cellular modules (CMs). Upper panel shows study-corrected cell-type proportions averaged across age bins and z-scored per cell subset to highlight relative temporal dynamics. The lower panel shows scaled LMM coefficients for age, sex, and age-by-sex interaction terms. Cell subsets are annotated by Level 1 annotation and grouped by CMs. Asterisks denote BH-adjusted p < 0.05. (**B**) Barplots of 122 cell subsets (Level 4 annotation) used for DEG analysis. Upper panel shows the sex composition of pseudobulk samples (red = female, blue = male; n = total samples per cell subset). Lower panel shows the direction of significant sex-associated DEGs (BH-adjusted p < 0.05, |logFC| ≥ 0.25; red = female-higher, blue = male-higher; n = total significant DEGs per cell subset). (**C**) Heatmap of log fold change (LFC) across all 98 cell subsets belonging to CMs, prior to filtering for cell subsets with ≥10 DEGs and a consistent sex direction, colored as in Fig. 4E (red = female-high, blue = male-high expression). Filled circles denote BH-adjusted p < 0.05 and |LFC| ≥ 0.25; grey cells indicate cell subsets not tested due to power limitations.

**Supplementary Figure 9. Sex-associated variation in cell type-specific gene expression profiles (related to Main** Figure 4**). (A**) Pathway enrichment analysis of recurrent sex-DEGs, for females (left) and males (right). **Left:** Over-representation analysis (Enrichr, MSigDB Hallmark 2020) of the 177 female-higher genes from recurrent sex-DEG set (adj. p < 0.05, |logFC| > 0.3, recurrent in ≥4 cell subsets). Bars show the top 15 significantly enriched terms (adj. p < 0.05), ranked by −log₁₀(adjusted p-value). **Right:** Over-representation analysis (Enrichr, GO Biological Process 2023) of the 69 male-higher genes from the same recurrent sex-DEG set. Bars show the top 14 significantly enriched terms (adj. p < 0.05), ranked by −log₁₀(adjusted p-value). **(B)** Beeswarm plot showing average log fold change (LFC) of selected prostaglandin pathway genes (**PTGDS, PTGDR, PTGDR2, PTGS2, PTGES, PTGER2, PTGER4**) across cell subsets. Each circle represents one cell subset; circles with bolded circumferences indicate cell subsets with statistical significance. Circle colour reflects the direction and magnitude of the LFC (red = female-high, blue = male-high). **(C)** Barplots of mean log2p(CPM + 1) **PTGDS** expression per sample in the Chinese Multiome Atlas^49^, shown across Level 1 cell annotations (left) and Level 4 NK/CD8 CTL/gamma-delta T cell subsets (right). Each dot represents one sample, coloured by sex (red = female, blue = male). Coloured bars indicate group means. BH-corrected p-values from Mann-Whitney U tests are shown above brackets. **(D)** Barplots of mean **PTGS2** expression per sample in the Chinese Multiome Atlas, shown across Level 1 annotations (left) and Level 4 monocyte/DC/pDC cell subsets (right). Each dot represents one sample, coloured by sex (red = female, blue = male). Coloured bars indicate group means. BH-corrected p-values from Mann-Whitney U tests are shown above brackets. (**E**) Scatter plots showing the correlation between **PTGS2** expression and innate inflammatory gene program (excluding **PTGS2**) activity across myeloid cell subsets in female samples. Each panel corresponds to one Level 4 cell subset (minimum 10 female samples), labelled by CM (CM1, CM4, or CM5) and cell subset. Each point represents one female sample. Black lines show linear fits. Panel labels report Pearson **r**, BH-adjusted **p**-value across the 25 tested cell subsets, and sample size.

**Supplementary Figure 10. Disease, age, and sex-associated variation in immune cell-subset abundance and gene program activity during COVID-19 (related to Main** Figure 5**). (A)** Lollipop plot depicting per-cell-subset coefficients of the COVID-19 disease versus controls covariate (log_10_(proportion) ∼ age_bin + sex + COVID-19_or_control + study). (**B**) Heatmap of Z-scores of coefficients of the male sex covariate; the interaction term between male sex and moderate, severe, or critical COVID-19 status; the age bin covariate; and the interaction term between age bin and moderate, severe, or critical COVID-19 status (log_10_(proportion) ∼ age_bin + sex + moderate_severe_critical_COVID-19_or_control + sex:moderate_severe_critical_COVID-19_or_control + age_bin:moderate_severe_critical_COVID-19_or_control + study). Z-scores were computed individually for each covariate across all cell subsets in the heatmap, and hierarchical clustering was performed to order the cell subsets. Asterisks indicate FDR < 0.05 for the corresponding coefficient-cell subset combination. Cell subsets are annotated by their CM membership. Proportions for each cell subset per donor were computed out of all PBMCs per donor. **(C)** Lollipop plots for three myeloid GPs not shown in Fig. 5C - Hypoxia/Metabolic Stress, Stress/Apoptosis, and Cytotoxic - across 14 myeloid cell subsets. Score computation, OLS contrasts, BH correction of p-values, and display conventions match Fig. 5H. **(D)** Severity-dependent IFN-related GP scores across all 14 myeloid cell subsets, complementary to Fig. 5I. Statistical framework, boxplot display, and annotations match Fig. 5I. **(E)** As in (D), for the Immunomodulation GP. **(F)** As in (D), for the Inflammatory GP.

**Supplementary Figure 11. Disease and sex-associated variation in gene program activity during COVID-19 (related to Main** Figure 5). (**A**) Lollipop plots of mean GP scores for eight GPs - IFN-related, Activation, Immunomodulation, Hypoxia/Metabolic Stress, Cytotoxic, Adhesion/Migration, Stress/Apoptosis and NF-kB signaling - across 35 lymphoid cell subsets, grouped by CM (CM9: B/Humoral memory; CM13: Innate-like/Cytotoxic; CM3: Activated Innate; CM2: Cytotoxic; CM7: Helper Humoral; CM6: Helper Effector; CM10: Naive/RTE). GP scores are computed as in Fig. 5H (within-study z-scored log₂(CPM +1) per gene, averaged across GP genes, OLS age residualisation), with the same three OLS contrasts per cell subset (female COVID vs. healthy; male COVID vs. healthy; female vs. male within COVID) and global BH correction of p-values per GP across lymphoid CMs × 3 contrasts. Visualization conventions and statistical significance thresholds as in Fig. 5H.

**Supplementary Fig. 12. Validation of rare and high-resolution immune cell subsets across independent datasets (related to Main** Figure 6). Dotplots showing marker gene expression for selected rare and fine-grained immune populations identified by scTiger, annotated by number of cells per cell subset (horizontal bars). Cell subsets include activation-associated, interferon-responsive, and lineage-specialized lymphoid, myeloid and NK-cell populations. Dot size indicates the fraction of expressing cells and color denotes mean expression. **(A)** Marker gene expression in the HARP reference atlas used for model training. **(B)** Corresponding expression profiles in the held-out validation dataset comprising unseen batches and studies. **(C)** Expression profiles in the external Gong et al. dataset following reference mapping. **(D)** Expression profiles in the external Terekhova et al. dataset following reference mapping.

**Supplementary Fig. 13. Additional benchmarking and interpretability analyses of scTiger (related to Main** Figure 6**). (A,B)** Additional performance evaluation of scTiger, CellTypist, and scANVI across held-out batches. **(A)** Relationship between balanced accuracy score (BAS) and weighted F1 score (WF1) across Level 2 populations. Each point represents one cell population. **(B)** Mean BAS and WF1 scores for each method; error bars indicate variability across populations. **(C,D)** Computational scalability benchmarking. **(C)** Query runtime as a function of dataset size (50,000– 1,000,000 cells). **(D)** Peak resident memory usage as a function of dataset size. **(E–G)** Interpretability of lineage-specific scTiger models. **(E)** Hierarchical clustering of monocyte and DC population centroids in the learned latent space using cosine distance. **(F,G)** Integrated Gradients feature-attribution scores for the DC_CCR7_LAMP3 and CD14_THBS1 populations, respectively. Canonical marker genes, including **LAMP3**, **CCR7** and **THBS1**, are amongst the highest-contributing features, indicating that lineage-specific models learn biologically meaningful representations and classification programs.

## Acknowledgements

We thank Lisa Dratva, Ioannis Sarropoulos, Ken To, Krzysztof Polanski, Kerstin Meyer, and Nir Hacohen for valuable discussions on the project. We thank John Randell, Lucia Robson, Ellen Todres, and Dave Rogers for their administrative and technical support in this project.

We acknowledge the contribution of Bo Li, Monika S Kowalczyk, Michal Slyper, Gaublomme Jellert, Marcin Tabaka, Orr Ashenberg, Julia Waldman, Danielle Dionne, Knecht Abigail, Ma Hui, Yiming Yang, Orit Rozenblatt-Rosen, Aviv Regev in generating and making publicly available the dataset entitled **Single Cell Immune Cell Atlas of Human Hematopoietic System** (INSDC Project Accessions: ERP122984; BioStudies Accessions: S-SUBS12; https://explore.data.humancellatlas.org/projects/cc95ff89-2e68-4a08-a234-480eca21ce79).

The AIDA project was funded by grants CZF2019-002446 (S.P., W.-Y.P., J.W.S., and J.C.C.) and CZF2021-238829 (5022) (S.P., W.-Y.P., and J.W.S.) from Chan Zuckerberg Foundation and 2020-224570 (S.P., V.C., P.M., and P.P.M.) and 2021-240178 (S.P., W.-Y.P., J.W.S., J.C.C., V.C., P.M., and P.P.M.) from Chan Zuckerberg Initiative (CZI) DAF, an advised fund of Silicon Valley Community Foundation. The AIDA Singapore donor samples were obtained through the Health for Life in Singapore (HELIOS) Study (Lee Kong Chian School of Medicine [LKCMedicine], Nanyang Technological University [NTU]; National Healthcare Group [NHG], Singapore; Imperial College London), which was supported by the Singapore Ministry of Health’s National Medical Research Council (OF-LCG: MOH-000271-00) and intramural funding (NTU; LKCMedicine; NHG). We would like to express our gratitude to AIDA and HELIOS study participants. We thank the HELIOS operation team for recruitment, organization, and data and sample collection, including Yoke Yin Terry Tong, Swat Kim Kerk, Guo Liang Low, and Halimah Binte Ibrahim (HELIOS Biobanking team).

This project was supported by several training awards and fellowships: NovoNordisk NovoSTAR fellowship (A.M.C.; grant number: 47335), Lupus Foundation of America’s Career Development Award (J.N.), Rheumatology Research Foundation’s Career Development Bridge Funding Award (J.N.), Harvard Medical School Eleanor and Miles Shore Faculty Development Award (J.N.), Royal Society Newton International Fellowship (L.K.; grant number:NIF-R1-232597), The Manton Foundation (K.K.), The Leona M. and Harry B. Helmsley Charitable Trust (K.K.), Wellcome Trust [A.Y.; grant number: 226795/Z/22/Z]. Furthermore, this project was supported by the following grants, including CZIF2022-007488 (F.J.T, S.A.T, M.D.L, S.P., A.C.V., G.R.), CZIF2020-216954 (A.C.V.), and CZIF 2023-323359 (A.C.V., G.R.) from the Chan Zuckerberg Initiative Foundation, as well as funding from the Schmidt Futures for the HCA Cell Annotation Platform (A.C.V). Additional support was received from Wellcome Trust (B.G.; grant numbers: 226795/Z/22/Z; 309075/Z/24/Z; 206328/Z/17/Z), the Ministerio de Ciencia e Innovación (MCI) (H.H.; PLEC2021-007654, PID2024-161652OB-I00), the LaCaixa Foundation (H.H.; HR22-0031, HR22-0172 and HR25-00861), the European Union (F.J.T.; ERC, DeepCell - 101054957), and the Dutch Cancer Society and Dutch Ministry of Health, Welfare, and Sport (R.G.H.L.).

## Consortia

1-Below is the list of all MGH COVID Collection & Processing Team participants:

Co-Lead Investigators: Marcia B Goldberg^1-4^, Michael R Filbin^1,5-6^, Moshe Sade-Feldman^1,5,7^, Nir Hacohen ^1,5,7^, Alexandra-Chloe Villani^1,5,7,8^

Collection Team: Kendall Lavin-Parsons, Blair Parry^6^, Brendan Lilley^6^, Carl Lodenstein^6^, Brenna McKaig^6^, Nicole Charland^6^, Hargun Khanna^6^, and Justin Margolin^6^.

Blood Processing Team: Anna Gonye^1,7^, Irena Gushterova^1,7^, Tom Lasalle^1,7^, Nihaarika Sharma^7^, Brian C. Russo^2,3^, Maricarmen Rojas-Lopez^2,3^, Kasidet Manakongtreecheep^1,7,8^, Jessica Tantivit^1,7,8^, Molly Fisher Thomas^1,5,7,8^, and Thomas Eisenhaure^1^.

Single-Cell Data Generation and Analysis: Jessica Tantivit^1,7,8^, Kasidet Manakongtreecheep^1,7,8^, Alice Tirard^1,7,8^, Benjamin Y Arnold^1,7,8^, Miguel Reyes^1^

## Afffiliations

^1^Broad Institute of the Massachusetts Institute of Technology and Harvard University, Cambridge, MA, 2142, USA

^2^Department of Medicine, Division of Infectious Diseases, Massachusetts General Hospital, Boston, MA, 2114, USA

^3^Department of Microbiology, Harvard Medical School, Boston, MA, 2115, USA

^4^Harvard T.H. Chan School of Public Health, Boston, MA, 2115, USA

^5^Harvard Medical School, Boston, MA, 2115, USA

^6^Department of Emergency Medicine, Massachusetts General Hospital, Boston, MA, 2114, USA

7Krantz Family Center for Cancer Research, Massachusetts General Hospital, Boston, MA, 2114, USA

^8^Department of Medicine, Center for Immunology and Inflammatory Diseases, Massachusetts General Hospital, Boston, MA, 2114, USA

## 2-Below is the list of Asian Immune Diversity Atlas (AIDA) Network participants

Atlas assembly authors are arranged by area of contribution and ordered alphabetically by last name.

Single-cell experimental dataset generation leads: Varodom Charoensawan^1,2,3,4,5,6,7^, Chung-Chau Hon^8,9^, Partha P. Majumder^10,11^, Ponpan Matangkasombut^3,12^, Woong-Yang Park^13^, Shyam Prabhakar^14,15,16^, Jay W. Shin^14,17^

Cohort and sample collection leads: Piero Carninci^18,19^, John C. Chambers^15^, Marie Loh^14,15^, Manop Pithukpakorn^6,20^, Bhoom Suktitipat^2,5^, Kazuhiko Yamamoto^21^

Overall study design and protocol optimisation and development: Deepa Rajagopalan^14^, Nirmala Arul Rayan^14^, Shvetha Sankaran^14^

Sample isolation and processing, single-cell experimental data generation: Juthamard Chantaraamporn^1,2,3^, Ankita Chatterjee^10^, Supratim Ghosh^22^, Kyung Yeon Han^13^, Damita Jevapataraku^l3,12^, Sarintip Nguantad^1,2,3^, Sumanta Sarkar^22^, Narita Thungsatianpun^3,12^

Sample isolation and processing: Mai Abe^21^, Seiko Furukawa^21^, Gyo Inoue^21^, Keiko Myouzen^21^, Jin-Mi Oh^13^, Akari Suzuki^21^

Single-cell experimental data generation: Yoshinari Ando^17,18^, Miki Kojima^18^, Tsukasa Kouno^17^, Jinyeong Lim^13^, Arindam Maitra^22^, Le Min Tan^14^, Prasanna Nori Venkatesh^14^

Single-cell experimental data generation and analysis: Murim Choi^23^, Jong-Eun Park^24^

Single-cell data analysis up to cell type annotation: Eliora Violain Buyamin^14^, Kian Hong Kock^14^, Quy Xiao Xuan Lin^14^, Jonathan Moody^8^, Radhika Sonthalia^14^

Genotype QC and imputation: Kazuyoshi Ishigaki^25^, Masahiro Nakano^21,26^, Yukinori Okada^27,28,29,30,31,32^, Yoshihiko Tomofuji^27,28,29^

## Affiliations

^1^Department of Biochemistry, Faculty of Science, Mahidol University, Bangkok 10400, Thailand

^2^Integrative Computational BioScience (ICBS) Center, Mahidol University, Nakhon Pathom 73170, Thailand

^3^Single-cell Omics and Systems Biology of Diseases Research Unit, Faculty of Science, Mahidol University, Bangkok 10400, Thailand

^4^Research Department, Faculty of Medicine Siriraj Hospital, Mahidol University, Bangkok 10700, Thailand

^5^Department of Biochemistry, Faculty of Medicine Siriraj Hospital, Mahidol University, Bangkok 10700, Thailand

^6^Siriraj Genomics, Faculty of Medicine Siriraj Hospital, Mahidol University, Bangkok 10700, Thailand

^7^School of Chemistry, Institute of Science, Suranaree University of Technology, Nakhon Ratchasima 30000, Thailand

^8^Laboratory for Genome Information Analysis, RIKEN Center for Integrative Medical Sciences, 1-7-22 Suehiro-cho, Tsurumi-ku, Yokohama, Kanagawa 230-0045, Japan

^9^Graduate School of Integrated Sciences for Life, Hiroshima University, 1-3-3-2 Kagamiyama, Higashihiroshima, Hiroshima 739-0046, Japan

^10^John C. Martin Centre for Liver Research and Innovations, Sonarpur, Kolkata 700150, India

^11^Indian Statistical Institute, 203 B.T. Road, Kolkata 700108, India

^12^Department of Microbiology, Faculty of Science, Mahidol University, Bangkok 10400, Thailand

^13^Samsung Genome Institute, Samsung Medical Center, Seoul 06351, Republic of Korea

^14^Genome Institute of Singapore (GIS), Agency for Science, Technology and Research (A*STAR), 60 Biopolis Street, Genome, Singapore 138672, Republic of Singapore

^15^Nanyang Technological University, Lee Kong Chian School of Medicine, Clinical Sciences Building, Level 18, 11 Mandalay Road, Singapore 308232, Republic of Singapore

^16^Cancer Science Institute of Singapore, National University of Singapore, 14 Medical Drive, Singapore 117599, Republic of Singapore

^17^Laboratory for Advanced Genomics Circuit, RIKEN Center for Integrative Medical Sciences, 1-7-22 Suehiro-cho, Tsurumi-ku, Yokohama, Kanagawa 230-0045, Japan

^18^Laboratory for Transcriptome Technology, RIKEN Center for Integrative Medical Sciences, 1-7-22 Suehiro-cho, Tsurumi-ku, Yokohama, Kanagawa 230-0045, Japan

^19^Genomics Research Center, Fondazione Human Technopole, Viale Rita Levi-Montalcini, 1 - Area MIND, Milano, Lombardy 20157, Italy

^20^Department of Medicine, Faculty of Medicine Siriraj Hospital, Mahidol University, Bangkok 10700, Thailand

^21^Laboratory for Autoimmune Diseases, RIKEN Center for Integrative Medical Sciences, 1-7-22 Suehiro-cho, Tsurumi-ku, Yokohama, Kanagawa 230-0045, Japan

^22^Biotechnology Research and Innovation Council - National Institute of Biomedical Genomics, Kalyani, West Bengal 741251, India

^23^Department of Biomedical Sciences, Seoul National University College of Medicine, Seoul 03080, Republic of Korea

^24^Graduate School of Medical Science and Engineering, KAIST, Daejeon 34051, Republic of Korea

^25^Laboratory for Human Immunogenetics, RIKEN Center for Integrative Medical Sciences, 1-7-22 Suehiro-cho, Tsurumi-ku, Yokohama, Kanagawa 230-0045, Japan

^26^Department of Allergy and Rheumatology, Graduate School of Medicine, The University of Tokyo, 7-3-1 Hongo, Bunkyo-ku, Tokyo 113-8654, Japan

^27^Laboratory for Systems Genetics, RIKEN Center for Integrative Medical Sciences, 1-7-22 Suehiro-cho, Tsurumi-ku, Yokohama, Kanagawa 230-0045, Japan

^28^Department of Statistical Genetics, Graduate School of Medicine, Osaka University, 2-2 Yamadaoka, Suita, Osaka 565-0871, Japan

^29^Integrated Frontier Research for Medical Science Division, Institute for Open and Transdisciplinary Research Initiatives, Osaka University, 2-2 Yamadaoka, Suita, Osaka 565-0871, Japan

^30^Department of Genome Informatics, Graduate School of Medicine, The University of Tokyo, 7-3-1 Hongo, Bunkyo-ku, Tokyo 113-8654, Japan

^31^Laboratory of Statistical Immunology, Immunology Frontier Research Center (WPI-IFReC), Osaka University, 3-1 Yamadaoka, Suita, Osaka 565-0871, Japan

^32^Premium Research Institute for Human Metaverse Medicine (WPI-PRIMe), Osaka University, Suita 565-0871, Japan

## Author contributions

F.J.T., S.A.T., M.D.L., S.P., A.C.V., and G.R. conceived, and led the overall study design; A.M.C., S.A.F., W.S., M.F.M., and K.H.K. led the data integration effort; M.F.M. and M.D.L. developed and implemented data integration metrics with support from A.M.C., S.A.F., W.S., K.H.K., S.P., A.C.V., and G.R.; A.A. and G.R. developed scTiger with support from S.A.F. and P.R. who tested the implementation; A.M.C., K.H.K., J.N., R.S., E.V.B, L.K., R.G.H.L., L.B., A.C.V., and G.R. co-led the HARP cell annotation; J.N., S.A.F, and M.F. transferred final cell annotations and associated rationales on the HCA Cell Annotation Platform (CAP); A.M.C., S.A.F., W.S., and K.H.K. co-led downstream HARP analyses; J.N., L.F., O.D., M.A., S.M, R.G.H.L., R.N., H.H. contributed to downstream HARP analysis strategy, execution, and data interpretation; K.S. developed data portal; A.M.C., M.F.M., S.A.F., K.K., M.F., I.Z., A.C., and P.N. contributed to gathering Tier 1 meta-data, Tier 2 meta-data, gene expression matrices, and cell annotation meta-data; K.S., P.S., C.V.C., MGH COVID-19 Collection & Processing Team, and A.C.V. contributed the Slowikowski et al. COVID19 dataset before publication; K.H.K., Y.A., A.C., J.E.P., P.P.M., P.M., V.C., J.W.S., W.Y.P., Asian Immune Diversity Atlas (AIDA) Network, and S.P. contributed the AIDA dataset before publication; B.B. and J.P. contributed the unpublished CITE-Seq *HCAunpublished1* dataset; A.M.C., S.A.F., M.F.M., K.S., A.P., L.J., B.B., A.Y., S.S., and B.G.G. contributed in remapping datasets to be integrated; M.D.L., S.P., A.C.V., and G.R. managed and supervised the entire study; A.M.C., S.A.F., A.A., W.S., M.F.M., K.H.K., J.N., M.D.L., S.P., A.C.V., and G.R. wrote the manuscript with input from all authors.

