## Supplementary Figures for "A Community-Driven Single-Cell PBMC Reference Integrating Landmark Datasets Spanning Health and Disease"

#### ***Extended figures***

Extended Fig 1

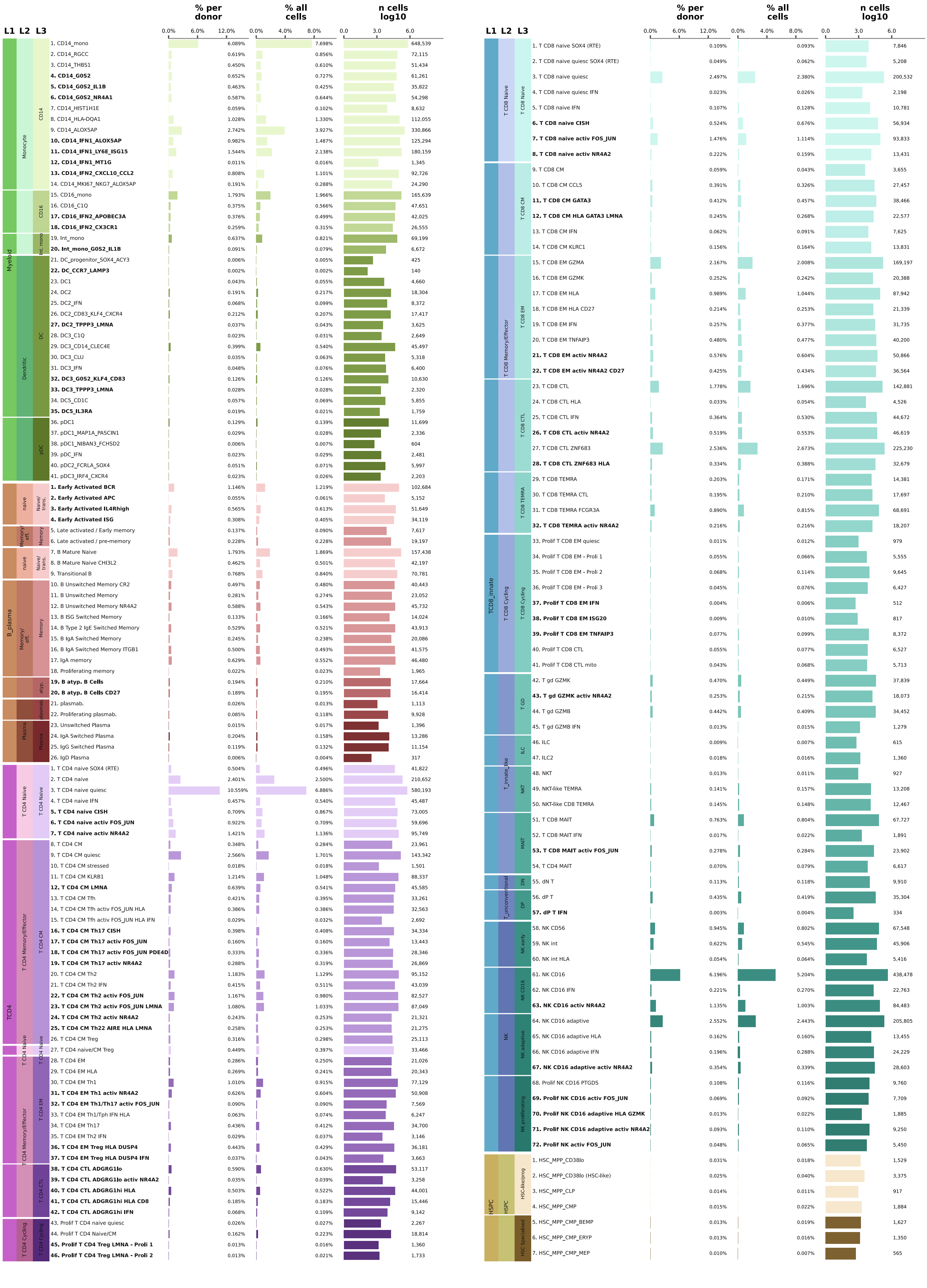

**Extended Figure 1. Summary of hierarchical organisation of annotated cell subsets and cell subset abundance (related to Main Figure 3).** Cell subsets are grouped by Level 1, Level 2, and Level 3 annotations, with Level 4 cell subsets shown as row labels. Colors indicate lineage/sublineage groupings within the annotation hierarchy. Bold Level 4 labels denote cell subsets we identified as novel/rare populations. Barplots show (from left to right) the average frequency per donor, the frequency across all cells, and the total number of cells per Level 4 cell subset represented as log10-transformed cell counts. Cell subsets are ordered within each Level 1 annotation according to their numbered Level 4 annotation in Figure 3A. Abbreviations: Int. mono = intermediate monocytes, plasmab. = plasmablasts; atyp. = atypical B cells; eff. = effector; trans. = transitional, HSC = Hematopoietic Stem Cell, MPP = Multipotent Progenitor, CLP = Common Lymphoid Progenitor, CMP = Common Myeloid Progenitor, BEMP = Basophil/Eosinophcil/Mast Cell Progenitor, ERYP = Erythroid Progenitor, MEP = Megakaryocyte–Erythroid Progenitor.

#### ***Supplementary figures***

Supplementary Fig. 1

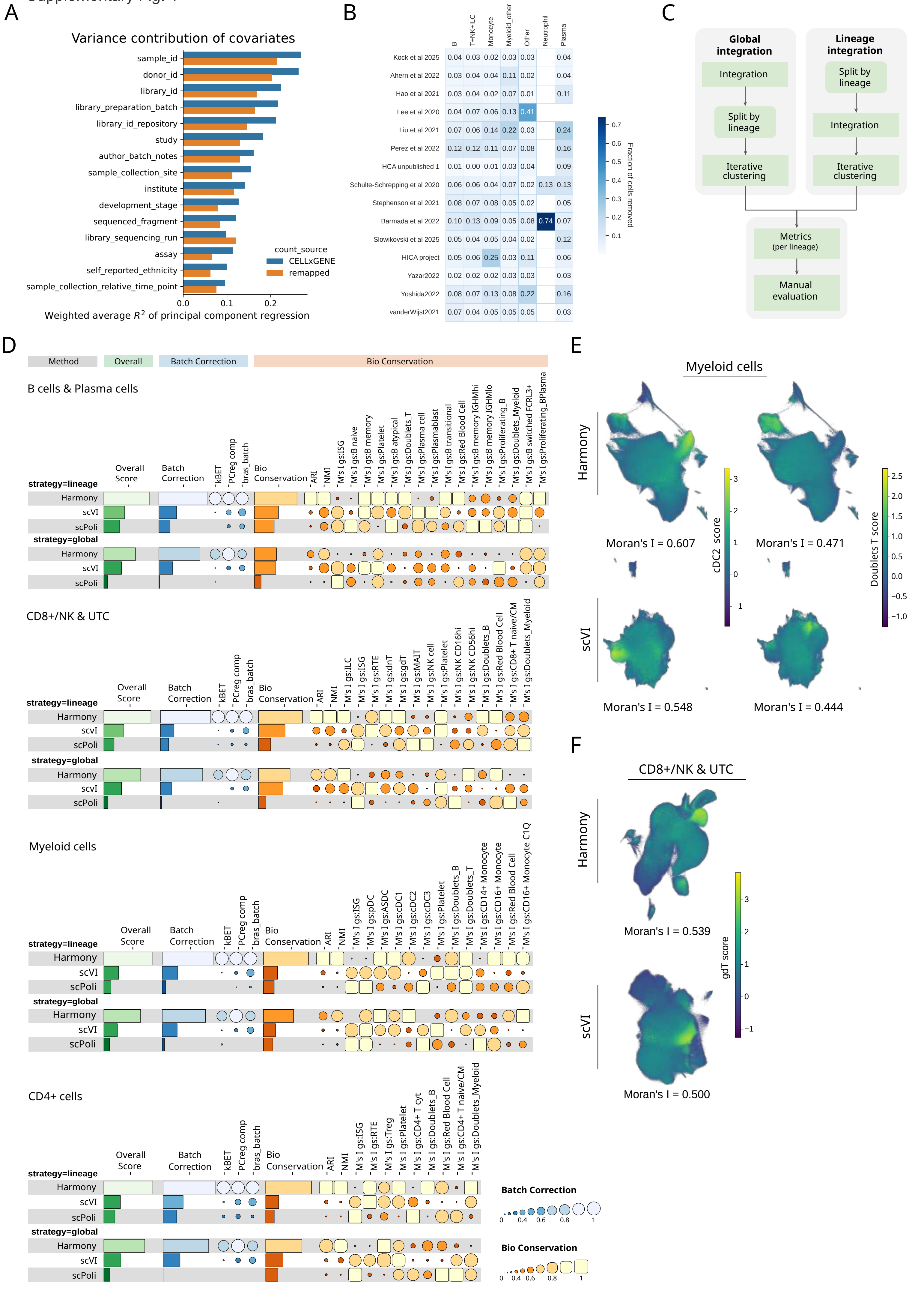

**Supplementary Figure 1. Assessment of data integration strategies (related to Main Figure 2). (A)** Linear variance contributions of technical covariates towards merged count matrix from study-provided count matrices published on CZ CELLxGENE[82] compared to merged count matrix from remapped studies. **(B)** Overview of fraction of cells removed per study and per cell lineage using semi-automated QC approach. **(C)** Schema of global vs lineage-specific integration and evaluation workflow as implemented with scAtlasTb. **(D)** Benchmark of lineage-specific and global integrations across different integration methods evaluated per lineage. **(E,F)** UMAP of additional Moran's I examples for **(E)** cDC2, T cell doublets, and **(F)** γδ T cells highlighting differences in biologically relevant gene set score correlation with kNN graph between Harmony (top) and scVI (bottom) integrations.

Supplementary Fig. 2

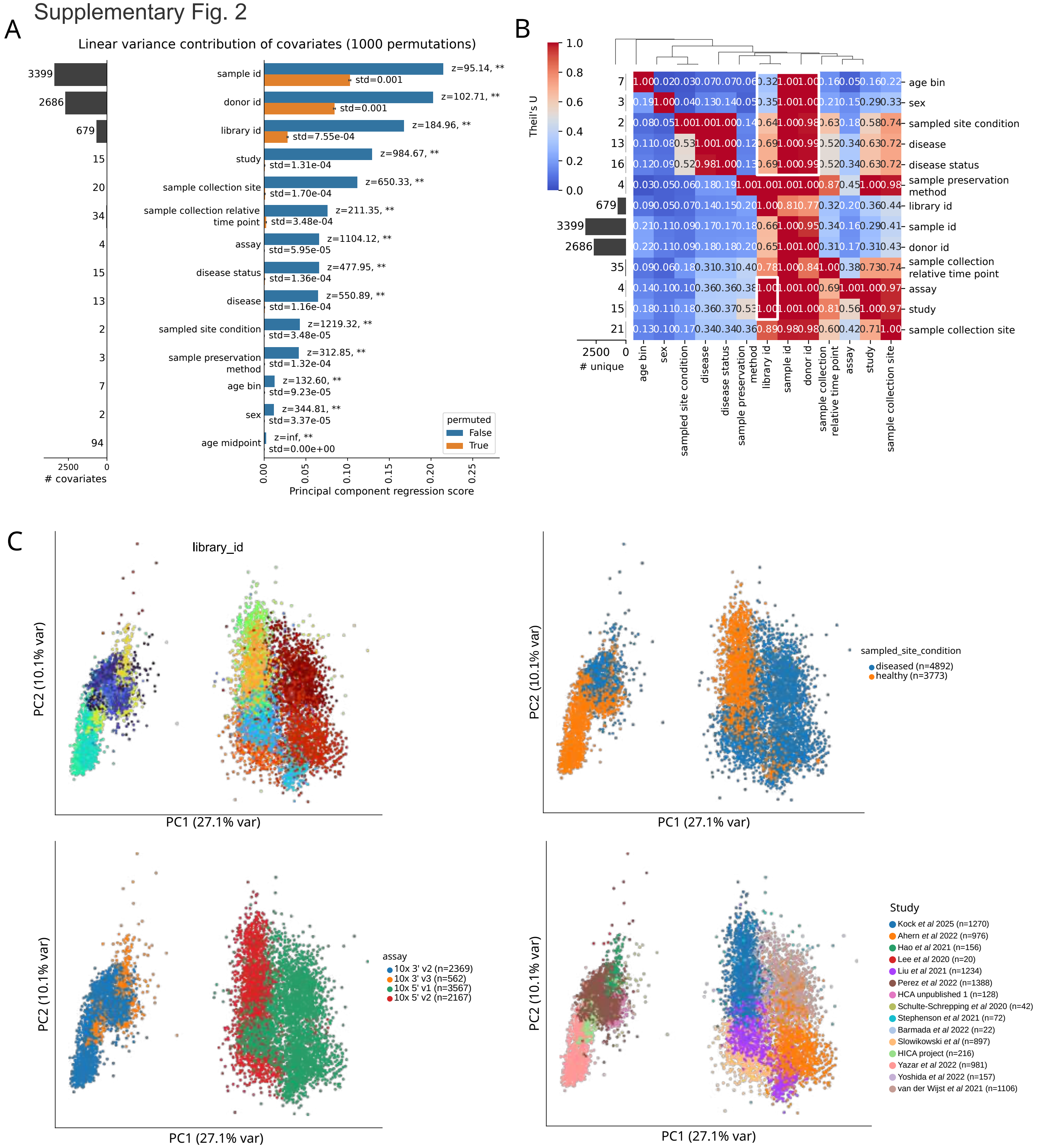

**Supplementary Figure 2. Batch analysis for determining optimal batch variable (related to Main Figure 2). (A)** Principal component regression analysis for technical and biological covariates. Significance is calculated based on a permuted background distribution per covariate (Methods). **(B)** Confounding analysis of technical and biological covariates using Theil's U. Lower values indicate less confounding between two covariates; Theil's U is non-symmetric; given one variable, it represents how much information it explains in another. **(C)** PCA plots of pseudobulk profiles (per smallest entity of a batch; Methods) for full dataset, colored by library ID, sampled site condition, assay, and study.

Supplementary Fig. 3

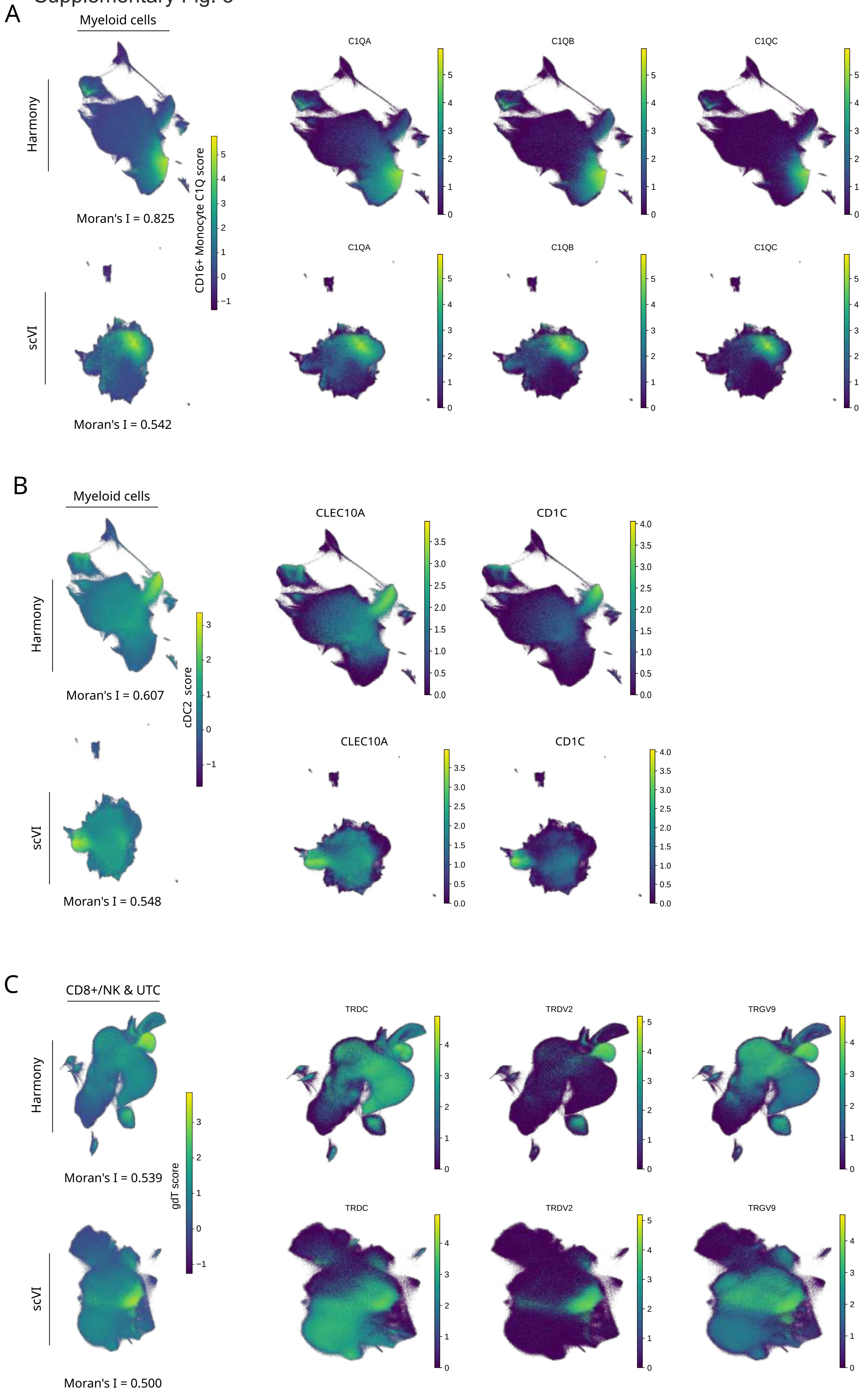

**Supplementary Figure 3. UMAPs with gene set scores and marker genes (related to Main Figure 2). (A)** Gene set score and gene expression for CD16 C1Q$^{Hi}$ monocytes in myeloid lineage. **(B)** Gene set score and gene expression for cDC2 cells in myeloid lineage. **(C)** Gene set score and gene expression for γδ T cells in CD8 T, NK, and unconventional T cell (UTC) population.

Supplementary Fig. 4

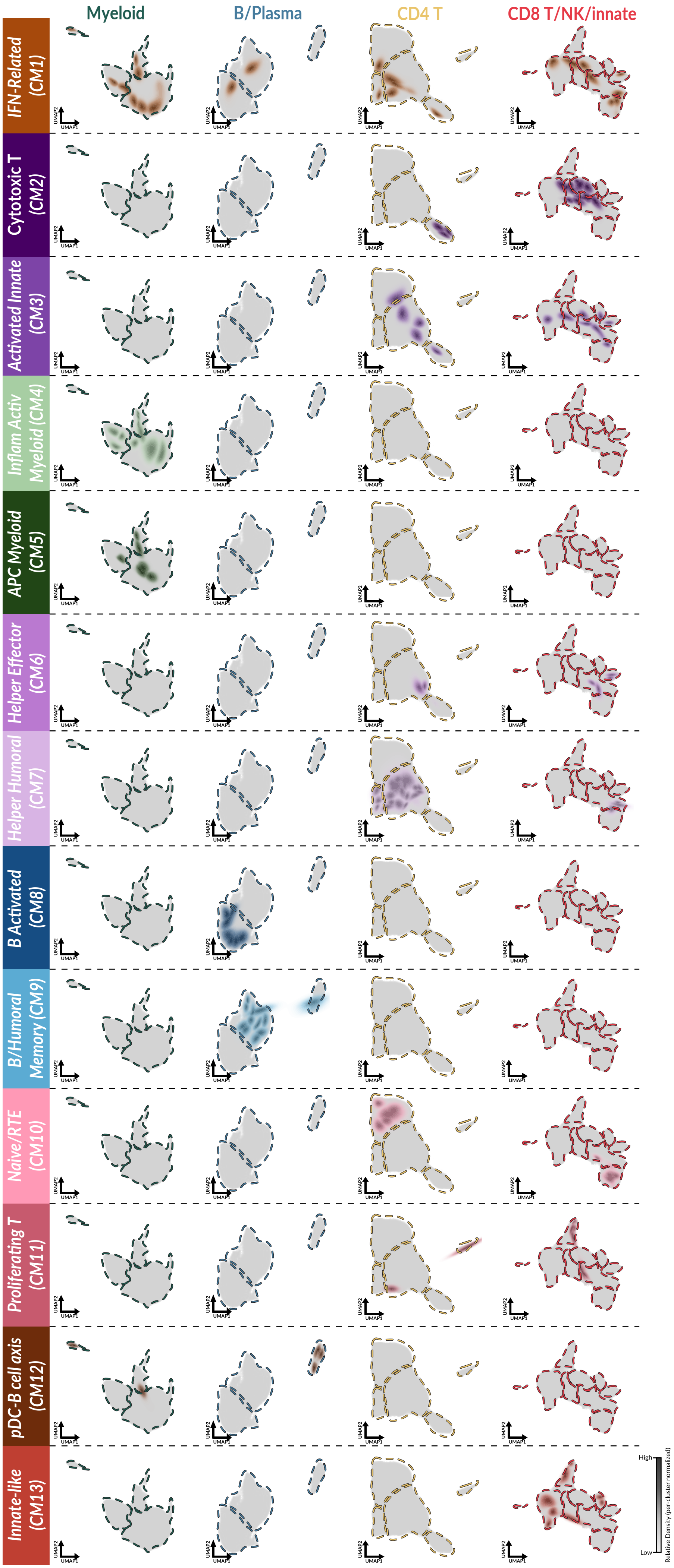

**Supplementary Figure 4 (Related to Main Figure 3). Contributions of different lineages to co-abundance Cellular Modules (CMs).** (**A**) Density plot UMAPs of all 13 CMs (extension of Main Fig 3E), illustrating the varying degrees of multi-lineage cell subset composition across CMs. Density scales are normalized per cell type to emphasize localized peaks across low- and high-abundance populations.

### Supplementary Fig. 5

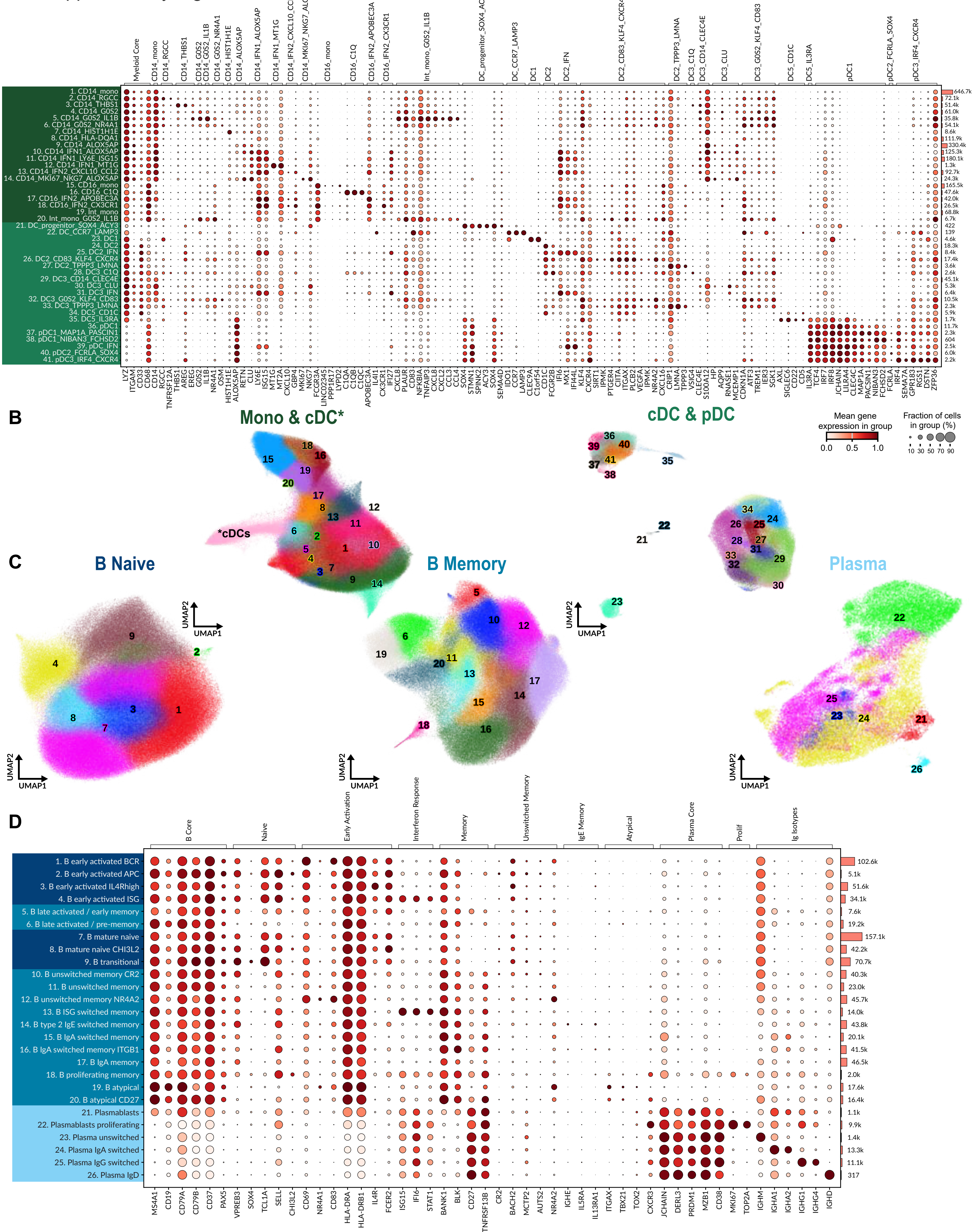

**Supplementary Figure 5 (Related to Main Figure 3). Annotation evidence for monocyte, dendritic cell, B, and plasma cell subsets.** (**A**) Dotplot of marker gene expression, annotated by cell numbers for each of the monocyte and dendritic cell subsets (horizontal bars). The background shading of the cell subset labels on the y-axis is color-coded to match the colors of their respective sublineage titles in (B). (**B**) UMAP embeddings of monocyte and dendritic cell populations from Mono/cDC and cDC/pDC sublineages. *Note that cDCs were included in the Mono/cDC embedding for additional context to annotate Monocyte subpopulations; however, cDC cells were ultimately annotated using the cDC/pDC embedding. (**C**) UMAP embeddings of B/Plasma cell populations from B Naive, B Memory, and Plasma sublineages. (**D**) Dotplot of marker gene expression, annotated by cell numbers for each of the B/Plasma cell subsets (horizontal bars). The background shading of the cell-type labels on the y-axis is color-coded to match their respective sublineage title color in (**C**). Numerical identifiers for all cell subsets correspond directly to those in the full lineage UMAPs in Figure 3A.

Supplementary Fig. 6

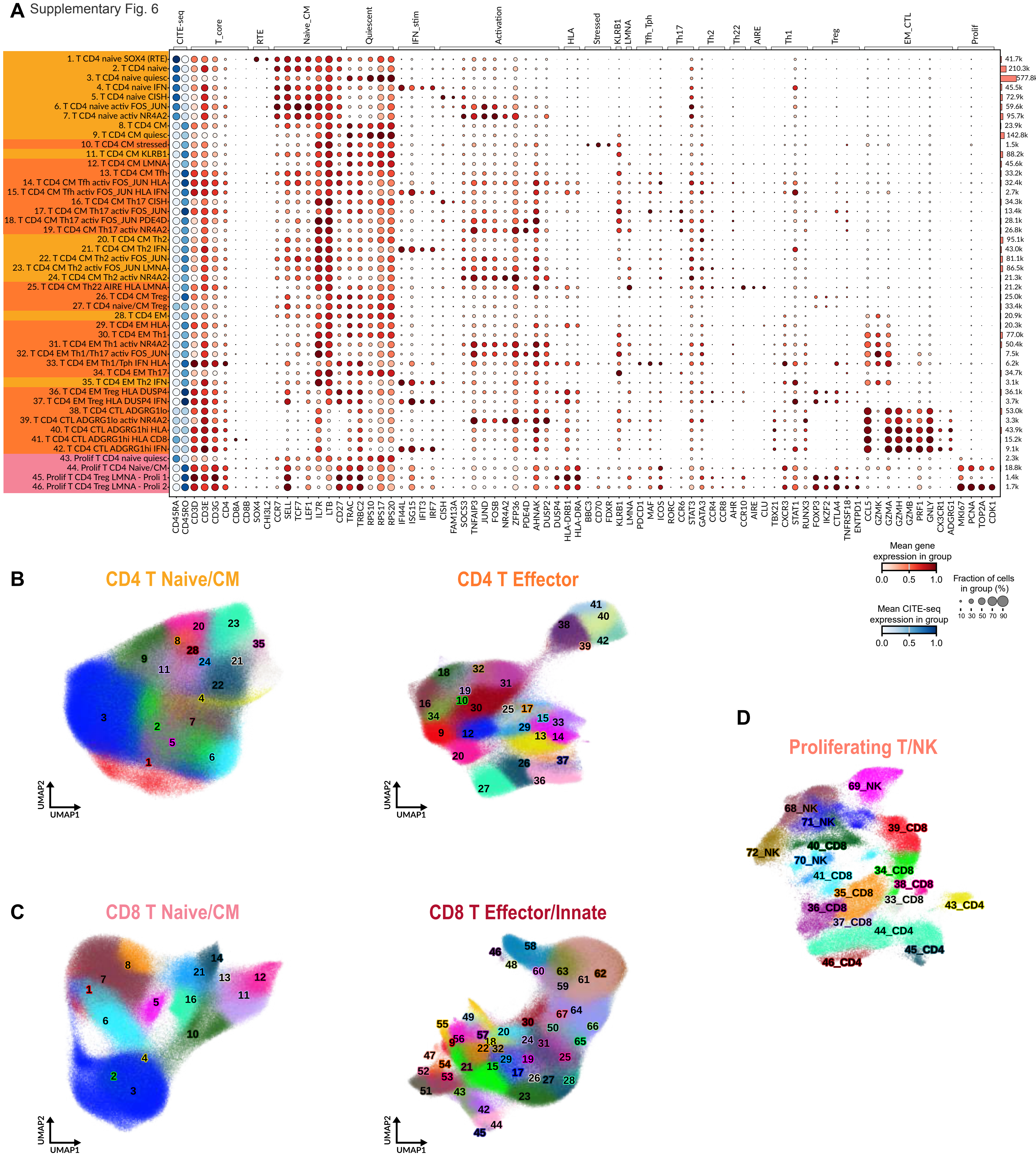

**Supplementary Figure 6 (Related to Main Figure 3). Annotation evidence for CD4 and CD8 T cell subsets.** (**A**) Dotplot of marker gene expression, annotated by cell numbers for each of the CD4 T cell subsets (horizontal bars). The background shading of the cell-type labels on the y-axis is color-coded to match their respective sublineage title color in (**B**). (**B**) UMAP embeddings of CD4 T cell populations from T Naive/Central Memory and CD4 T Effector sublineages. (**C**) UMAP embeddings of CD8 T/innate populations from CD8 T Naive/Central Memory and CD8 T Effector/innate lymphoid cells ("Innate") sublineages. (**D**) UMAP embedding of Proliferating T and NK cells which span CD4 T, CD8 T, and NK lineages.

Supplementary Fig. 7

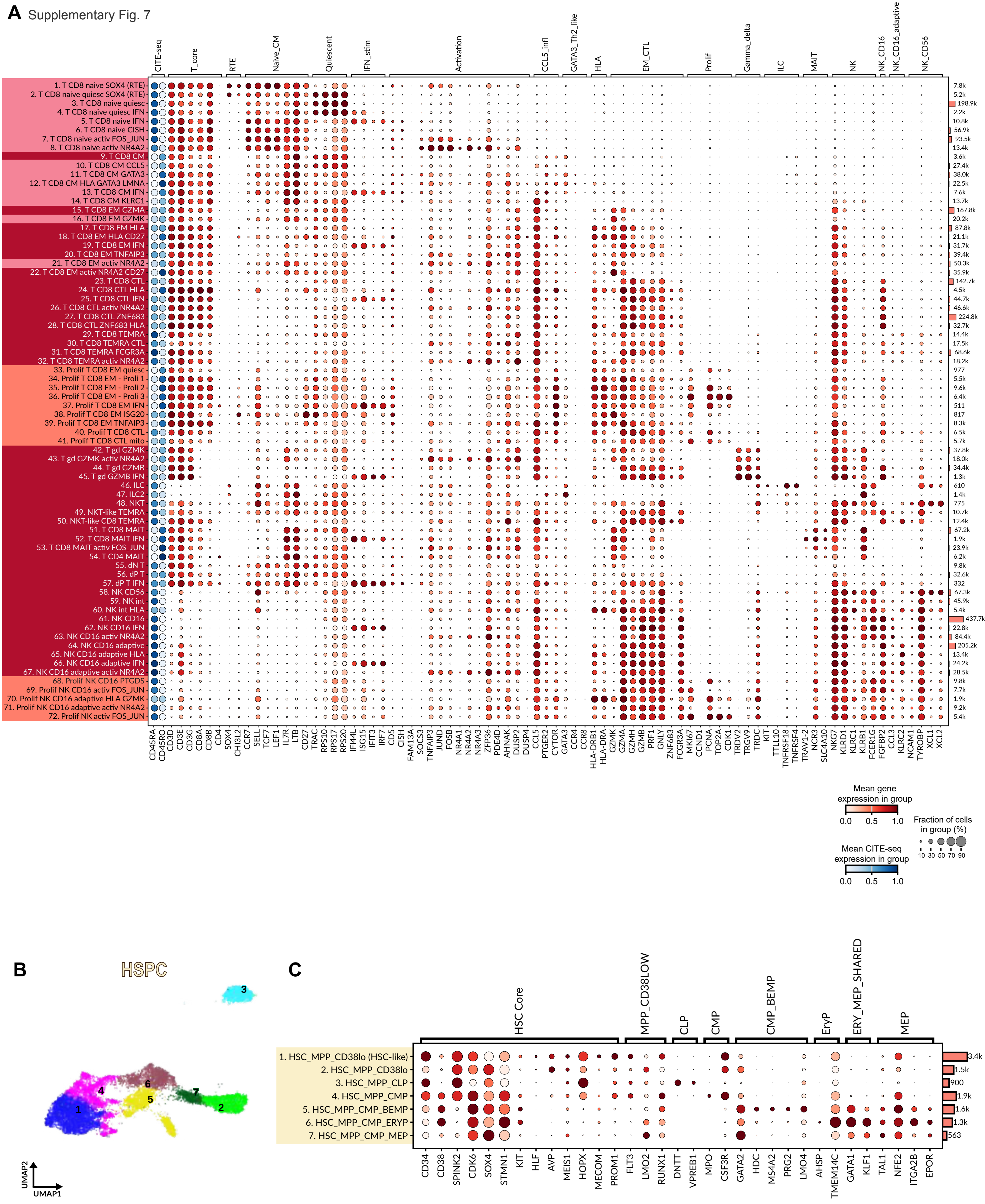

**Supplementary Figure 7 (Related to Main Figure 3). Annotation evidence for CD8 T, gdT, innate lymphoid cell, and hematopoietic stem and progenitor cell (HSPC) subsets.** (**A**) Dotplot of marker gene expression, annotated by cell numbers for each of the CD8 T, gdT, and innate lymphoid cell (including NK cell) subsets (horizontal bars). The background shading of the cell-type labels on the y-axis is color-coded to match their respective sublineage title color in Supplementary Fig. 6C. Numerical identifiers for all cell subsets correspond directly to those in the full lineage UMAP in Figure 3A. (**B**) UMAP embeddings of HSPC cell subsets (**C**) Dotplots of marker gene expression for the HSPC cell subtypes. HSC = Hematopoietic Stem Cell, MPP = Multipotent Progenitor, CLP = Common Lymphoid Progenitor, CMP = Common Myeloid Progenitor, BEMP = Basophil/Eosinophil/Mast Cell Progenitor, ERYP = Erythroid Progenitor, MEP = Megakaryocyte–Erythroid Progenitor.

Supplementary Fig. 8

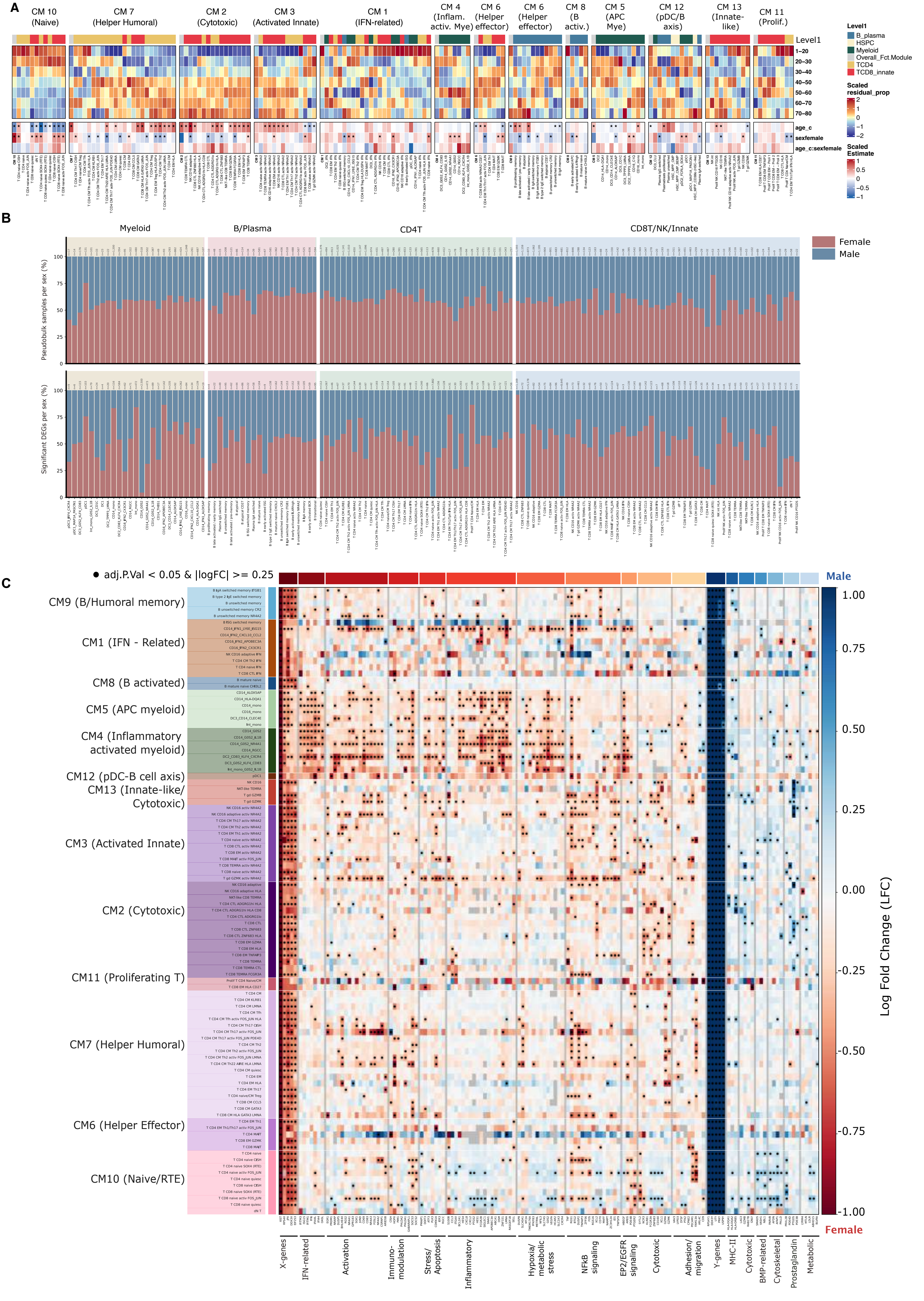

**Supplementary Figure 8. Age and sex-associated variation in immune cell-subset abundance and gene program activity (related to Main Figure 4).** (**A**) Summary heatmap showing age-associated cell subset abundance trajectories and linear mixed-effects model (LMM) coefficients across cellular modules (CMs). Upper panel shows study-corrected cell-type proportions averaged across age bins and z-scored per cell subset to highlight relative temporal dynamics. The lower panel shows scaled LMM coefficients for age, sex, and age-by-sex interaction terms. Cell subsets are annotated by Level 1 annotation and grouped by CMs. Asterisks denote BH-adjusted $p < 0.05$. (**B**) Barplots of 122 cell subsets (Level 4 annotation) used for DEG analysis. Upper panel shows the sex composition of pseudobulk samples (red = female, blue = male; n = total samples per cell subset). Lower panel shows the direction of significant sex-associated DEGs (BH-adjusted $p < 0.05$, $|logFC| \geq 0.25$; red = female-higher, blue = male-higher; n = total significant DEGs per cell subset). (**C**) Heatmap of log fold change (LFC) across all 98 cell subsets belonging to CMs, prior to filtering for cell subsets with ≥10 DEGs and a consistent sex direction, colored as in Fig. 4E (red = female-high, blue = male-high expression). Filled circles denote BH-adjusted $p < 0.05$ and $|LFC| \geq 0.25$; grey cells indicate cell subsets not tested due to power limitations.

Supplementary Fig.9

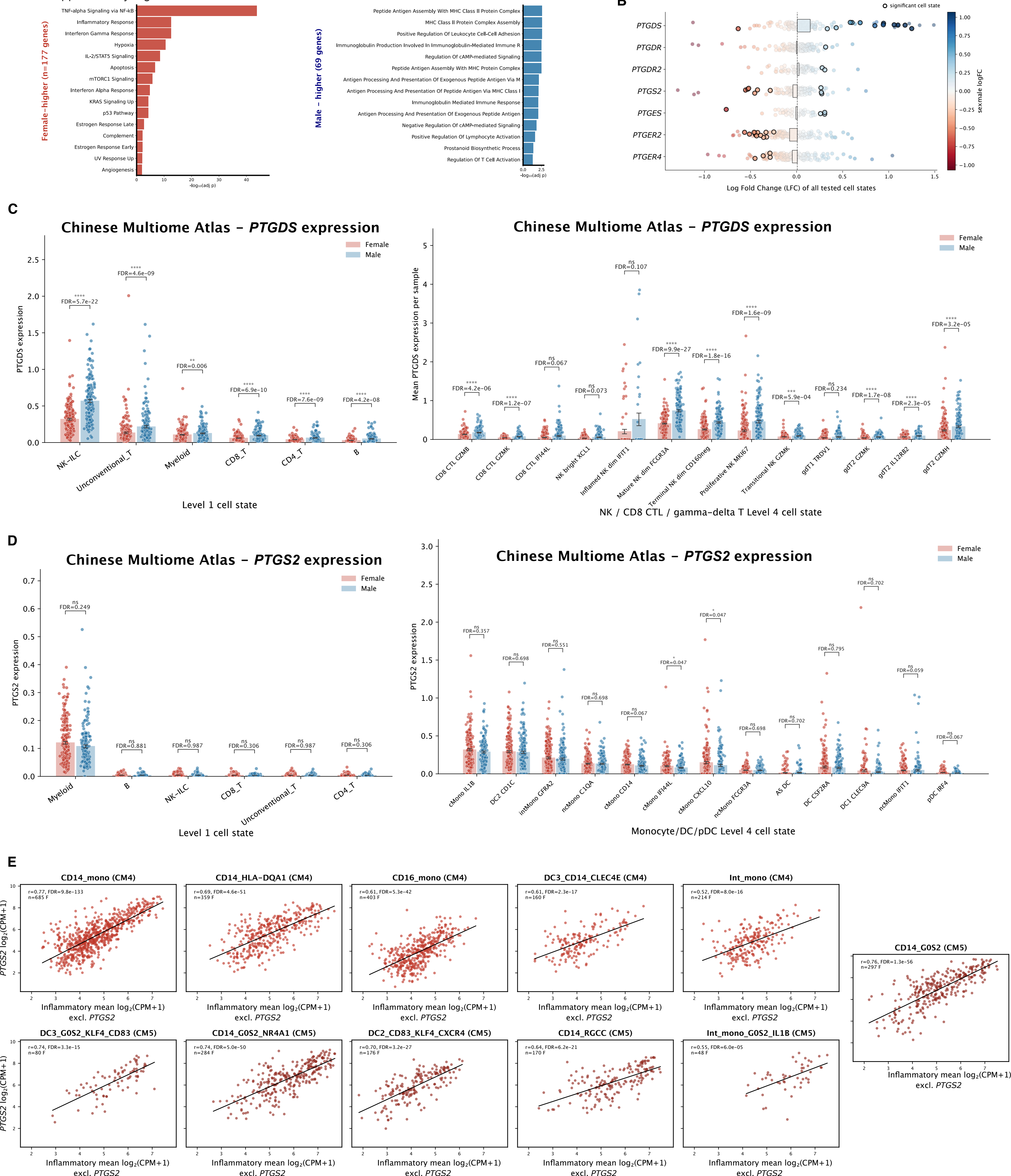

**Supplementary Figure 9. Sex-associated variation in cell type-specific gene expression profiles (related to Main Figure 4). (A**) Pathway enrichment analysis of recurrent sex-DEGs, for females (left) and males (right). *Left:* Over-representation analysis (Enrichr, MSigDB Hallmark 2020) of the 177 female-higher genes from recurrent sex-DEG set (adj. $p < 0.05$, $|logFC| > 0.3$, recurrent in ≥4 cell subsets). Bars show the top 15 significantly enriched terms (adj. $p < 0.05$), ranked by $-\log_{10}$(adjusted p-value). *Right:* Over-representation analysis (Enrichr, GO Biological Process 2023) of the 69 male-higher genes from the same recurrent sex-DEG set. Bars show the top 14 significantly enriched terms (adj. $p < 0.05$), ranked by $-\log_{10}$(adjusted p-value). **(B)** Beeswarm plot showing average log fold change (LFC) of selected prostaglandin pathway genes (*PTGDS, PTGDR, PTGDR2, PTGS2, PTGES, PTGER2, PTGER4*) across cell subsets. Each circle represents one cell subset; circles with bolded circumferences indicate cell subsets with statistical significance. Circle colour reflects the direction and magnitude of the LFC (red = female-high, blue = male-high). **(C)** Barplots of mean log2p(CPM + 1) *PTGDS* expression per sample in the Chinese Multiome Atlas[49], shown across Level 1 cell annotations (left) and Level 4 NK/CD8 CTL/gamma-delta T cell subsets (right). Each dot represents one sample, coloured by sex (red = female, blue = male). Coloured bars indicate group means. BH-corrected p-values from Mann-Whitney U tests are shown above brackets. **(D)** Barplots of mean *PTGS2* expression per sample in the Chinese Multiome Atlas, shown across Level 1 annotations (left) and Level 4 monocyte/DC/pDC cell subsets (right). Each dot represents one sample, coloured by sex (red = female, blue = male). Coloured bars indicate group means. BH-corrected p-values from Mann-Whitney U tests are shown above brackets. (**E**) Scatter plots showing the correlation between *PTGS2* expression and innate inflammatory gene program (excluding *PTGS2*) activity across myeloid cell subsets in female samples. Each panel corresponds to one Level 4 cell subset (minimum 10 female samples), labelled by CM (CM1, CM4, or CM5) and cell subset. Each point represents one female sample. Black lines show linear fits. Panel labels report Pearson *r*, BH-adjusted *p*-value across the 25 tested cell subsets, and sample size.

Supplementary Fig. 10

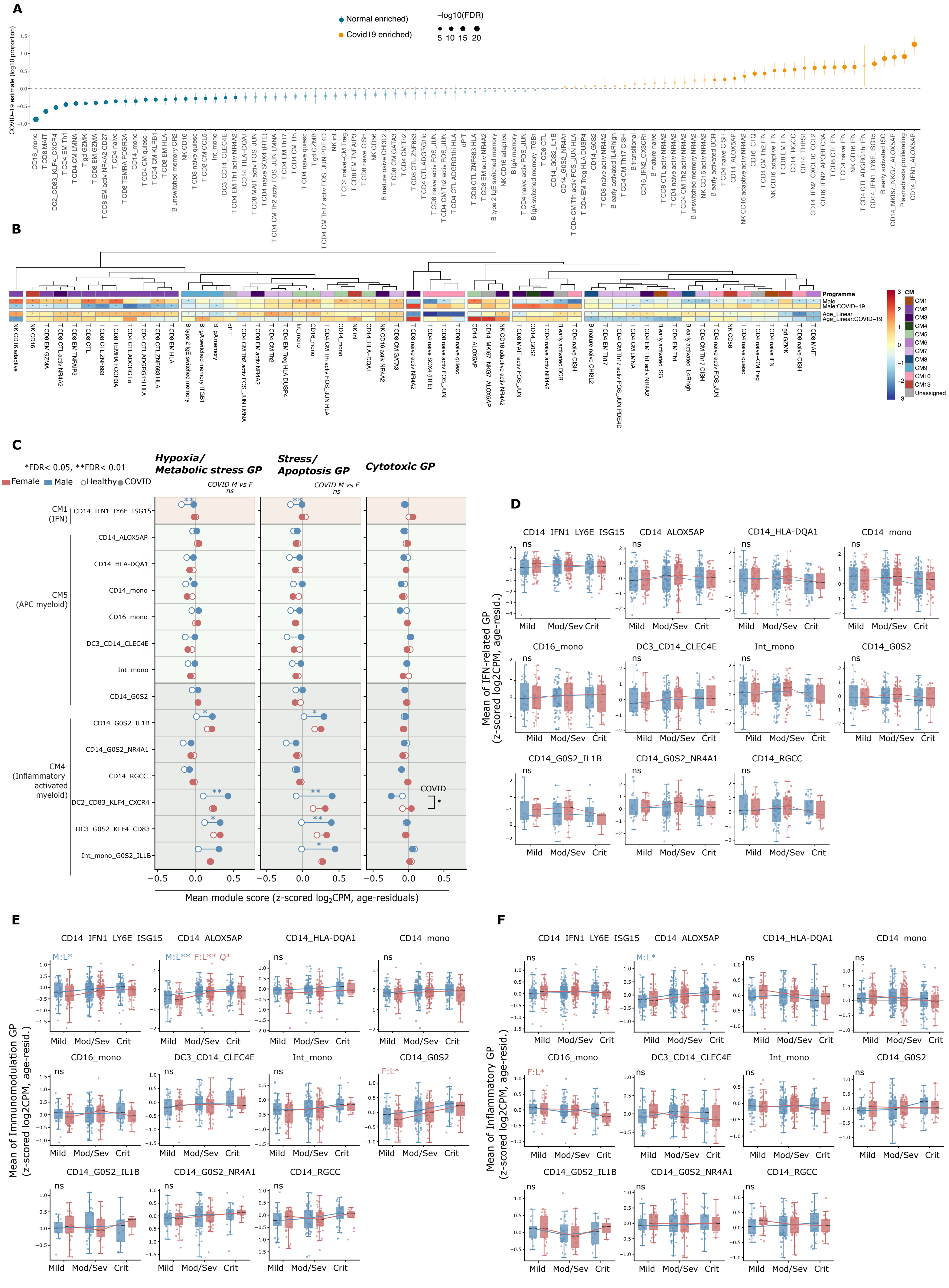

**Supplementary Figure 10. Disease, age, and sex-associated variation in immune cell-subset abundance and gene program activity during COVID-19 (related to Main Figure 5).** (**A**) Lollipop plot depicting per-cell-subset coefficients of the COVID-19 disease versus controls covariate ($\log_{10}$(proportion) ~ age_bin + sex + COVID-19_or_control + study). (**B**) Heatmap of Z-scores of coefficients of the male sex covariate; the interaction term between male sex and moderate, severe, or critical COVID-19 status; the age bin covariate; and the interaction term between age bin and moderate, severe, or critical COVID-19 status ($\log_{10}$(proportion) ~ age_bin + sex + moderate_severe_critical_COVID-19_or_control + sex:moderate_severe_critical_COVID-19_or_control + age_bin:moderate_severe_critical_COVID-19_or_control + study). Z-scores were computed individually for each covariate across all cell subsets in the heatmap, and hierarchical clustering was performed to order the cell subsets. Asterisks indicate FDR < 0.05 for the corresponding coefficient-cell subset combination. Cell subsets are annotated by their CM membership. Proportions for each cell subset per donor were computed out of all PBMCs per donor. **(C)** Lollipop plots for three myeloid GPs not shown in Fig. 5C - Hypoxia/Metabolic Stress, Stress/Apoptosis, and Cytotoxic - across 14 myeloid cell subsets. Score computation, OLS contrasts, BH correction of p-values, and display conventions match Fig. 5H. **(D)** Severity-dependent IFN-related GP scores across all 14 myeloid cell subsets, complementary to Fig. 5I. Statistical framework, boxplot display, and annotations match Fig. 5I. **(E)** As in (D), for the Immunomodulation GP. **(F)** As in (D), for the Inflammatory GP.

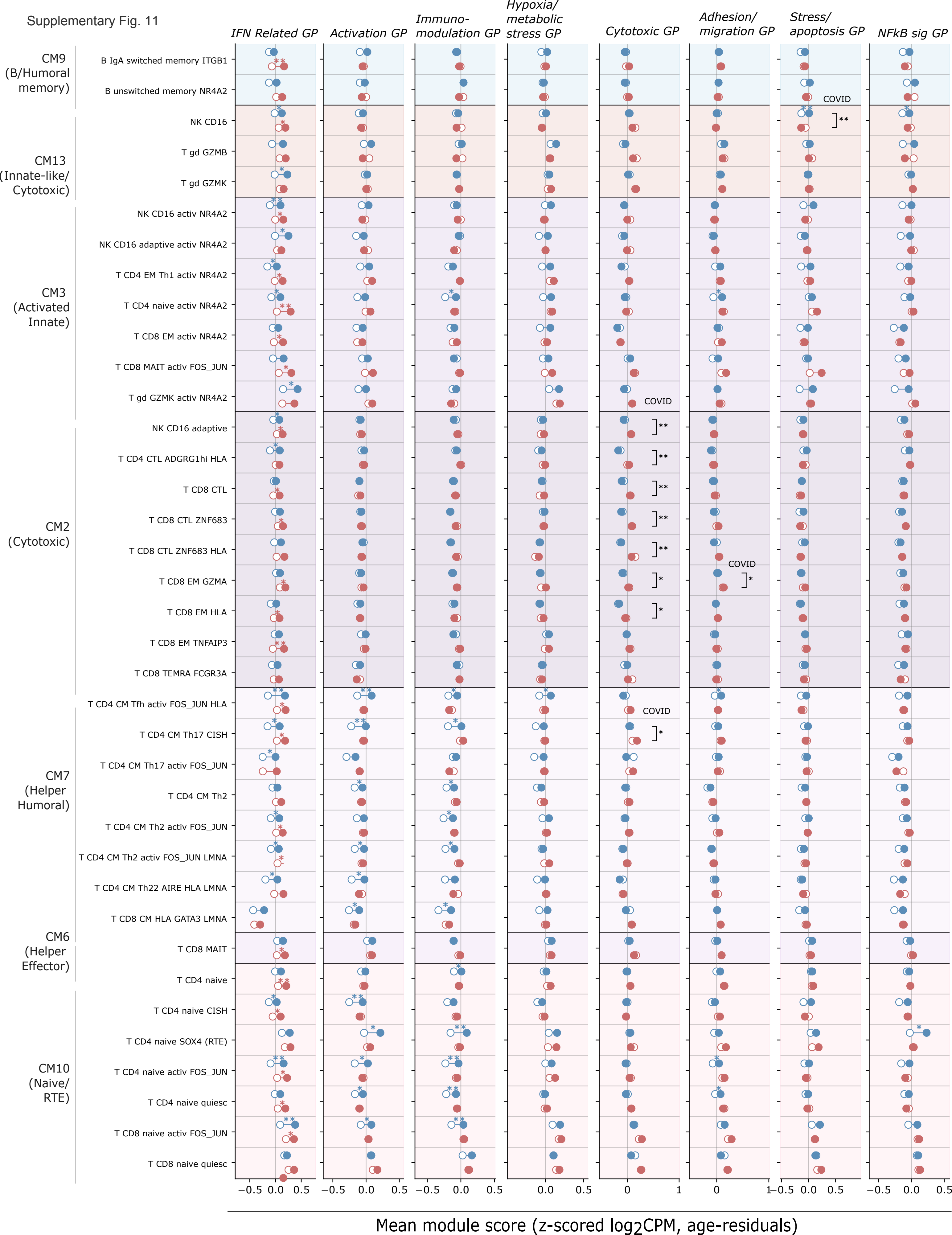

Supplementary Fig. 11
IFN Related GP
Activation GP
Immuno-modulation GP
Hypoxia/ metabolic stress GP
Cytotoxic GP
Adhesion/ migration GP
Stress/ apoptosis GP
NFkB sig GP
CM9 (B/Humoral memory)
B IgA switched memory ITGB1
B unswitched memory NR4A2
CM13 (Innate-like/ Cytotoxic)
NK CD16
T gd GZMB
T gd GZMK
CM3 (Activated Innate)
NK CD16 activ NR4A2
NK CD16 adaptive activ NR4A2
T CD4 EM Th1 activ NR4A2
T CD4 naive activ NR4A2
T CD8 EM activ NR4A2
T CD8 MAIT activ FOS_JUN
T gd GZMK activ NR4A2
CM2 (Cytotoxic)
NK CD16 adaptive
T CD4 CTL ADGRG1hi HLA
T CD8 CTL
T CD8 CTL ZNF683
T CD8 CTL ZNF683 HLA
T CD8 EM GZMA
T CD8 EM HLA
T CD8 EM TNFAIP3
T CD8 TEMRA FCGR3A
CM7 (Helper Humoral)
T CD4 CM Tfh activ FOS_JUN HLA
T CD4 CM Th17 CISH
T CD4 CM Th17 activ FOS_JUN
T CD4 CM Th2
T CD4 CM Th2 activ FOS_JUN
T CD4 CM Th2 activ FOS_JUN LMNA
T CD4 CM Th22 AIRE HLA LMNA
T CD8 CM HLA GATA3 LMNA
CM6 (Helper Effector)
T CD8 MAIT
CM10 (Naive/ RTE)
T CD4 naive
T CD4 naive CISH
T CD4 naive SOX4 (RTE)
T CD4 naive activ FOS_JUN
T CD4 naive quiesc
T CD8 naive activ FOS_JUN
T CD8 naive quiesc
COVID
−0.5 0.0 0.5
0 1
Mean module score (z-scored $log_2$CPM, age-residuals)

**Supplementary Figure 11. Disease and sex-associated variation in gene program activity during COVID-19 (related to Main Figure 5).** (**A**) Lollipop plots of mean GP scores for eight GPs - IFN-related, Activation, Immunomodulation, Hypoxia/Metabolic Stress, Cytotoxic, Adhesion/Migration, Stress/Apoptosis and NF-kB signaling - across 35 lymphoid cell subsets, grouped by CM (CM9: B/Humoral memory; CM13: Innate-like/Cytotoxic; CM3: Activated Innate; CM2: Cytotoxic; CM7: Helper Humoral; CM6: Helper Effector; CM10: Naive/RTE). GP scores are computed as in Fig. 5H (within-study z-scored $\log_2$(CPM +1) per gene, averaged across GP genes, OLS age residualisation), with the same three OLS contrasts per cell subset (female COVID vs. healthy; male COVID vs. healthy; female vs. male within COVID) and global BH correction of p-values per GP across lymphoid CMs × 3 contrasts. Visualization conventions and statistical significance thresholds as in Fig. 5H.

Supplementary Fig. 12

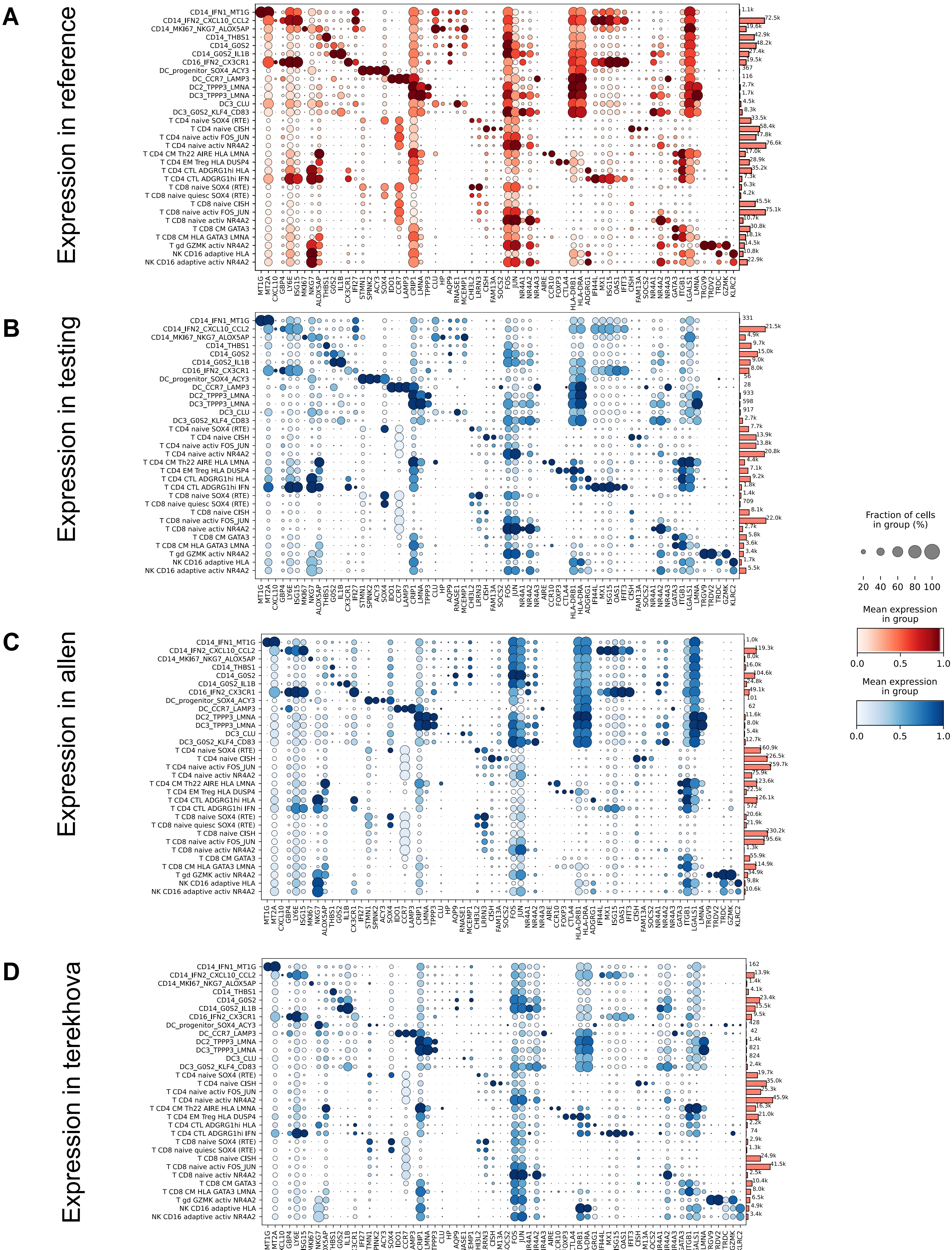

**Supplementary Figure 12. Validation of rare and high-resolution immune cell subsets across independent datasets (related to Main Figure 6).** Dotplots showing marker gene expression for selected rare and fine-grained immune populations identified by scTiger, annotated by number of cells per cell subset (horizontal bars). Cell subsets include activation-associated, interferon-responsive, and lineage-specialized lymphoid, myeloid and NK-cell populations. Dot size indicates the fraction of expressing cells and color denotes mean expression. **(A)** Marker gene expression in the HARP reference atlas used for model training. **(B)** Corresponding expression profiles in the held-out validation dataset comprising unseen batches and studies. **(C)** Expression profiles in the external Gong et al. dataset following reference mapping. **(D)** Expression profiles in the external Terekhova et al. dataset following reference mapping.

Supplementary Fig. 13

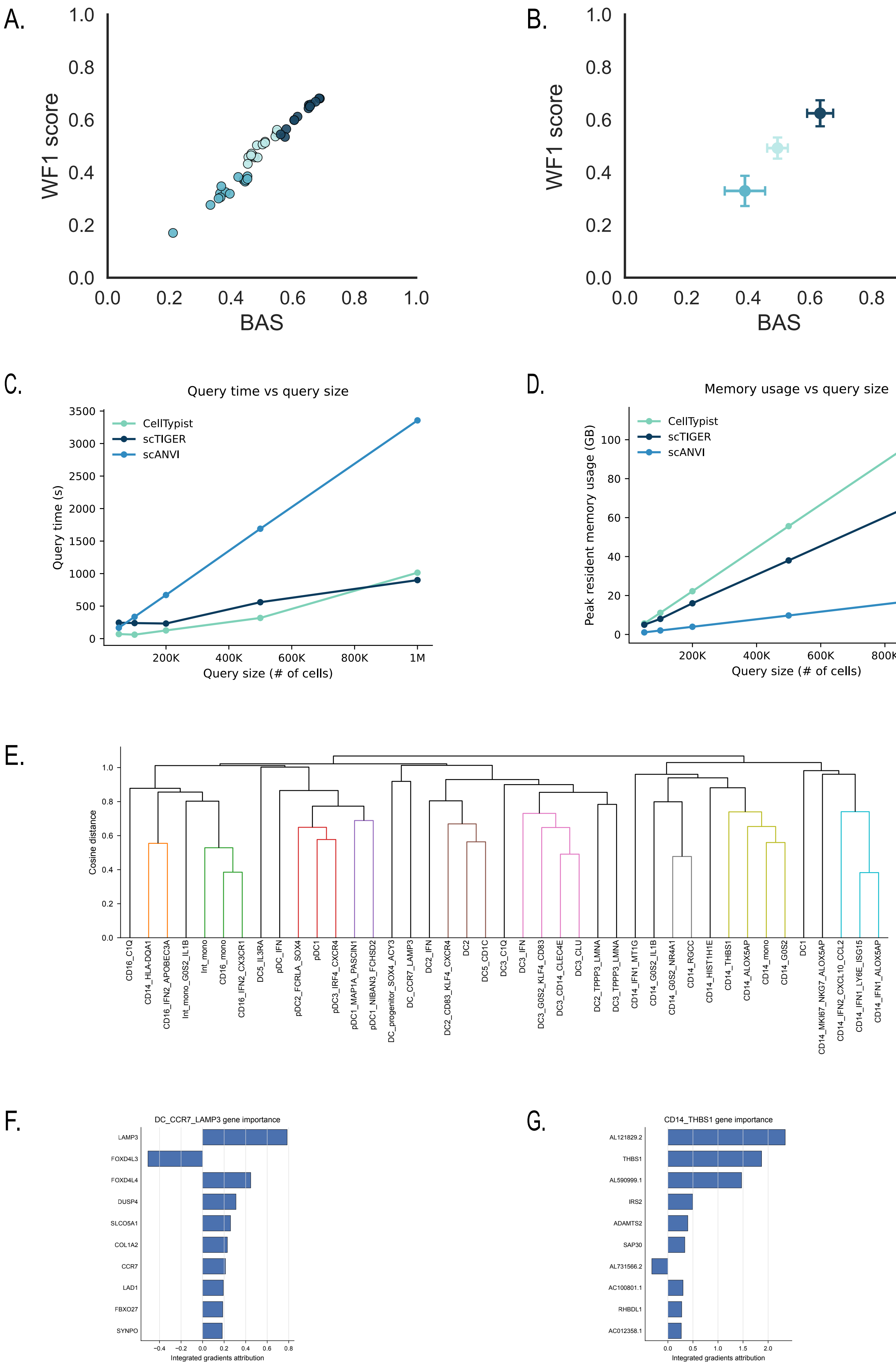

**Supplementary Figure 13. Additional benchmarking and interpretability analyses of scTiger (related to Main Figure 6). (A,B)** Additional performance evaluation of scTiger, CellTypist, and scANVI across held-out batches. **(A)** Relationship between balanced accuracy score (BAS) and weighted F1 score (WF1) across Level 2 populations. Each point represents one cell population. **(B)** Mean BAS and WF1 scores for each method; error bars indicate variability across populations. **(C,D)** Computational scalability benchmarking. **(C)** Query runtime as a function of dataset size (50,000–1,000,000 cells). **(D)** Peak resident memory usage as a function of dataset size. **(E–G)** Interpretability of lineage-specific scTiger models. **(E)** Hierarchical clustering of monocyte and DC population centroids in the learned latent space using cosine distance. **(F,G)** Integrated Gradients feature-attribution scores for the DC_CCR7_LAMP3 and CD14_THBS1 populations, respectively. Canonical marker genes, including *LAMP3*, *CCR7* and *THBS1*, are amongst the highest-contributing features, indicating that lineage-specific models learn biologically meaningful representations and classification programs.
